# Genome-wide chromosomal association of Upf1 is linked to Pol II transcription in *Schizosaccharomyces pombe*

**DOI:** 10.1101/2021.04.12.437332

**Authors:** Sandip De, Vibha Dwivedi, Jianming Wang, David M. Edwards, Wazeer Varsally, Anand K. Singh, Hannah L. Dixon, Marcin W. Wojewodzic, Francesco Falciani, Nikolas J. Hodges, Marco Saponaro, Kayoko Tanaka, Claus M. Azzalin, Peter Baumann, Daniel Hebenstreit, Saverio Brogna

## Abstract

Although the RNA helicase Upf1 has hitherto been examined mostly in relation to its cytoplasmic role in nonsense mediated mRNA decay (NMD), here we report high-throughput ChIP data indicating genome-wide association of Upf1 with active genes in *Schizosaccharomyces pombe*. This association is RNase sensitive and it correlates with Pol II transcription and mRNA expression levels. While changes in Pol II occupancy were detected at only some genes in a Upf1-deficient (*upf1Δ*) strain, there is an increased Ser2 Pol II signal at all highly transcribed genes examined by ChIP-qPCR. Furthermore, *upf1Δ* cells are hypersensitive to the transcription elongation inhibitor 6-azauracil and display Pol II abnormalities suggestive of Pol II hyperphosphorylation. A significant proportion of the genes associated with Upf1 in wild-type conditions are also mis-regulated in *upf1Δ.* These data envisage that by operating on the nascent transcript Upf1 might influence Pol II phosphorylation and transcription.

## Introduction

Upf1 is a conserved protein of eukaryotes that has so far been primarily studied for its key role in nonsense-mediated mRNA decay (NMD). NMD is a translation-coupled cytoplasmic mechanism believed to recognise and rapidly destroy mRNAs carrying a premature termination codon or other features that place a stop codon in abnormal sequence contexts (Brogna and Wen, 2009; Fatscher et al., 2015; Goetz and Wilkinson, 2017; He and Jacobson, 2015; Hug et al., 2016; Karousis et al., 2016; Kim and Maquat, 2019; Lykke-Andersen and Jensen, 2015). However, the precise role of Upf1 in NMD as well as the significance and NMD mechanisms remain unclear (Brogna et al., 2016; Neu-Yilik et al., 2017).

Upf1 belongs to the 1B superfamily (S1B) of helicases that are involved in a diverse range of cellular activities in all domains of life. These are characterised by conserved sequence motifs and the ability to translocate in a 5’ to 3’ direction on both RNA and DNA (Saikrishnan et al., 2009; Singleton et al., 2007). Specifically, there is evidence that Upf1 uses ATP hydrolysis to translocate on RNA and to displace RNA-bound proteins (Bhattacharya et al., 2000; Chakrabarti et al., 2011; Czaplinski et al., 1995; Fiorini et al., 2015; Franks et al., 2010).

In yeast as in other organisms, Upf1 is typically most abundant in the cytoplasm. For this reason, it is assumed that Upf1 operates on mRNAs only after their nuclear export. However, there is evidence that Upf1 traffics in and out of the nucleus in mammalian cells (Ajamian et al., 2015; Mendell et al., 2002). It was initially proposed that within the nucleus Upf1 plays a direct role in DNA replication, telomere maintenance and DNA repair (Azzalin and Lingner, 2006). However, the effects of Upf1 depletion on DNA replication and cell division might be an indirect consequence of changes in the expression of genes involved in these processes (Rehwinkel et al., 2005; Varsally and Brogna, 2012). There is circumstantial evidence that Upf1 might instead play a direct role in RNA-based processes of gene expression within the nucleus (Varsally and Brogna, 2012).

The putative association of Upf1 with chromatin was examined by chromatin immunoprecipitation (ChIP) in *Schizosaccharomyces pombe* with the aim to understand what roles Upf1 may have in the nucleus. The data demonstrate Upf1 binding to chromatin and indicate that this occurs primarily at active genes. This association is genome-wide and positively correlates with RNA Pol II loading and mRNA expression. Notably, this interaction is RNase sensitive and Upf1 does not co-purify with Pol II directly. These data therefore indicate that Upf1 binds the nascent transcript. Genes that are associated with Upf1 in wild-type conditions are more likely to be mis-regulated in *upf1*Δ cells. Upf1 depleted cells are also hypersensitive to the 6-azauracil, a drug that can affect transcription elongation, and present Pol II abnormalities suggestive of CTD hyperphosphorylation. Cumulatively these findings predict that Upf1, by operating on the nascent mRNA Pol II genes, can regulate both transcription and mRNA processing.

## Results

### Upf1 associates genome-wide with protein coding genes

It has been reported that the nuclear level of Upf1 increases upon incubation with leptomycin-B (LMB) in *S. pombe* (Orfeome localization data on Pombase). LMB specifically inhibits the CRM1-mediated protein and mRNP nuclear export pathway of eukaryotes (Fukuda et al., 1997; Hutten and Kehlenbach, 2007), therefore it is probable that Upf1 is also shuttling between the nucleus and cytoplasm in *S. pombe*. To explore what role it might play in the nucleus, we examined whether Upf1 is associated with individual genes by ChIP. Endogenous *Upf1* gene was tagged with the hemagglutinin (HA) epitope by homologous recombination and ChIP was performed using an HA antibody. This allowed for genome-wide enrichment profiles which were determined by hybridisation of the immunoprecipitated DNA to genomic tiling chip arrays (Affymetrix, see Materials and Methods). We examined both asynchronous and S phase synchronised cell cultures, in duplicate experiments. Significantly enriched regions were identified using the Model-based Analysis of Tiling Arrays (MAT) software (Material and Methods). A total of 594 and 696 genes that are significantly enriched by Upf1 ChIP in 50% or more of their sequence were identified, in asynchronous and S phase cultures respectively (Figure 1 shows the enrichment profiles over a representative region of chromosome 1; Table S1 gives the lists of the enriched genes in the two datasets). The Upf1 enrichment profiles are similar between the two samples, however there are several genes enriched more in the S phase sample, such as histone H2A beta (*hta2*, labelled in Figure 1A). Most of the enrichment regions correspond to protein coding genes, which is the class of genes we have investigated in further detail here, yet several non-coding RNA and tRNA genes also appear to be enriched (Figure 1E).

**Figure 1.**
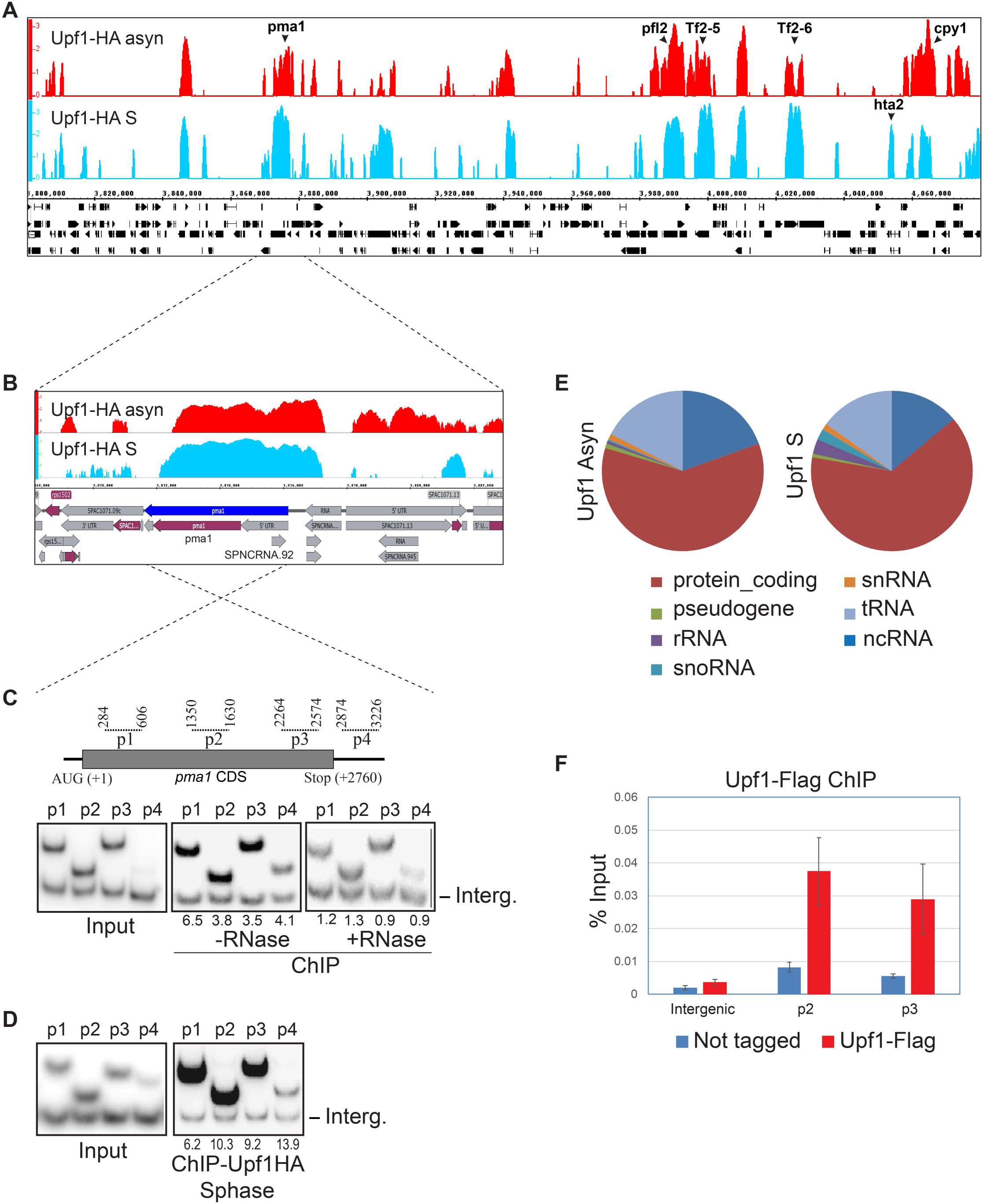
Genome-wide association of Upf1 with protein coding genes. **A)** IGB visualisation of Upf1-HA ChIP-chip enrichment in asynchronous culture (top track, shown in red) and in S-phase culture (bottom track, shown in sky blue) at a representative chromosomal region (270 kb) of *S. pombe* indicating enrichment of several specific genes, those discussed in the main text are labelled. Genes and genomic features are shown in black below. **B**) Zoomed-in view of Upf1 enrichment over the entire *pma1* gene (highlighted in dark blue) - 5’UTR (in grey), CDS (in purple) and 3’UTR (in grey) of *pma1* are shown in the bottom row schematic. **C**) Top panel-diagram of the *pma1* gene (cDNA region in grey) and the positions of amplicons used for the PCR-ChIP assay are indicated by the dotted lines above (numbers correspond to the position of the primers relative to start codon). Bottom panel-polyacrylamide gels showing radiolabelled PCR products produced by the *pma1*-specific primer pairs (top bands) and by the intergenic region specific primer (bottom bands, labelled Int.); using input DNA before ChIP (left panel) and using ChIP-enriched DNA from asynchronous culture without (middle panel) and with RNase pre-treatment of the chromatin (right panel). The relative enrichment of *pma1* DNA relative to intergenic sequence is expressed as a ratio of the intensity of the same fragments produced with the input DNA. **D**) PCR analysis as in **C** using input DNA before ChIP (left panel) and using ChIP-enriched DNA from S-phase culture without the RNase treatment (right panel). **E**) Pie-charts showing the proportion of different classes of gene associated to Upf1 in asynchronous and S-phase culture of *S. pombe.* **F**) ChIP-qPCR of Upf1-Flag enrichment over *pma1* in three independent asynchronous cultures.

The ChIP-chip data also show enrichment of repetitive sequences such as centromere and telomere regions, as well as Tf2 retrotransposons (Tf2-5 and Tf2-6 are indicated within the region shown in Figure 1A, and Tf2-9 is shown in Figure S1A). The seemingly high ChIP-chip enrichment of some of these repetitive regions may be a technical artefact of the hybridization due to their high sequence similarity. However, Tf2 enrichment was similarly detected in additional ChIP-qPCR experiments using a strain expressing Flag-tagged-Upf1, and was RNase-sensitive (Figure S1B). ChIP enrichment of some tRNA genes was also confirmed by qPCR in independent experiments for three arbitrary selected tRNA genes, which is similarly RNase-sensitive (Figure S1C).

The ChIP association of Upf1 with protein coding genes was further confirmed by PCR at *pma1* and *act1*, two highly transcribed genes that showed high enrichment in the ChIP-chip datasets. Multiple regions of these genes were examined by radioactive PCR and all showed Upf1 enrichment (Figure 1C and 1D and Figure S2A-2B show *pma1* and *act1* respectively). The association was sensitive to RNase treatment of the chromatin (Figure 1C) (Material and Methods). This binding of Upf1 with *pma1* was also confirmed by ChIP-qPCR in several later experiments using the Flag-tagged *upf1* strain (Figure 1F and Figure S2C). The association with several other active genes was also confirmed by ChIP-qPCR (described in the section below).

### Upf1 chromatin association correlates with Pol II transcription and RNA levels

Upf1 ChIP-chip signals were compared with that of Pol II, similarly calculated from a previously published Ser5 Pol II ChIP-chip dataset (Material and Methods) - Note that unlike in other organisms, Ser5 Pol II is not restricted to promoter-proximal regions in *S. pombe*, it is instead loaded throughout the coding region of active genes (Daulny et al., 2016; Wilhelm et al., 2008). There is a clear correlation between the Upf1 and Pol II ChIP signals at active genes in both the asynchronous and S-phase samples (Spearman’s rank correlation test of 0.56 and 0.58 respectively, Figure 2A).

**Figure 2.**
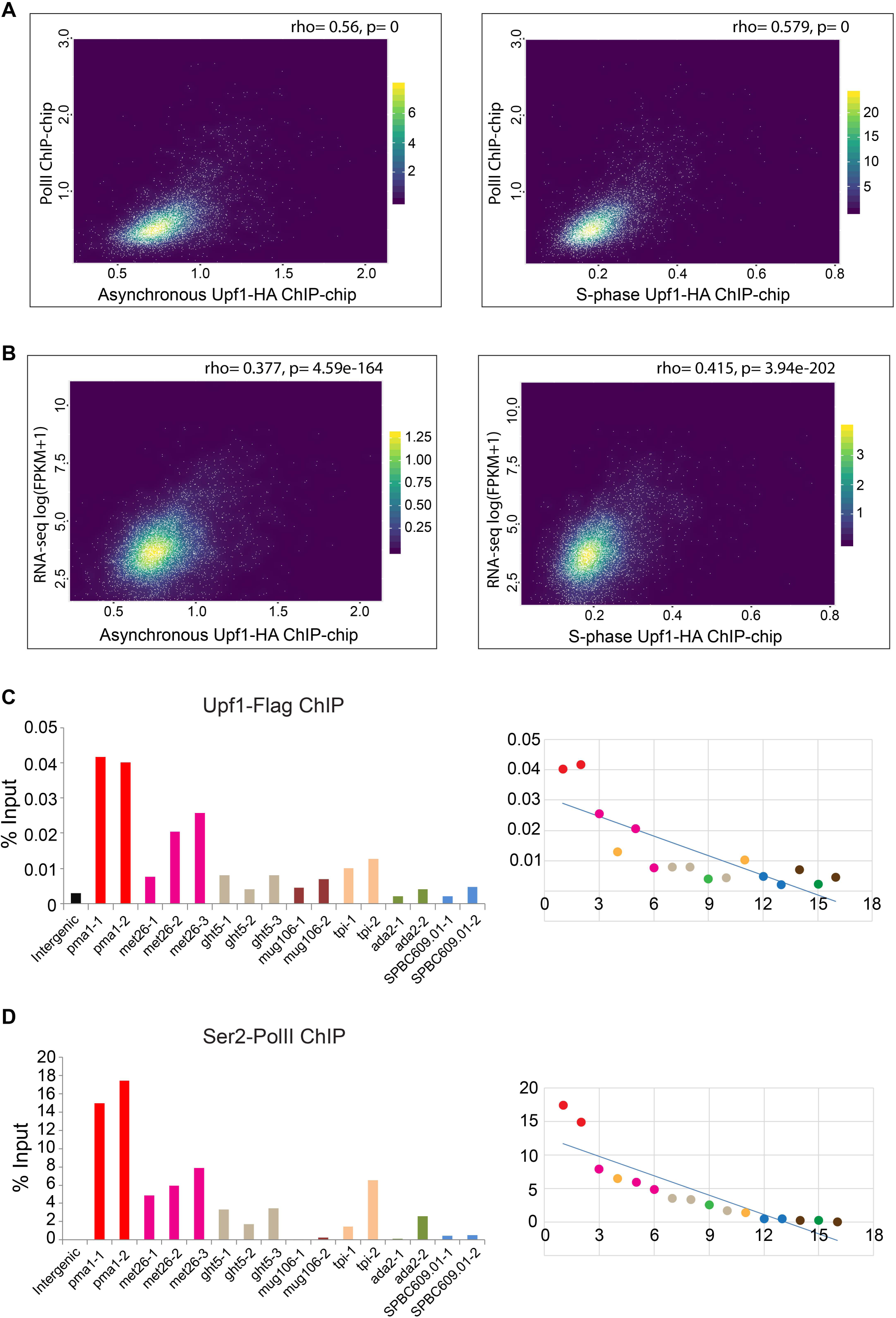
Upf1 association to chromatin positively correlates with Pol II loading and gene expression genome-wide. **A**) Scatter plots showing genome-wide Upf1(x-axis) versus Pol II (y-axis) ChIP-chip signals in asynchronous (left) and S-phase cultures (right). **B**) Scatter plots showing genome-wide Upf1 ChIP-chip signals (x-axis) versus RNA-seq signal (y-axis, log(FPKM + 1) in asynchronous and S-phase cultures. **C**) Left panel-Upf1-Flag qPCR-ChIP signal on specific regions of 7 selected genes and 1 intergenic control (−1, −2 and −3 refer to separate amplicons of the named gene). Genes were selected based on having different levels of Upf1 ChIP-chip association. Right panel-Upf1-Flag ChIP fold enrichment values at specific gene regions plotted according to the Ser2 Pol II ChIP signal, as described in the main text. The primer pairs for each gene are coded with same colour. **D**) Left panel-Ser2 Pol II ChIP signal on the 7 selected genes and the intergenic control. Right panel-the enrichment values for Ser2 Pol II ChIP with each primer pair ordered from high to low values (from left to right). The set of primer pairs for each gene are coded with same colour.

Next we compared Upf1 enrichment values to gene expression by comparing the Upf1 ChIP-chip signal values with RNA-seq FPKM values for all genes (Material and Methods). Again, a positive correlation is observed between the Upf1 enrichment values from both asynchronous and S phase samples and the RNA-seq FPKM values (based on the top 90% expressed genes according to the RNA-seq data) - Spearman’s rank correlation of 0.38 and 0.42, asynchronous and S phase respectively (Figure 2B). These correlations further indicate that the association of Upf1 with gene loci primarily depends on their transcription, both in S phase cells and normal asynchronously growing cells.

The correlation between Upf1 and Pol II enrichment signals is visually apparent on the genome browser at some arbitrarily chosen genes that are associated with Upf1 based on the ChIP-chip data analysis: *pma1*, *met26*, *ght5*, *mug106* and *tpi1* (Figure S3A-S3E). On the other hand, two other randomly chosen genes which show no Upf1 signals, *ada2* and *SPBC609.0, also* displayed no or minimal Pol II signals (Figure S3F-S3G). One exception in this set is *mug106,* which despite having an apparent Upf1 signal throughout the gene, does not show a Pol II signal according to the ChIP-chip data (Figure S3D) – as these data refer to Ser5, it is possible that the Pol II that transcribes this gene is not Ser5 phosphorylated (data discussed further below supports this interpretation). There are also some highly transcribed genes that show no or little association with Upf1, for example, *gpm1*, a gene just downstream of *mug106* (Figure S3D). The reason why Upf1 is not associated with these genes is yet to be determined.

With regard to mRNA expression levels, the selected genes range from *pma1*, one of the most highly expressed in *S. pombe*, to *SPBC609.01* and *mug106* which are expressed at a much lower level in standard growth conditions (Figure S4). The association of Upf1 with all these genes was further examined by ChIP-qPCR in an independent experiment using the Flag-tagged *upf1* strain (Figure 2C). The levels of Ser2 Pol II at these genes was also assessed by ChIP-qPCR (Figure 2D) and these correlate with Upf1 signals at most of the gene regions (Figure 2C right panel shows correlation of Upf1 with Ser2 values taken from Figure 2D; Figure 2D right panel is correlation of Ser2 level with itself, showing the signals’ ranking.

In summary, although there are exceptions, the degree of Upf1 association with active genes correlates genome-wide with Pol II loading and RNA levels.

### Upf1 does not copurify with Pol II

The RNase sensitivity of the ChIP signal discussed above indicates that Upf1 is primarily associated with the nascent transcript. We examined whether Upf1 copurifies with Pol II to investigate potential other interactions between the two. Pol II was purified using a strain encoding the Rpb3 subunit of Pol II functionally tagged with a single copy of Flag (Kimura et al., 2002). In a similar strain Upf1 was also tagged with HA. The Pol II purification procedure (Material and Methods) was validated by silver-stained SDS-PAGE of the Flag elution fraction, which confirmed co-purification of the expected bands corresponding to Rpb1 (two top bands) and most of other Pol II subunits (Figure 3A). There were no apparent experimentally reproducible changes in the protein banding pattern of the Pol II complex purified from wild-type and *upf1Δ* (Figure 3A). The identity of the putative Rpb1 bands was confirmed by mass spectrometry (not shown) and by western blotting (Figure 3B). However, there was no evidence of a putative Upf1 band in the Pol II fraction (Figure 3A). Furthermore, Upf1 could not be detected in the Pol II elution fraction by western blotting of purified Pol II from the Flag-Rpb3/Upf1-HA double tagged strain (Figure 3C, lane 5). There is also no evidence of Pol II copurifying with Upf1 in the reverse experiment using a strain carrying Flag-tagged Upf1 (Figure 3D). These data thus show no evidence of a direct stable interaction of Upf1 with Pol II and that, as the purification was performed under conditions that should keep the nascent transcript intact, the association with the nascent mRNA should be dynamic and it is lost during the purification.

**Figure 3.**
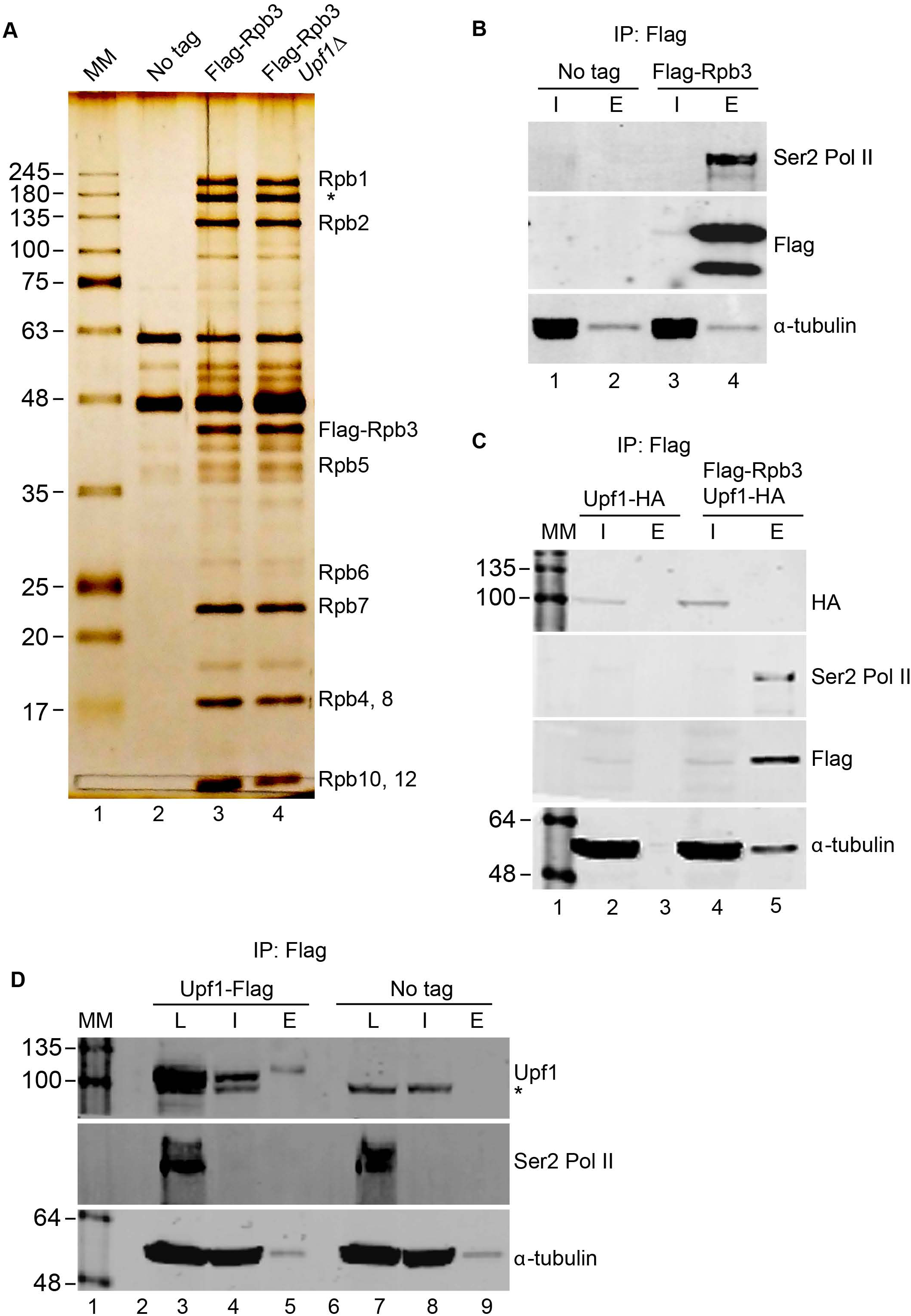
Upf1 does not copurify with Pol II. **A**) Silver stained gel of affinity purified Pol II protein complex using Flag tagged Rpb3 (labelled IP: Flag). Three strains used for this assay were-non-tagged wild-type (lane 2), Flag tagged Rpb3 (lane 3) and Flag tagged Rpb3 in a *upf1Δ* strain (lane 4). Molecular kDa marker (MM) was loaded in the lane 1. Different subunits of the Pol II complex copurifying with Flag tagged Rpb3 are labelled on the right hand side – Note the two bands at the top of the gel correspond to Rpb1, the lower band (asterisk) was confirmed to be also a Rpb1 species by mass spectrometry (data not shown) and could be a cleavage product produced during the purification. **B**) Western blots for Ser2 Pol II, Flag-Rpb3 and *α*-tubulin (as indicated on the right) of protein extract or purified fractions from non-tagged wild-type control (lanes-1 and 2) and Flag-Rpb3 (lanes-3 and 4) strains. Where I is input and E eluted Flag purified proteins. **C**) Western blots for the 4 proteins indicated on the right of each panel from protein extracts or purified fractions from Upf1-HA (lanes-2 and 3) and Upf1-HA+Flag-Rpb3 strains (lanes-4 and 5). Where I is input and E-eluted Flag purified proteins. **D**) Western blots for the 3 proteins indicated of extract/purified fractions from Upf1-Flag (lanes-3, 4 and 5) and non-tagged WT (lanes-7, 8 and 9) strains. Where L is total lysate (prior DNase treatment and centrifugation clearing), I is input (centrifugation supernatant), E is eluted Flag purified proteins. The asterisks indicate a non-specific cross-reacting band.

### There is increased Ser2 Pol II signal at active genes in *upf1Δ*

Next we examined whether there were changes in the genome-wide distribution of total Pol II by ChIP-seq of Flag-tagged Rpb3 in *upf1Δ* and wild-type strains. The ChIP-seq data were processed and metagene plots were produced by taking the coverage signal values from 1kb upstream of the transcription start site (TSS), to 1kb downstream of the transcript end site (TES) and for the gene body of all annotated protein coding genes (Material and Methods). This analysis indicated the expected Pol II gene loading for *S. pombe* in both wild-type or *upf1Δ* with the characteristic increased signal of Pol II downstream of the TES (Figure 4A). This 3’ end skewed total Pol II metagene pattern, which differs from that seen in other organisms, has been previously discussed in *S. pombe* (Daulny et al., 2016; Knoll et al., 2018). This pattern is observed at many individual highly transcribed genes (several examples are shown in Supplementary Data File 1).

**Figure 4.**
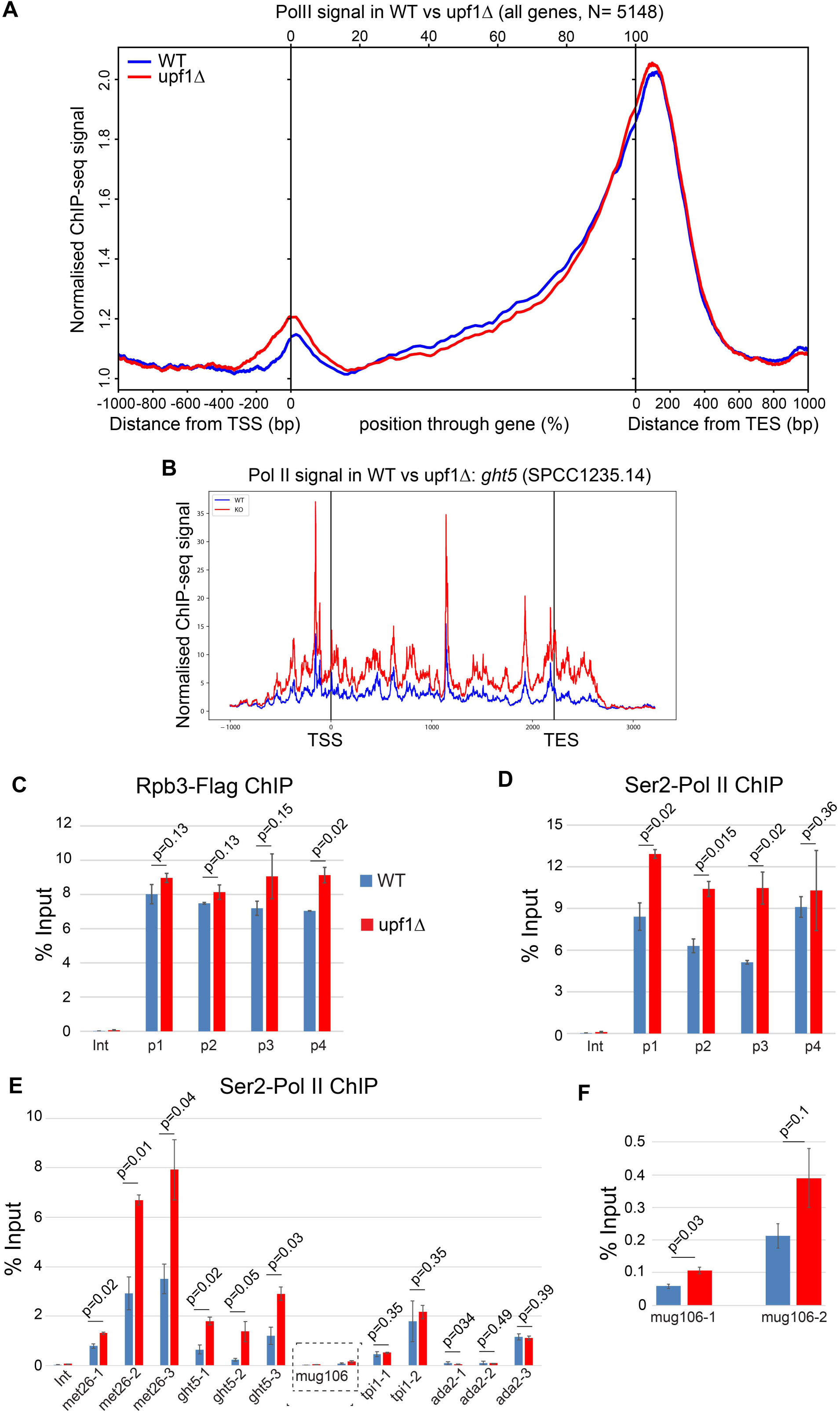
Increased Ser2 Pol II loading on active genes. **A)** Metagene analysis of genome-wide binding of total-Pol II from ChIP-seq performed with wild-type (WT) and *upf1*Δ strains. The averaged profiles for all the genes are displayed. The analysed region covers the sequence 1000 bp upstream of the TSS, the ORF region and 1000 bp downstream of the TES. **B**) Profiles of total-Pol II ChIP-seq signals at *ght5* in WT and *upf1*Δ (termed KO in this panel) strains. Units on x-axis are base pairs relative to the gene TSS and TES sites. **C**) qPCR validation of Flag-Rpb3 ChIP signal on *pma1* in WT and *upf1*Δ strains. Results are shown as percentage of input enrichment for the average of three independent biological samples with three replicates each (mean±SEM), the signal at an intergenic (Int) region is shown as background control for each replicate. Statistical analysis of differential Flag-Rpb3 accumulation was performed using one-tailed Student’s t-test. P-values are shown above each primer pair. **D**) Ser2 Pol II ChIP signal on *pma1* in WT and *upf1*Δ strains. Statistical analysis of differential Ser2 Pol II accumulation was calculated as mentioned above. **E**) Flag-Rpb3 ChIP signal on 5 genes in WT and *upf1*Δ strains. **F**) Flag-Rpb3 ChIP signal on *mug106* gene. Statistical analysis of differential Flag-Rpb3 accumulation was calculated as mentioned above.

This analysis indicates that at most of the genes there are no major changes in total Pol II loading between wild-type and *upf1Δ* (Figure 4A), other than a possible slightly increased signal proximal to the TSS. However, there are a few genes, among those strongly associated with Upf1 in wild-type cells, that show increased total Pol II signal in *upf1Δ*, either in the gene body, or around the TES or downstream of the TES (Supplementary Data File 1). Several of the genes displaying this increased Pol II association are also mis-regulated in *upf1Δ* (Supplementary Data File 2), a striking example is seen at *ght5* (Figure 4B), which is a moderately expressed gene that is associated with Upf1 in wild-type (Figure S3), and one that is up-regulated in *upf1Δ* (discussed further below).

Both ChIP-seq and ChIP-qPCR data showed that total Pol II loading does not obvious change at *pma1* in *upf1Δ*, with the possible exception of a small increase in the 3’ flanking region (Figure 4C). However, the levels of Ser2 Pol II signal are significantly increased throughout the gene barring the 3’ flanking region (Figure 4D). Higher Ser2 signal is also seen at the two other highly active genes tested: *met26*, and *ght5* (Figure 4E). There could additionally be a small Ser2 Pol II signal increase at the low transcribed *mug106* gene (Figure 4F).

### There is a significant overlap between genes bound by Upf1 and genes differentially expressed in *upf1*Δ cells

To explore further whether Upf1 may have some function in the expression of the genes to which it is associated, we compared these genes with genes that are differentially expressed in *upf1Δ*. We analysed a previous RNA microarray dataset (Rodriguez-Gabriel et al., 2006), and used significance analysis of microarrays (SAM) with a 1% FDR to find differentially expressed genes between the wild-type and upf1Δ samples (Material and Methods). We identified a total of 543 genes differentially expressed between the wild-type and *upf1Δ* using these parameters. Of these, 159 show reduced mRNA levels, whereas almost double this number (384) of genes show increased mRNA levels in *upf1Δ* (Figure 5A and Table S2). Of the 543 differentially expressed genes, 47 are also strongly bound by Upf1 according to our ChIP-chip data (Figure 5B; red and green codes indicate up and down regulated genes, respectively). Based on the number of different genes represented on the microarrays we calculated the p-value of this overlap to be 0.001 (Material Methods). Notably, two of these genes are *met26* and *ght5*, which as described above show increased Ser2 signal, and in the case of *ght5*, increased total Pol II in *upf1Δ* compared to wild-type. Both genes are upregulated in *upf1Δ* (indicate in blue in Figure 5B). Note that although *pma1* is not misregulated, an antisense ncRNA gene (SPNCRNA.92) located in the 5’ UTR of *pma1* is downregulated in *upf1Δ* (Figure 5B). This ncRNA gene is strongly associated with Upf1 according to the ChIP-chip data (Fig. 1B). The RNA level of several Tf2 retrotransposons, which might also be associated with Upf1, as discussed, are also increased in *upf1Δ* according to both the microarray dataset and qRT-PCR validation (Figure 5B and Figure S1A, S1B). Notably, one of the Tf2 elements (Tf2-5, SPAC2E1P3.03c) also shows increased total Pol II loading in *upf1Δ* cells compared to wild-type, including in the non-repetitive 3’ end region, suggesting that Tf2-5 may be transcriptionally up-regulated in these cells (Supplementary Data File 2).

**Figure 5.**
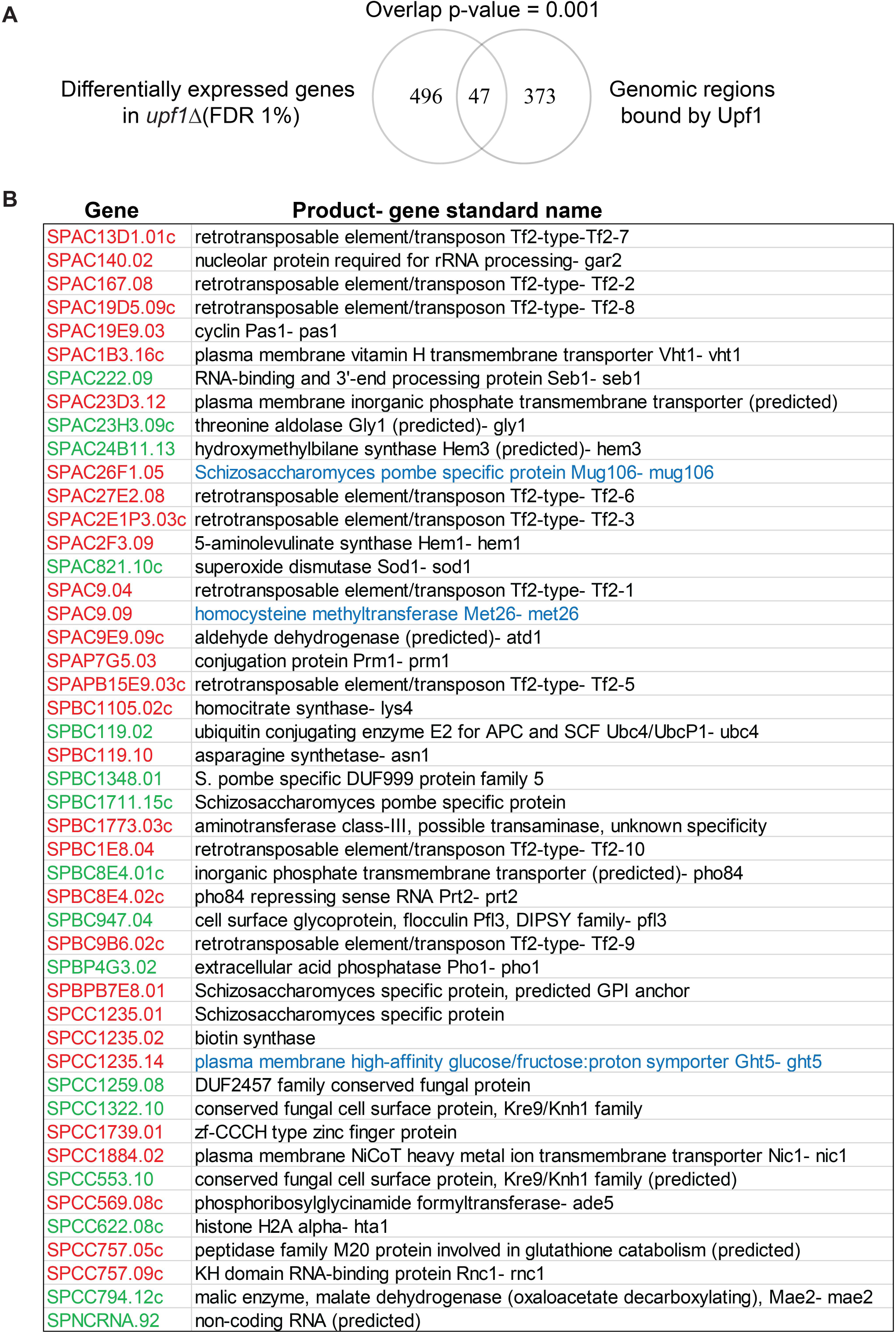
Significant overlap between genes bound by Upf1 and genes differentially expressed in a *upf1*Δ strain. **A**) Venn diagram showing the overlap between the genes/genomic regions strongly associated with the Upf1 (420, identified using the p-value cut off of 10^-4^ in the MAT software) and differentially expressed genes in *upf1*Δ (543). This overlap of 47 genes corresponds to a p-value of 0.001 **B**) List of the 47 overlapping genes. Genes highlighted in red are upregulated and genes highlighted in green are downregulated. Genes which were tested and show increased Ser2 CTD phosphorylation are show with their full names in blue.

### Upf1 deficient cells are hypersensitive to 6-Azauracil

To explore further whether Upf1 has a role in Pol II transcription, we examined whether *upf1*Δ cells are hypersensitive to 6-azauracil (6AU). It has previously been reported that strains carrying mutations in components of the RNA polymerase II transcription elongation machinery are hypersensitive to 6AU (Shaw and Reines, 2000; Zhou et al., 2015). 6AU is an inhibitor of enzymes that are involved in nucleotide biosynthesis; 6AU treatment leads to nucleotide depletion and hence can diminish transcription elongation (Exinger and Lacroute, 1992). When grown in 0.8 mM 6AU, frequent morphology and septation defects were observed in *upf1*Δ but not in the wild-type strain, with the appearance of long unseparated chains of cells (Figure 6A, panels II vs IV). These phenotypes, although they were not quantified, are similar to those previously described for the elongation mutants referred above. These appear 2 hours after addition of the drug under standard growth conditions and persisted at all later time points examined up to 3 hours (not shown). The *upf1Δ* strain also shows a slow growth phenotype in the presence of 6AU, in both liquid cultures and on agar plates (Figure 6B and 6C).

**Figure 6.**
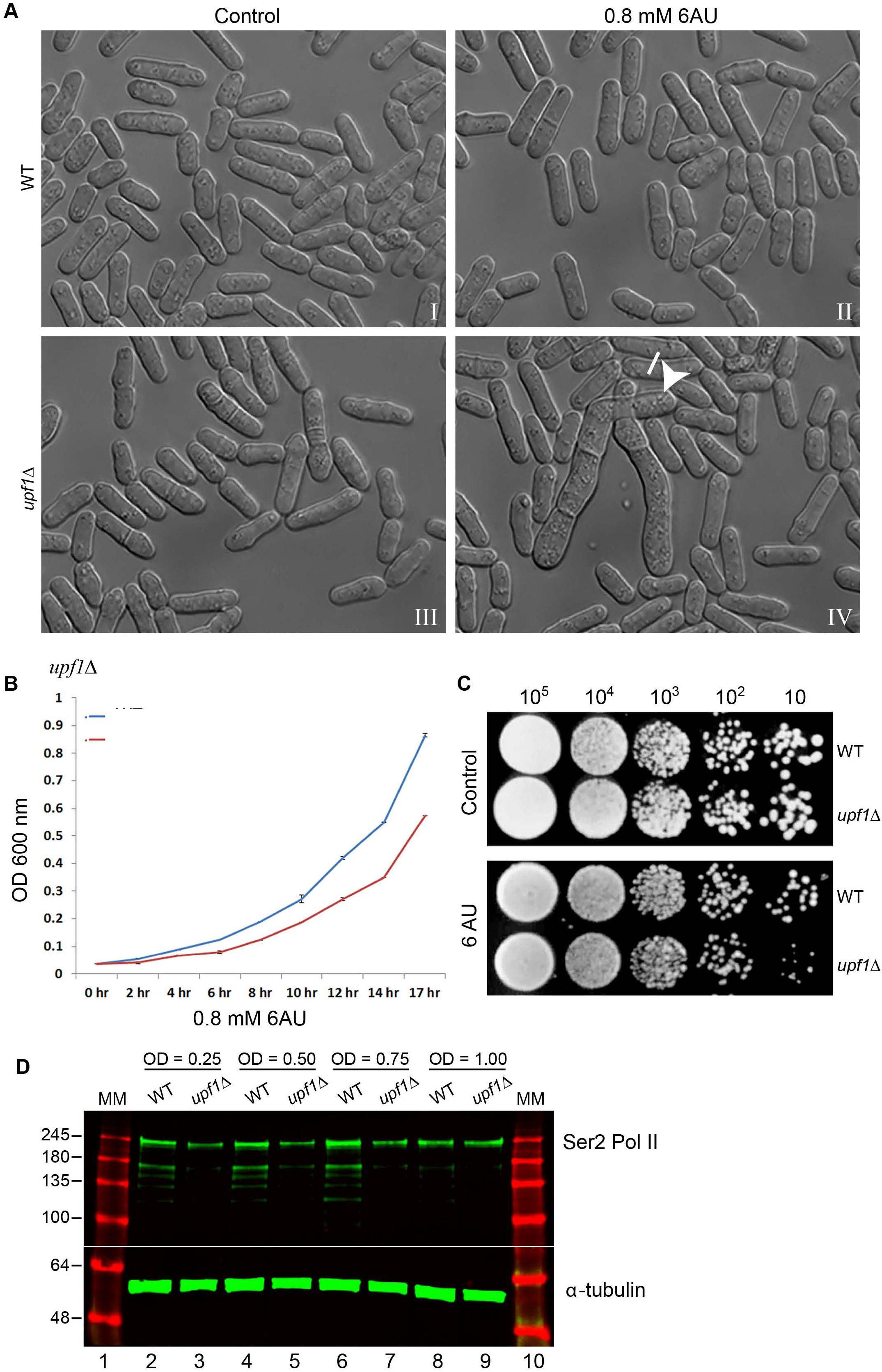
Upf1 deficient cells are hypersensitive to 6-Azauracil. **A**) Photomicrographs of WT and *upf1*Δ *S. pombe* cells growing in standard YES (left panel) or YES+0.8 mM 6AU (right panel) medium. Long, unseparated chains of cells are indicated with a white arrow. **B**) Growth assay curves of WT and *upf1*Δ cells grown in YES+0.8 mM 6AU media. X-axis shows the different time points at which OD_600_ was measured. **C**) Serial dilution colony growth assay of the WT and *upf1*Δ cells, spotted on YES and YES+0.8 mM 6AU plates. D) Western blotting of total S. pombe protein extract from cells at different time points from a growing culture (the OD_600_ of the culture at each point is indicated above). The top half of the membrane was incubated with a Ser2 Pol II antibody and the bottom half with an alpha-tubulin antibody (as loading control). Lanes 1 and 10 show the same protein molecular weights marker (values are in kilodaltons).

### Pol II abnormalities in *upf1Δ*

It was also examined whether there are changes in Pol II phosphorylation by western blotting of whole-cell lysates of cells taken from a growing culture at different intervals. A Ser2 specific antibody was used, which is expected to detect Ser2 phosphorylates CTD of Pol II largest subunit Rpb1. The slowest/top migrating species that this antibody detects should represent Rpb1 with a fully phosphorylated CTD. Notably, it appears that the largest band that the Ser2 antibody detects is sharper and slightly upward shifted in *upf1Δ* compared to wild-type (Figure 6D). This mobility shift is most apparent in cells taken at the start of the fastest growing stage of the culture (Figure 6D, O. D. 0.5, lanes 4 vs. 5 – unless otherwise specified, all the experiments described in this study were performed with culture at ∼ O.D. 0.5). The overall level of Ser2 seems lower in *upf1Δ* barring in the densest culture examined (O.D 1, lanes 8-9). There might also be less of the cross-reacting faster migrating bands in *upf1Δ*; these could represent Rpb1 cleavage products of different sizes. Consistent with this interpretation, the largest of these products is also prominent in the SDS-PAGE of affinity purified Pol II fractions and its identity was validated by mass-spectrometry (Figure 3A, indicated by the asterisk). These data indicate that Pol II and CTD phosphorylation might be abnormal in *upf1Δ,* particularly in fast growing cells.

## Discussion

The RNA helicase Upf1 has been mainly studied for its cytoplasmic role in NMD in *S. pombe* as well as in other model eukaryotes. In contrast to this broadly accepted view, we provide evidence that Upf1 is associated genome-wide with active genes in *S. pombe*. The association is mostly with protein-coding genes, RNase sensitive and positively correlates with Poll II loading as well as mRNA expression levels at most genes.

There are no obvious changes in total Pol II loading at the vast majority of genes in *upf1*Δ cells, other than a possible small increase proximal to the TSS (Figure 4A). However, Pol II loading is clearly increased throughout the transcribed region at a number of genes with which Upf1 clearly associates in wild-type (for example at *ght5* and at other genes, Figure 4F and Supplementary Data File 1, respectively). Notably, most of the genes examined have an increased Ser2 ChIP-qPCR signal in *upf1*Δ. Since there is no/or little change in total Pol II loading at most of these genes (as determined in more detail for *pma1*) the higher Ser2 signal might be due to Ser2 CTD hyperphosphorylation rather than the genes being more densely loaded with Pol II in *upf1*Δ. Consistent with hyperphosphorylation, the band corresponding to Ser2 phosphorylated Pol II (Rpb1) migrates slower in *upf1*Δ compared to wild-type. It is therefore likely that more of the CTD repeats are phosphorylated in wild-type compared to *upf1*Δ. Ser2 hyperphosphorylation has previously been linked to slow transcription elongation in mammalian cells (Fong et al., 2017).

Although there is no obvious change in total Pol II loading by ChIP, the possibility that Pol II elongation is altered in *upf1*Δ cannot be ruled out. As discussed, 6AU hypersensitivity is a characteristic of transcription elongation *S. pombe* mutant strains (Zhou et al., 2015). Additionally, *S. pombe* strain carrying a mutant Pol II with reduced elongation rate is also hypersensitive to 6AU (Yague-Sanz et al., 2020). Perhaps the role of Upf1 in transcription is more important in conditions of stress; exemplified by the nucleotide depleting conditions that the drug 6AU induces. Alternatively, it is plausible that CTD hyperphosphorylation is a consequence or a functional output of some yet unknown broad compensatory mechanism that maintains almost-normal transcription elongation rate in some abnormal/mutant conditions. For example, when the formation of the nascent mRNP and transcription are not correctly coupled such as in *upf1*Δ. Whether this putative compensatory mechanism relates to the mRNA decay dependent transcription adaptation phenomenon recently described in zebrafish, remains a possibility to be investigated in future studies (El-Brolosy et al., 2019).

In summary the data we have discussed indicate that Upf1 interacts with the nascent transcript, playing a role in transcription-coupled processes in *S. pombe*. We envisage that by a yet undefined cross-talk mechanism, Upf1 may take part in a feedback system between the formation of the nascent mRNP and Pol II CTD phosphorylation. A model is that by operating on the mRNA, Upf1 straightens the nascent transcript preventing its entanglement, hence transcription stress and Ser2 hyperphosphorylation (Figure 7).

**Figure 7.**
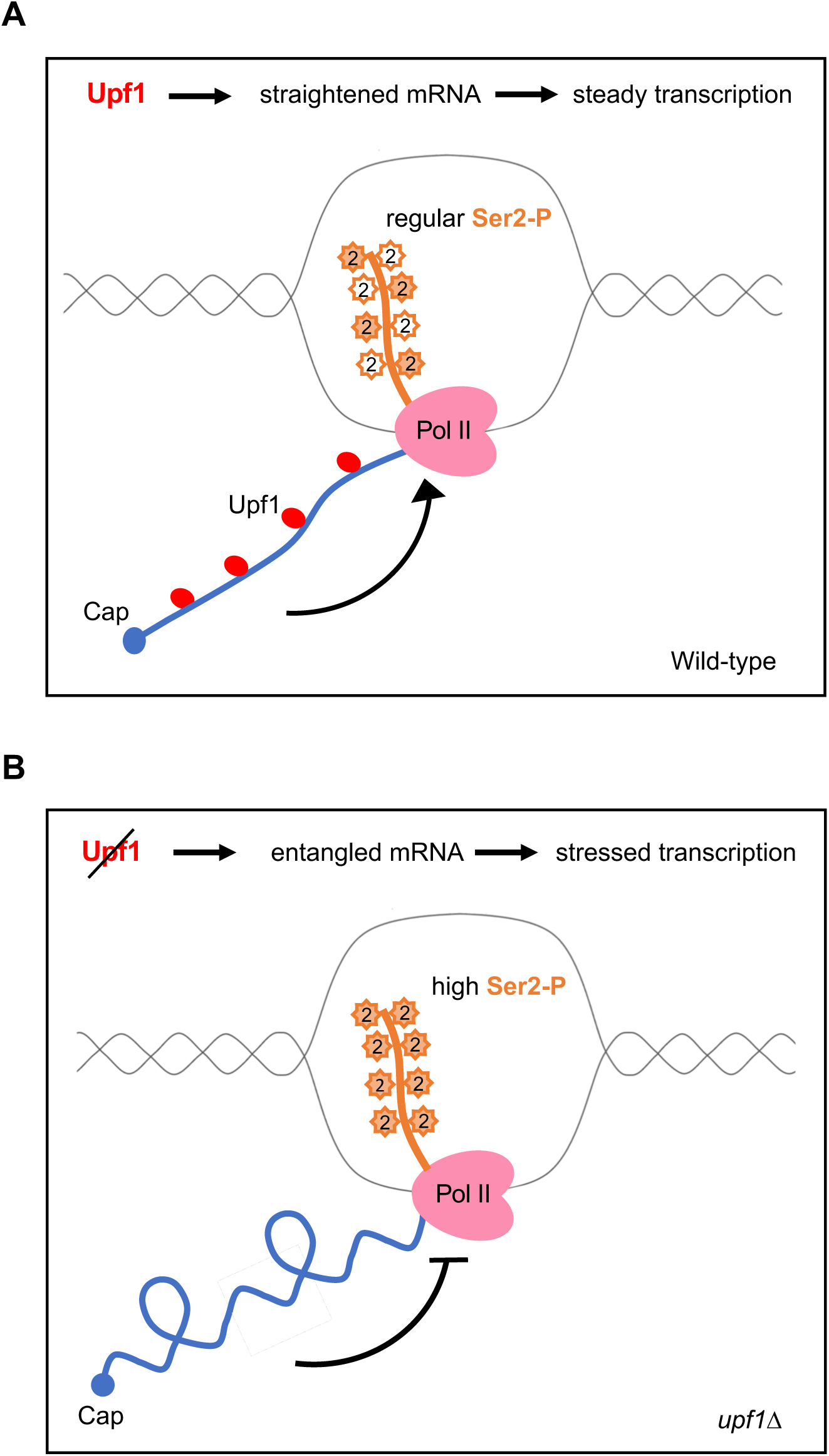
Upf1 association with the nascent mRNA controls transcription. **A)** Schematic of the site of transcription in wild-type cells. Upf1(red) by operating on the mRNA, straightens the nascent transcript. This permits steady transcription and normal serine 2 phosphorylation at the Pol II (pink) carboxy-terminal domain (CTD) (orange, coloured stars mean phosphorylated, white starts mean unphosphorylated). **B)** Schematic of the site of transcription in *Upf1Δ* cells. The absence of Upf1 results in entanglement of the nascent mRNA leading to stressed transcription and CTD serine 2 hyperphosphorylation (orange stars).

The data we have presented are in agreement with the recent report that Upf1 is associated genome-wide with nascent transcripts in *Drosophila* (Singh et al., 2019). Taken together, the findings from these two highly divergent organisms predict that the roles of Upf1 in transcription-coupled processes are conserved in eukaryotes. Species specific differences are likely though. For example, apart from the slightly increased Pol II loading at TSS-proximal sites, which was also observed in *Drosophila*, there was no obvious evidence of increased Ser2 phosphorylation in UPF1 depleted cells in this organism (Singh et al., 2020). In Drosophila, UPF1 might play a positive role in Ser2 phosphorylation (Singh et al., 2020).

Furthermore, these data re-prompt questioning the fields’ communally accepted syllogism that transcripts whose levels are increased in Upf1 depleted cells are therefore likely to be NMD targets. This might not always be a valid explanation: it has previously been argued that only a fraction of these increases might actually be a direct consequence of cytoplasmic mRNA stabilisation (Brogna et al., 2016; Brogna and Wen, 2009; Wen and Brogna, 2010; Wen et al., 2020). Here we have shown that there is overlap between genes associated with Upf1 and genes that have increased mRNA levels in *upf1*Δ. Such increases in mRNA levels would have, directly or indirectly, previously been attributed to NMD suppression in the cytoplasm. This explanation cannot be ruled out. However, it is plausible that the increased cellular mRNA levels might be caused by the absence of Upf1 from the transcription site of an affected gene, resulting in its abnormal expression. Some of the affected genes are apparently transcriptionally up-regulated in *upf1*Δ, as discussed. Whether the apparent upregulation of these set of genes is a direct consequence of the absence of Upf1 from their transcription sites, or indirectly, due to mis-expression of transcription regulators involved in their expression or due to some transcription adaptation mechanism remains to be addressed.

## Material and Methods

### Yeast strains and methods

The complete list of *S. pombe* strains used in this study is shown in Table S3. Fission yeast transformation was carried out as described earlier (De et al., 2011). The target proteins were HA and FLAG-tagged by homologous recombination using a PCR-amplified fragment containing the kanMX6 or hphMX6 cassette flanked by targeting sequences (Bahler et al., 1998); all the PCR primers used for tagging the target genes are listed in Supplemental Table S4.

### Western blotting

Protein extraction from *S. pombe* cells was done as described before (Matsuo et al., 2006). Membranes were probed with the required primary antibodies: rabbit anti FLAG (F7425, Sigma Aldrich), Rat anti-HA (11867423001, Sigma Aldrich), mouse anti-alpha-tubulin (AB_477579, Sigma-Aldrich, 1:2500), rat anti-Ser2 Pol II (AB_11212363, Merck Millipore, 1:5000). Respective secondary antibodies, IRdye 800 and/or 680 were used to detect the signal using an Odyssey infra-red imaging system (LI-COR Biosciences).

### ChIP

Freshly harvested cells from exponentially growing cultures (OD_600_=0.5) were fixed for 5 minutes at room temperature with 1% formaldehyde (Sigma Aldrich) followed by 10 minutes incubation with a further addition of 2.5 M glycine. The cell pellet was collected and washed twice with ice-chilled 1X PBS with spinning at 5000 rpm for 3 min each. The pellet was resuspended in ice-cold FA lysis buffer [HEPES-KOH-100 mM (pH 7.5), NaCl- 300 mM, EDTA- 2mM, Triton X-100- 2%, Na-Deoxycholate- 0.2%] containing 1X protease inhibitor (EDTA-free protease inhibitors cocktail tablet, Roche). Cells were pelleted at 6000 rpm for 2 minutes at 4°C and the pellet was resuspended in FA lysis buffer and Zirconia beads (0.7 mm diameter, Biospec). Cells were broken using a cell homogenizer (Bertin Instruments, Precellys 24, 10 cycles: 30 seconds at 5500 rpm and 2 minutes in ice). The bottom of each screw cap tube was pierced three times with a red-hot 25 G needle and each tube was immediately transferred to the barrel of a syringe fitted in a 15 ml falcon tube. The lysate was collected at 1000 rpm for 1 minute at 4°C. To increase sonication efficiency and prevent proteases, 20 µl of 10% SDS and 20 µl of 100 mM PMSF were added to the mixture.

Samples were sonicated for 15 cycles using a Bioruptor (Diagenode), to generate ∼ 500bp average fragment size. Immunoprecipitation was done by adding Dynabeads (Thermofisher) and incubated overnight at 4°C on a rotor. The supernatant was removed and beads were washed for 5 minutes at room temperature on a rotor using buffers as mentioned: Wash Buffer I [HEPES-KOH- 50 mM (pH 7.5), NaCl- 150 mM, EDTA- 1mM (pH 8.0), Triton X-100- 1%, Sodium deoxycholate- 0.1%, SDS- 0.1%)- 2 times; Wash Buffer II [HEPES-KOH-50 mM (pH 7.5), NaCl- 500 mM, EDTA- 1 mM (pH 8.0), Triton X-100- 1%, Na-deoxycholate- 0.1%, SDS- 0.1%] - 2 times; Wash Buffer III [Tris-HCl- 10 mM (pH 8.0), EDTA- 1 mM (pH 8.0), LiCl- 0.25 mM, IGEPAL CA630- 0.5%, Na-deoxycholate- 1%]- 2 times and TE [Tris-HCl- 10 mM (pH 8.0), EDTA- 1 mM (pH 8.0)]- 2 times. After the final wash, beads were resuspended in 100 µl Elution Buffer (EB) [Tris-HCl- 50 mM (pH 7.5), EDTA-10 mM, SDS- 1%] and incubated for 10 minutes at 65°C and occasionally vortexed. The supernatant (elution) was recovered and transferred to a fresh 1.5 ml DNA low bind tube. To the input, 150 µl EB was added and incubated at 65°C overnight to allow de-crosslinking. The IP sample was de-crosslinked in parallel using the same condition. To remove proteins from the DNA, 5 µl Proteinase K (20 mg/ml) was added and samples were incubated at 50°C for two hours. DNA was then extracted using the Monarch PCR purification kit, as previously described (Singh et al., 2019).

RNase ChIP, radioactive PCR and ChIP-chip was carried out as previously described (De et al., 2011). qPCR quantification of DNA samples was carried out using the SensiFAST SYBR Hi-ROX Kit (Bioline, BIO-92005) in 96-well plates using a ABI PRISM 7000 system (Applied Biosystems). For ChIP-seq, all ChIP-DNA libraries were produced using the NEBNext Ultra II DNA Library Prep Kit (NEB, E7645L) and NEBNext Multiplex Oligos for Illumina (NEB, E7600S), using provided protocols with 10 ng of fragmented ChIP DNA. Pipetting was done with a Biomek FxP robotic work station (Beckman Coulter, A31842). Constructed libraries were assessed for quality using the TapeStation 2200 (Agilent, G2964AA) with High Sensitivity D1000 DNA ScreenTape (Agilent, 5067-5584).

### Analysis of ChIP-chip data

We used the Model-based Analysis of Tiling Arrays (MAT) software to analyse the Affymetrix hybridization data (Johnson et al., 2006). ChIP input DNA sample was used as control and was compared against the Upf1 (asynchronous and S-phase) and Pol II samples. A p-value cut off of 10^-4^ or 10^-3^ was used, whereas the remaining MAT parameters remained as default. Results of MAT were visualised in Affymetrix’s Integrated Genome Browser (IGB) (Nicol et al., 2009). When 50% or more of a genomic region was significantly bound by Upf1 and Pol II, we called it an enriched gene/genomic region. Enrichment scores were assigned to genomic features using the *S. pombe* genome coordinates (ftp://ftp.sanger.ac.uk/pub/yeast/pombe/GFF). The average enrichment was calculated between the start and end coordinates of enriched genomic regions, thereby giving each enriched region a score based on fold enrichment. Identification of significantly bound genomic features and enrichment score calculation was done using the statistical computing language R (http://www.R-project.org/). Functional annotation of the enriched regions was done using DAVID (Dennis et al., 2003).

### Pol II and Upf1 purification

Exponentially growing cultures (OD600=0.5) of healthy *S. pombe* cells were pelleted down for 10 min at 5000 rpm at 4°C (Rotor: F10-6X500, FiberLite Beckman J2-MC). The pellet was washed with buffer 1 (HEPES- 20 mM, KAc- 110 mM) and resuspended in lysis buffer [HEPES- 20 mM, KAc- 110 mM, TritonX-100- 0.5%, Tween20- 0.1%, MnCl_2_- 10 mM, PMSF- 1 mM, protease inhibitor- 1X, PhosSTOP-1X (Roche), RNase inhibitor −50 U/ml, RVC-10 mM]. Small droplets of the lysate were made in liquid nitrogen and immediately kept at −80^0^C, they were typically processed the day after. Cells were subjected to grinding with SPEX Sampleprep freezer mill- 6775 (Grind cycle: precool- 2 min, 1 cycle of 1 min cooling & 2 min grinding, impact rate 14). An equal volume of lysis buffer and 110 U/ml of DNase (DNase I recombinant, RNase-free solution, Cat no. 4716728001, Roche) was added to the ground cell lysate followed by incubation for 1 hour at 4°C. The sample was centrifuged at 16000g for 10 min and the supernatant was transferred to a fresh tube. The supernatant (input) was incubated with 5 µg of anti-Flag antibody-coated Dynabeads for 1 hr at 4°C on a rotator. After incubation, Dynabeads were washed 6 times for 10 mins each with wash buffer (HEPES- 20 mM, KAc- 110 mM, TritonX-100- 0.5%, Tween20- 0.1%, RNase inhibitor- 50 U/ml, MgCl_2_- 4 mM). Beads were incubated with elution buffer (lysis buffer, MgCl_2_- 4 mM, Flag-peptide- 2 mg/ml) for 30 min at 4°C on a rotator. The elution fraction was collected by separating the beads using a magnet.

### ChIP-chip data processing and correlation analysis

For the ChIP-chip data correlation analysis (Figure 2A-2B), the raw ChIP-chip probe intensities were processed according to the data preparation and expression value calculation sections of the Affymetrix statistical algorithms description document (http://tools.thermofisher.com/content/sfs/brochures/sadd_whitepaper.pdf) in order to obtain a signal value for each gene for each array. The previous Ser5 Pol II ChIP-chip datasets were downloaded from https://www.ebi.ac.uk/arrayexpress/experiments/E-MTAB-18/ (Wilhelm et al., 2008). The signal values in the two pairs of input control datasets (one pair for the Upf1 IPs and one for the Pol II IPs) also underwent the following additional processing. For each pair of control datasets, the two sets of signal values were scaled to each other so that they had the same median, before taking the mean to obtain a single signal value for each gene for each pair (one for Upf1 controls and one for Pol II controls). The pair of ChIP-chip datasets with Pol II IP were also combined into one in the same manner described for the control dataset pairs. One of the two datasets from each of the asynchronous and S-phase Upf1 pairs of IP datasets were found to be of low quality. Therefore, only the higher quality dataset from each pair was used. The signal value of each gene in each IP dataset was then normalised by dividing by the signal value of that gene in the appropriate control dataset. These normalised signal values were used for the correlation analysis.

### ChIP-seq

Exponential cultures of *S. pombe* growing at 30°C in 400 ml YES with an OD600 of 0.8 were fixed by 1% formaldehyde at room temperature for 5 min followed by the addition of 0.125 M glycine. The fixed cells were washed with ice-cold PBS, resuspended in cell lysis buffer [HEPES- 50 mM (pH 7.6), EDTA- 1 mM (pH 8.0), NaCl- 150 mM, Triton X- 100- 1%, Na-Doc- 0.1% and 1X protease inhibitor (EDTA-free protease inhibitors cocktail tablet, Roche], and lysed by acid washed glass beads. Chromatin extracted from cell lysates was fragmented by 8 sonicating cycles of 5 min with 30s ON/30s OFF at HIGH setting (Bioruptor® Plus). Immunoprecipitations were performed using Protein G Dynabeads (Life Technologies) coated with 10 µg monoclonal anti-FLAG M2 antibody (Sigma). Both IP and Input DNA were purified using MinElute PCR Purification Kit (QIAGEN). 10 ng of DNA was used for DNA library construction with the NEBNext Ultra II DNA Library Prep Kit (New England Biolab E7645L), indexed using NEBnext Multiplex Oligos for Illumina Dual Index Primers (New England Biolabs E7600S) and sequenced simultaneously using a Illumina HiSeq4000 System.

### ChIP-seq data metagene analysis

The sequence reads in the FASTQ files were trimmed using Trimmomatic to remove low quality reads (Bolger et al., 2014). The SE (single end) setting was used, with sliding window and minimum read length set to 4:22 and 32, respectively, while all other parameters were set to default. The trimmed FASTQ files were then converted to SAM files by aligning the reads to the EF2 *S. pombe* genome build (Ensembl), which was downloaded from (https://emea.support.illumina.com/sequencing/sequencing_software/igenome.html?langsel<u>=/ gb/</u>). The alignment was carried out using Bowtie2 (Langmead and Salzberg, 2012), which was set to automatically filter out unaligned reads using “--no-unal” option. The single-base resolution genome-wide coverage depth for each file was obtained by finding the number of reads mapping to each base position and dividing by the total number of aligned reads for that file to normalise for sequencing depth. The coverage value for each base position in each IP file was then normalised by dividing by the value for that base position in an input control file, which was used for all normalisations. The normalised base-wise coverage values for each of the two pairs of Pol II ChIP samples (in wild-type and *upf1*Δ cells) were averaged to obtain one set of Pol II coverage values for each cell type.

### Gene expression levels quantification

Two replicate *S. pombe* RNA-seq datasets were downloaded from the Gene Expression Omnibus, with sample IDs GSM2803075 and GSM2803077 from the series with ID GSE104546 (https://www.ncbi.nlm.nih.gov/geo/query/acc.cgi?acc=GSE104546), previously described (Gallagher et al., 2018). A single mean FPKM value was obtained for each gene by averaging the FPKM value for that gene from the two replicate datasets. The values were transformed via log (FPKM + 1), which was used for plotting and correlations.

### Identification of differentially expressed genes

Previous whole-genome microarray RNA expression data of *upf1Δ* and wild-type strains data was used (Rodriguez-Gabriel et al., 2006). Differentially expressed genes were identified from these datasets using significance analysis of microarrays (SAM) at time point 0 between wildtype and *upf1Δ* using a 1% FDR (Tusher et al., 2001). The overlap between differentially expressed genes and Upf1 associated genes was calculated by random sampling, as following: 1) sampling of 420 genes (corresponding to the number of enriched genes at the more stringent p-value threshold of 10^-4^ of the MAT software) from 7054 total annotated genes in the version of the genome analysed; 2) sampling 543 genes from 5280 (total genes tested by Rodriguez et al, 2006); 3) calculating the overlap between 420 and 543 randomly selected gene sets; 4) create an overlap distribution by repeating steps 1 – 3 1000 times, and calculate the p-value from where the true overlap value (47) falls in the distribution.

### Data and code availability

The ChIP-chip and ChIP-seq datasets as well as all associated metadata files are available from a Gene Expression Omnibus (GEO) SuperSeries record: GSE169425 - https://www.ncbi.nlm.nih.gov/geo/info/linking.html.

Description of the bioinformatics pipelines used, custom-made scripts developed for the correlation and metagene plots as well as the raw data files and processed data tables are available from GitHub repository: https://github.com/Brogna-Lab/PombeUpf1

## Acknowledgments

We thank John Arrand in the School of Cancer Sciences for doing the DNA microarray hybridization and Stephen Kissane, John Colbourne and Archana Sharma-Oates for NGS sequencing, technical and bioinformatics troubleshooting. We would also like to thank Vahid Aslanzadeh, and Jean Beggs for discussions and sharing unpublished observations. Thanks also to thank Laura O’Neill for advice on the ChIP experiments and Mike Tomlinson for providing Odyssey infrared imaging system for western blot detection. This project was funded by the following grants: Wellcome Trust 9340/Z/09/Z, BBSRC BB/M022757/1 and BBSRC BB/S017984/1 to SB; BBSRC BB/M017982/1 and BBSRC BB/L006340/1 to DH; Wellcome Trust 202115/Z/16/Z, Royal Society RG170246 and BBSRC BB/S016155/1 to MS. SD and JW were supported by Darwin Trust PhD scholarships; and DME and HLD by MIBTP-BBSRC scholarships.

---

**Figure S1.**
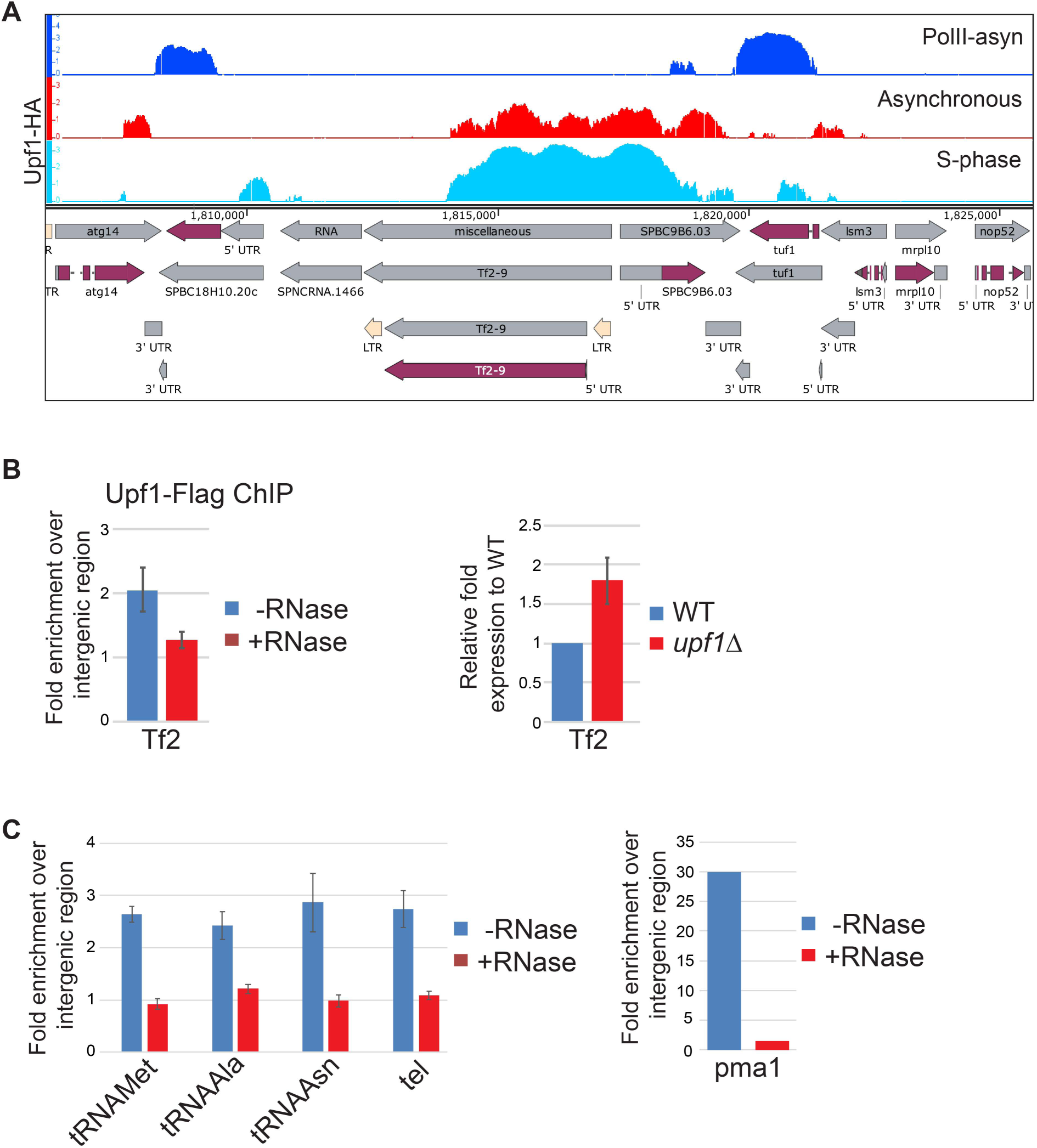
Upf1 association to tRNA genes and transposable elements loci is RNase sensitive. **A**) IGB screenshot of ChIP-chip enrichments of Pol II in asynchronous culture (top row), Upf1-HA in asynchronous culture (middle row) and in S-phase culture (bottom row) over Tf2-9 gene and it’s flanking region of *S. pombe*. Genes and genomic features are shown below. **B**) Left panel-Upf1-Flag qPCR-ChIP signal of a transposable element Tf2 sequence with and without RNase treatment. Right panel-expression of Tf2s in WT vs *upf1*Δ cells. **C**) Left panel-Upf1-HA ChIP signal on tDNAs and telomere. Right panel-Upf1-HA ChIP signal on *pma1* without and with the RNase treatment.

**Figure S2.**
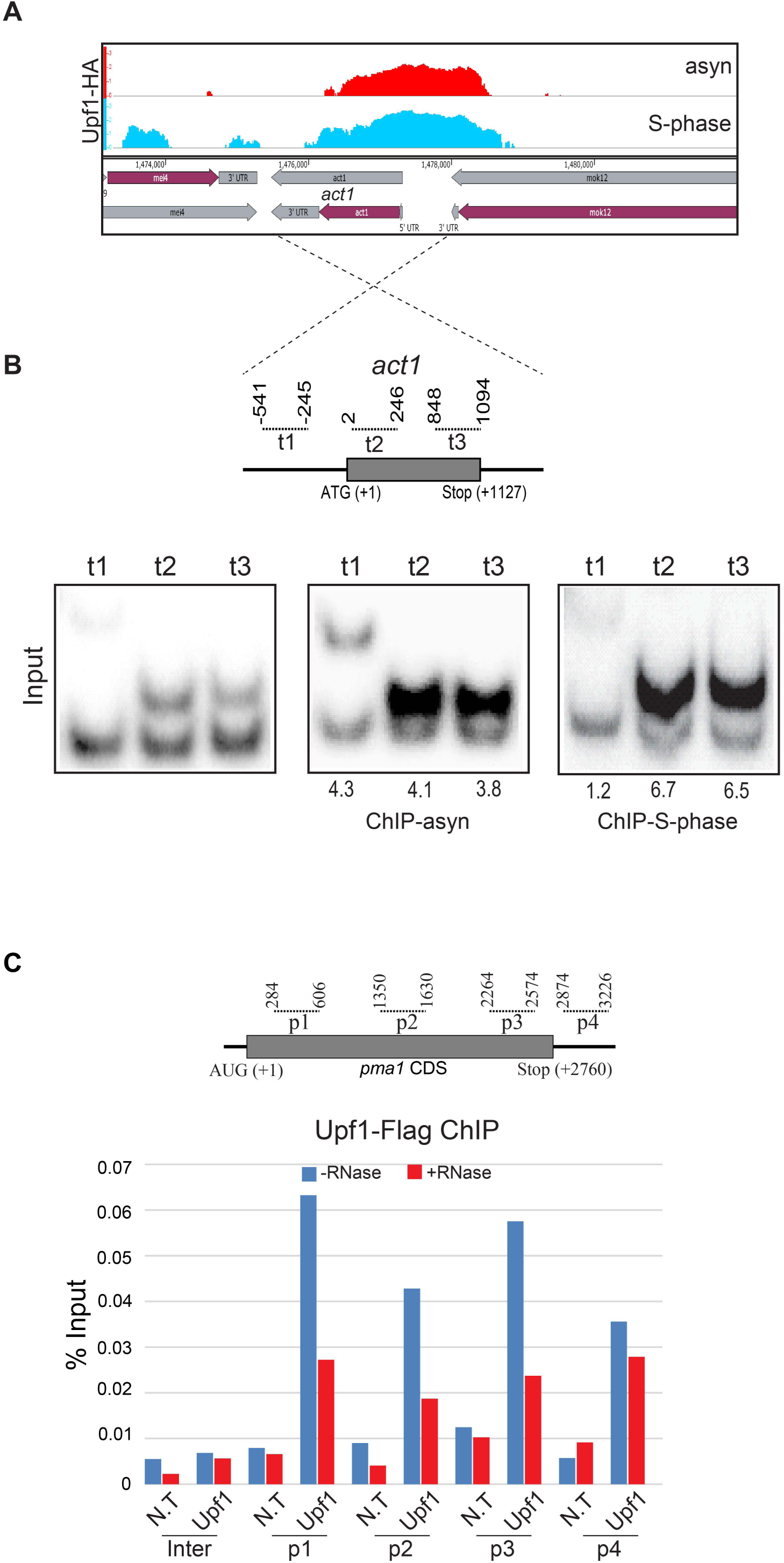
Upf1 association to specific genes is RNase sensitive. **A**) IGB screenshot of ChIP-chip enrichments of Upf1-HA in asynchronous culture (top row) and in S-phase culture (bottom row) over *act1* gene and its flanking region. Genes and genomic features are shown below. **B**) Top panel-schematic diagram of the *act1* gene with CDS sequence (in grey); the PCR amplicons used for the ChIP assay are indicated by the dotted lines above (numbers correspond to the primer positions relative to start codon). Bottom panel-polyacrylamide gels showing radiolabelled PCR products produced by the *act1* specific primer pairs (top bands) and by the pair specific for the intergenic region (bottom bands); using input DNA before ChIP (left panel) and using ChIP-enriched DNA from asynchronous (middle panel) and S-phase culture (right panel). The relative enrichment of *act1* DNA relative to intergenic sequence is expressed as a ratio of the intensity of the same fragments produced with the input DNA. **C**) Independent qPCR quantification of Upf1-Flag ChIP signal on 4 specific regions of *pma1* gene and 1 intergenic control in the absence and presence of RNase. N.T-non-tagged.

**Figure S3.**
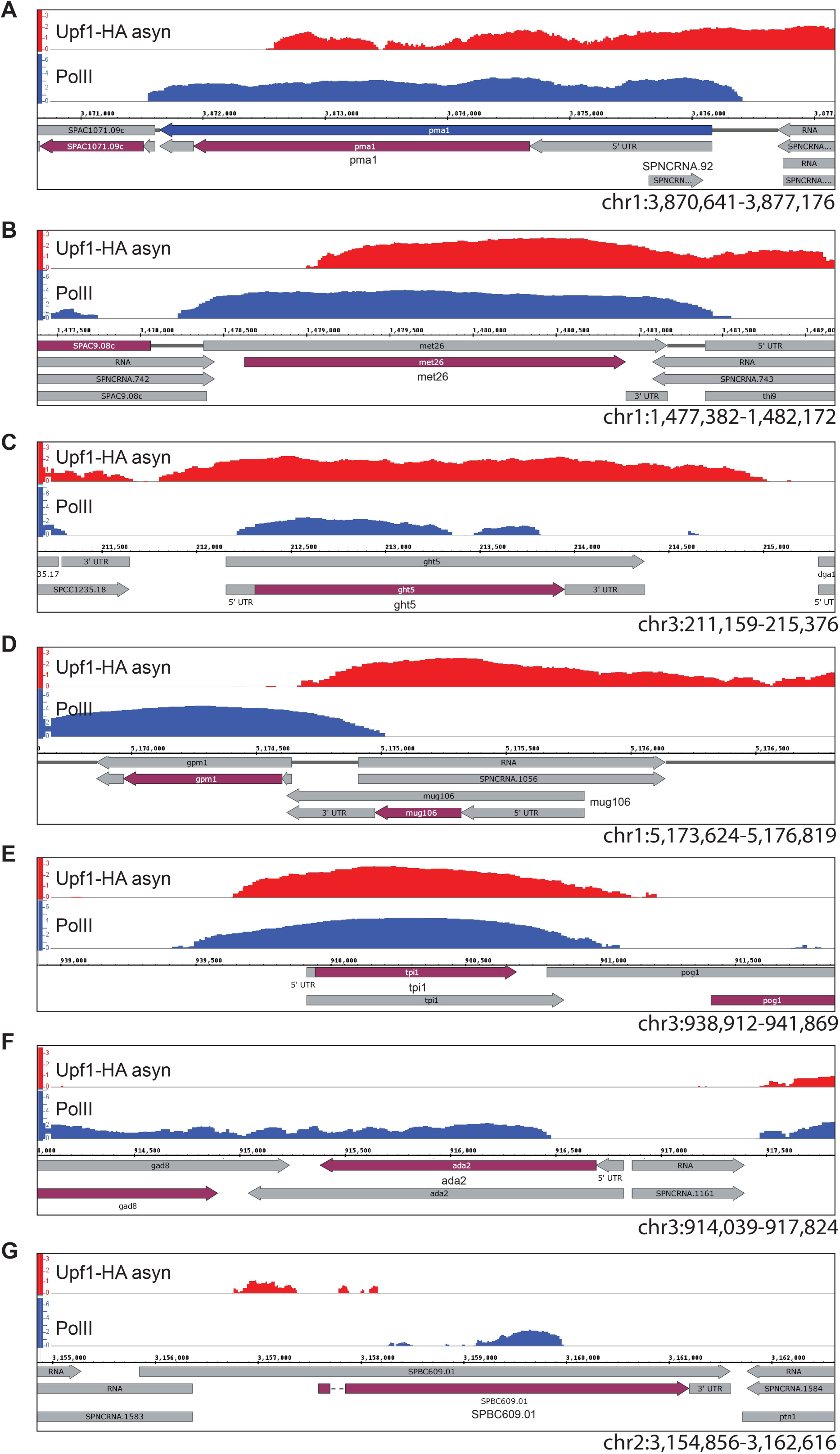
Upf1 binding positively correlates with Pol II association to specific genes. IGB screenshot of ChIP-chip enrichments of Upf1-HA (top row) and Pol II (bottom row) in asynchronous culture over 7 different genes tested in this study. Genes and genomic features are shown below. The 7 genes are **A**) *pma1*, **B**) *met26*, **C**) *ght5*, **D**) *mug-106*, **E**) *tpi1*, **F**) *ada2* and **G**) *SPBC609.01*. The coordinates of genes and flanking genomic regions are mentioned in the bottom right corners.

**Figure S4.**
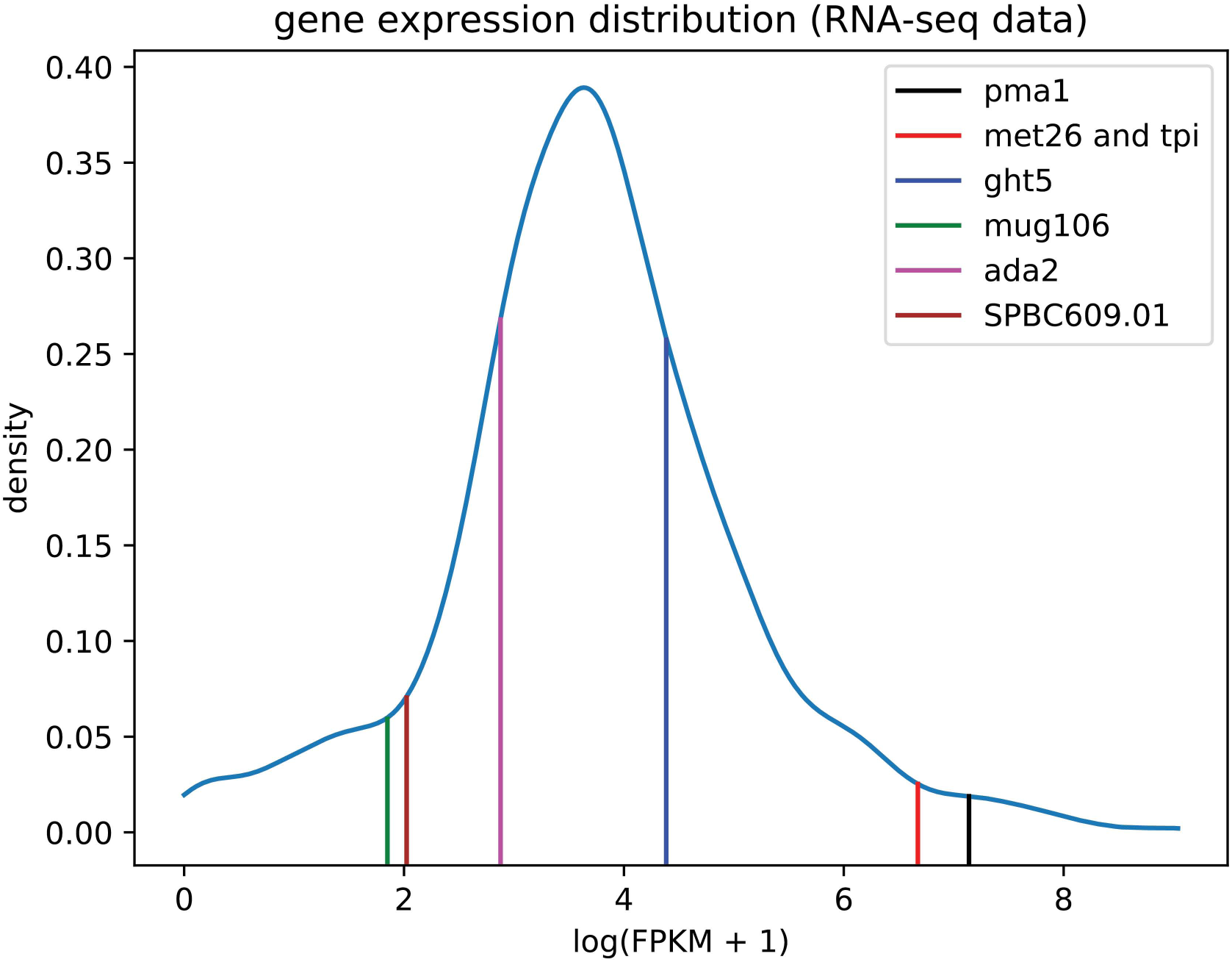
Gene expression distribution pattern of all the *S. pombe* genes. Density plot of log-transformed FPKM+1 values of all the genes expressed in WT strain of *S. pombe*. Expression level of 7 selected genes used in this study to test Upf1, Ser2 Pol II and total Pol II association is indicated on the plot by vertical lines of different colours, as labeled in the inset.

## References

Ajamian, L., Abel, K., Rao, S., Vyboh, K., Garcia-de-Gracia, F., Soto-Rifo, R., Kulozik, A.E., Gehring, N.H., and Mouland, A.J. (2015). HIV-1 Recruits UPF1 but Excludes UPF2 to Promote Nucleocytoplasmic Export of the Genomic RNA. Biomolecules 5, 2808–2839.

Azzalin, C.M., and Lingner, J. (2006). The human RNA surveillance factor UPF1 is required for S phase progression and genome stability. Curr Biol 16, 433–439.

Bahler, J., Wu, J.Q., Longtine, M.S., Shah, N.G., McKenzie, A., 3rd, Steever, A.B., Wach, A., Philippsen, P., and Pringle, J.R. (1998). Heterologous modules for efficient and versatile PCR-based gene targeting in Schizosaccharomyces pombe. Yeast 14, 943–951.

Bhattacharya, A., Czaplinski, K., Trifillis, P., He, F., Jacobson, A., and Peltz, S.W. (2000). Characterization of the biochemical properties of the human Upf1 gene product that is involved in nonsense-mediated mRNA decay. RNA 6, 1226–1235.

Bolger, A.M., Lohse, M., and Usadel, B. (2014). Trimmomatic: a flexible trimmer for Illumina sequence data. Bioinformatics 30, 2114–2120.

Brogna, S., McLeod, T., and Petric, M. (2016). The Meaning of NMD: Translate or Perish. Trends Genet 32, 395–407.

Brogna, S., and Wen, J. (2009). Nonsense-mediated mRNA decay (NMD) mechanisms. Nat Struct Mol Biol 16, 107–113.

Chakrabarti, S., Jayachandran, U., Bonneau, F., Fiorini, F., Basquin, C., Domcke, S., Le Hir, H., and Conti, E. (2011). Molecular mechanisms for the RNA-dependent ATPase activity of Upf1 and its regulation by Upf2. Mol Cell 41, 693–703.

Czaplinski, K., Weng, Y., Hagan, K.W., and Peltz, S.W. (1995). Purification and characterization of the Upf1 protein - a factor involved in translation and messenger RNA degradation. RNA 1, 610–623.

Daulny, A., Mejia-Ramirez, E., Reina, O., Rosado-Lugo, J., Aguilar-Arnal, L., Auer, H., Zaratiegui, M., and Azorin, F. (2016). The fission yeast CENP-B protein Abp1 prevents pervasive transcription of repetitive DNA elements. Biochim Biophys Acta 1859, 1314–1321.

De, S., Varsally, W., Falciani, F., and Brogna, S. (2011). Ribosomal proteins’ association with transcription sites peaks at tRNA genes in Schizosaccharomyces pombe. RNA 17, 1713–1726.

Dennis, G., Jr., Sherman, B.T., Hosack, D.A., Yang, J., Gao, W., Lane, H.C., and Lempicki, R.A. (2003). DAVID: Database for Annotation, Visualization, and Integrated Discovery. Genome biology 4, P3.

El-Brolosy, M.A., Kontarakis, Z., Rossi, A., Kuenne, C., Gunther, S., Fukuda, N., Kikhi, K., Boezio, G.L.M., Takacs, C.M., Lai, S.L., et al. (2019). Genetic compensation triggered by mutant mRNA degradation. Nature 568, 193–197.

Exinger, F., and Lacroute, F. (1992). 6-Azauracil inhibition of GTP biosynthesis in Saccharomyces cerevisiae. Curr Genet 22, 9–11.

Fatscher, T., Boehm, V., and Gehring, N.H. (2015). Mechanism, factors, and physiological role of nonsense-mediated mRNA decay. Cell Mol Life Sci 72, 4523–4544.

Fiorini, F., Bagchi, D., Le Hir, H., and Croquette, V. (2015). Human Upf1 is a highly processive RNA helicase and translocase with RNP remodelling activities. Nature communications 6, 7581.

Fong, N., Saldi, T., Sheridan, R.M., Cortazar, M.A., and Bentley, D.L. (2017). RNA Pol II Dynamics Modulate Co-transcriptional Chromatin Modification, CTD Phosphorylation, and Transcriptional Direction. Mol Cell 66, 546–557 e543.

Franks, T.M., Singh, G., and Lykke-Andersen, J. (2010). Upf1 ATPase-dependent mRNP disassembly is required for completion of nonsense-mediated mRNA decay. Cell 143, 938–950.

Fukuda, M., Asano, S., Nakamura, T., Adachi, M., Yoshida, M., Yanagida, M., and Nishida, E. (1997). CRM1 is responsible for intracellular transport mediated by the nuclear export signal. Nature 390, 308–311.

Gallagher, P.S., Larkin, M., Thillainadesan, G., Dhakshnamoorthy, J., Balachandran, V., Xiao, H., Wellman, C., Chatterjee, R., Wheeler, D., and Grewal, S.I.S. (2018). Iron homeostasis regulates facultative heterochromatin assembly in adaptive genome control. Nat Struct Mol Biol 25, 372–383.

Goetz, A.E., and Wilkinson, M. (2017). Stress and the nonsense-mediated RNA decay pathway. Cell Mol Life Sci 74, 3509–3531.

He, F., and Jacobson, A. (2015). Nonsense-Mediated mRNA Decay: Degradation of Defective Transcripts Is Only Part of the Story. Annu Rev Genet.

Hug, N., Longman, D., and Caceres, J.F. (2016). Mechanism and regulation of the nonsense-mediated decay pathway. Nucleic Acids Res 44, 1483–1495.

Hutten, S., and Kehlenbach, R.H. (2007). CRM1-mediated nuclear export: to the pore and beyond. Trends in cell biology 17, 193–201.

Johnson, W.E., Li, W., Meyer, C.A., Gottardo, R., Carroll, J.S., Brown, M., and Liu, X.S. (2006). Model-based analysis of tiling-arrays for ChIP-chip. Proc Natl Acad Sci U S A 103, 12457–12462.

Karousis, E.D., Nasif, S., and Muhlemann, O. (2016). Nonsense-mediated mRNA decay: novel mechanistic insights and biological impact. Wiley Interdiscip Rev RNA 7, 661–682.

Kim, Y.K., and Maquat, L.E. (2019). UPFront and center in RNA decay: UPF1 in nonsense-mediated mRNA decay and beyond. RNA 25, 407–422.

Kimura, M., Suzuki, H., and Ishihama, A. (2002). Formation of a carboxy-terminal domain phosphatase (Fcp1)/TFIIF/RNA polymerase II (pol II) complex in Schizosaccharomyces pombe involves direct interaction between Fcp1 and the Rpb4 subunit of pol II. Molecular and Cellular Biology 22, 1577–1588.

Knoll, E.R., Zhu, Z.I., Sarkar, D., Landsman, D., and Morse, R.H. (2018). Role of the pre-initiation complex in Mediator recruitment and dynamics. Elife 7.

Langmead, B., and Salzberg, S.L. (2012). Fast gapped-read alignment with Bowtie 2. Nature methods 9, 357–359.

Lykke-Andersen, S., and Jensen, T.H. (2015). Nonsense-mediated mRNA decay: an intricate machinery that shapes transcriptomes. Nat Rev Mol Cell Biol 16, 665–677.

Matsuo, Y., Asakawa, K., Toda, T., and Katayama, S. (2006). A rapid method for protein extraction from fission yeast. Biosci Biotechnol Biochem 70, 1992–1994.

Mendell, J.T., ap Rhys, C.M., and Dietz, H.C. (2002). Separable roles for rent1/hUpf1 in altered splicing and decay of nonsense transcripts. Science 298, 419–422.

Neu-Yilik, G., Raimondeau, E., Eliseev, B., Yeramala, L., Amthor, B., Deniaud, A., Huard, K., Kerschgens, K., Hentze, M.W., Schaffitzel, C., et al. (2017). Dual function of UPF3B in early and late translation termination. EMBO J 36, 2968–2986.

Nicol, J.W., Helt, G.A., Blanchard, S.G., Jr., Raja, A., and Loraine, A.E. (2009). The Integrated Genome Browser: free software for distribution and exploration of genome-scale datasets. Bioinformatics 25, 2730–2731.

Rehwinkel, J., Letunic, I., Raes, J., Bork, P., and Izaurralde, E. (2005). Nonsense-mediated mRNA decay factors act in concert to regulate common mRNA targets. RNA 11, 1530–1544.

Rodriguez-Gabriel, M.A., Watt, S., Bahler, J., and Russell, P. (2006). Upf1, an RNA helicase required for nonsense-mediated mRNA decay, modulates the transcriptional response to oxidative stress in fission yeast. Mol Cell Biol 26, 6347–6356.

Saikrishnan, K., Powell, B., Cook, N.J., Webb, M.R., and Wigley, D.B. (2009). Mechanistic basis of 5’-3’ translocation in SF1B helicases. Cell 137, 849–859.

Shaw, R.J., and Reines, D. (2000). Saccharomyces cerevisiae transcription elongation mutants are defective in PUR5 induction in response to nucleotide depletion. Mol Cell Biol 20, 7427–7437.

Singh, A.K., Choudhury, S.R., De, S., Zhang, J., Kissane, S., Dwivedi, V., Ramanathan, P., Orsini, L., Hebenstreit, D., and Brogna, S. (2019). The RNA helicase UPF1 associates with mRNAs co-transcriptionally and is required for the release of mRNAs from transcription sites. Elife 2019;8:e41444.

Singh, A.K., Zhang, J., Hebenstreit, D., and Brogna, S. (2020). Evidence of slightly increased Pol II pausing in UPF1-depleted Drosophila melanogaster cells. MicroPubl Biol 2020.

Singleton, M.R., Dillingham, M.S., and Wigley, D.B. (2007). Structure and mechanism of helicases and nucleic acid translocases. Annu Rev Biochem 76, 23–50.

Tusher, V.G., Tibshirani, R., and Chu, G. (2001). Significance analysis of microarrays applied to the ionizing radiation response. Proc Natl Acad Sci U S A 98, 5116–5121.

Varsally, W., and Brogna, S. (2012). UPF1 involvement in nuclear functions. Biochem Soc Trans 40, 778–783.

Wen, J., and Brogna, S. (2010). Splicing-dependent NMD does not require the EJC in Schizosaccharomyces pombe. The Embo Journal 29, 1537–1551.

Wen, J., He, M., Petric, M., Marzi, L., Wang, J., Piechocki, K., McLeod, T., Singh, A.K., Dwivedi, V., and Brogna, S. (2020). An intron proximal to a PTC enhances NMD in Saccharomyces cerevisiae. bioRxiv, 149245.

Wilhelm, B.T., Marguerat, S., Watt, S., Schubert, F., Wood, V., Goodhead, I., Penkett, C.J., Rogers, J., and Bahler, J. (2008). Dynamic repertoire of a eukaryotic transcriptome surveyed at single-nucleotide resolution. Nature 453, 1239–1243.

Yague-Sanz, C., Vanrobaeys, Y., Fernandez, R., Duval, M., Larochelle, M., Beaudoin, J., Berro, J., Labbe, S., Jacques, P.E., and Bachand, F. (2020). Nutrient-dependent control of RNA polymerase II elongation rate regulates specific gene expression programs by alternative polyadenylation. Genes Dev 34, 883–897.

Zhou, H., Liu, Q., Shi, T., Yu, Y., and Lu, H. (2015). Genome-wide screen of fission yeast mutants for sensitivity to 6-azauracil, an inhibitor of transcriptional elongation. Yeast 32, 643–655.

