## Supplemental files for "Genome-wide chromosomal association of Upf1 is linked to Pol II transcription in *Schizosaccharomyces pombe*"

### Supplementary files description

The two supplementary html files show plots of the normalised single-base resolution coverage (Materials and Methods) of the Pol II ChIP-seq signal for two different sets of genes. Two plots are shown for each gene. The first plot compares the coverage depth from 1000bp upstream of the TSS to 1000bp downstream of the TES in wild-type (WT, blue line) and *upf1*Δ (termed KO, red line) Upf1 cells for that particular gene. This is referred to as the "normalised ChIP-seq signal" in the y-axis. The second plots show the difference between the WT and *upf1*Δ signals, so that more negative values indicate increased Pol II occupancy when Upf1 is deleted. The vertical black line on the left side of each plot marks the TSS and the vertical black line on the right marks the position of the TES. The gene that each plot corresponds to is indicated by the ID in the plot title.

Supplementary File 1

The genes plotted in supplementary file 1 are a selection of those found to be strongly associated with Upf1 in WT cells, which also exhibit increased Pol II occupancy throughout the gene body, at the TES and/or downstream of the TES in *upf1*Δ (termed KO) cells, as demonstrated by the plots. This selection does not include the genes that are misregulated in *upf1*Δ cells, which are instead shown separately in Supplementary File 2 below.

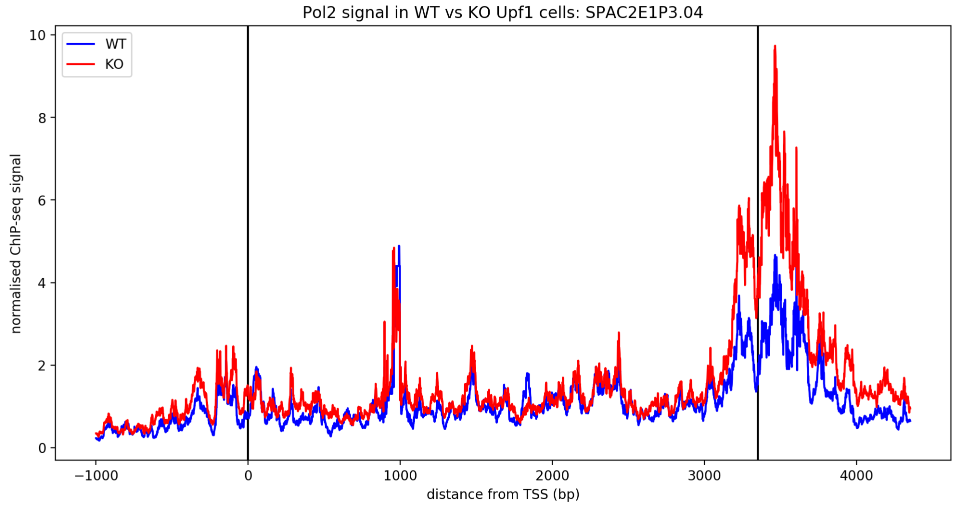
Pol2 signal in WT vs KO Upf1 cells: SPAC2E1P3.04
WT
KO
normalised ChIP-seq signal
distance from TSS (bp)
0
2
4
6
8
10
−1000
0
1000
2000
3000
4000

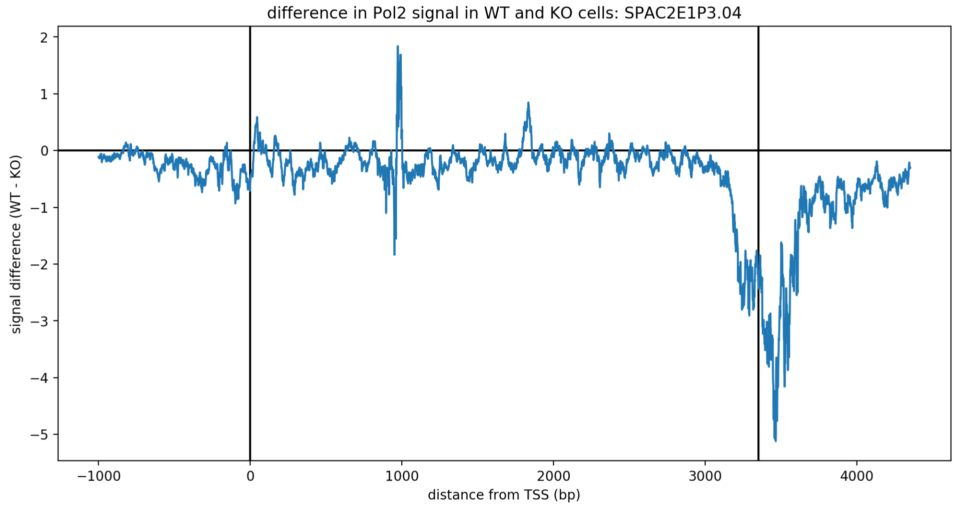
difference in Pol2 signal in WT and KO cells: SPAC2E1P3.04
signal difference (WT - KO)
distance from TSS (bp)
2
1
0
−1
−2
−3
−4
−5
−1000
0
1000
2000
3000
4000

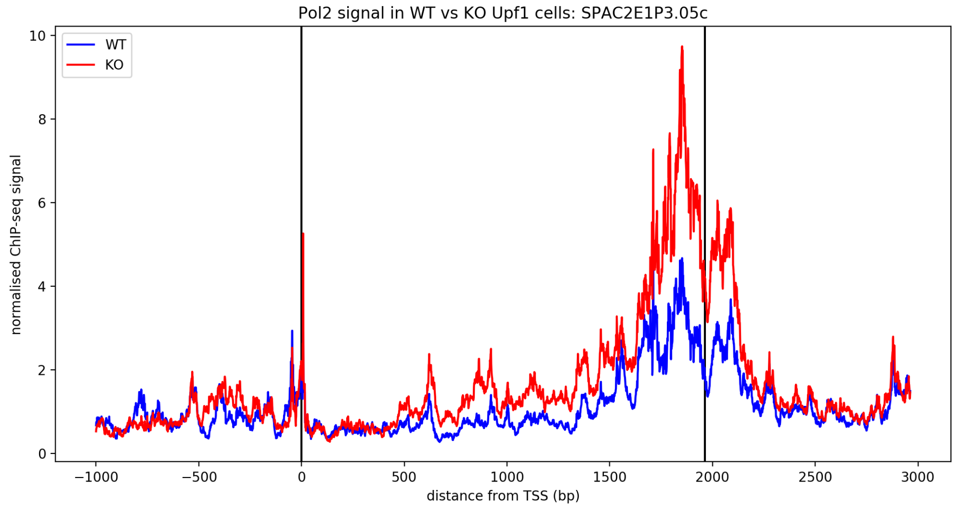
Pol2 signal in WT vs KO Upf1 cells: SPAC2E1P3.05c
WT
KO
normalised ChIP-seq signal
distance from TSS (bp)
0
2
4
6
8
10
−1000
−500
0
500
1000
1500
2000
2500
3000

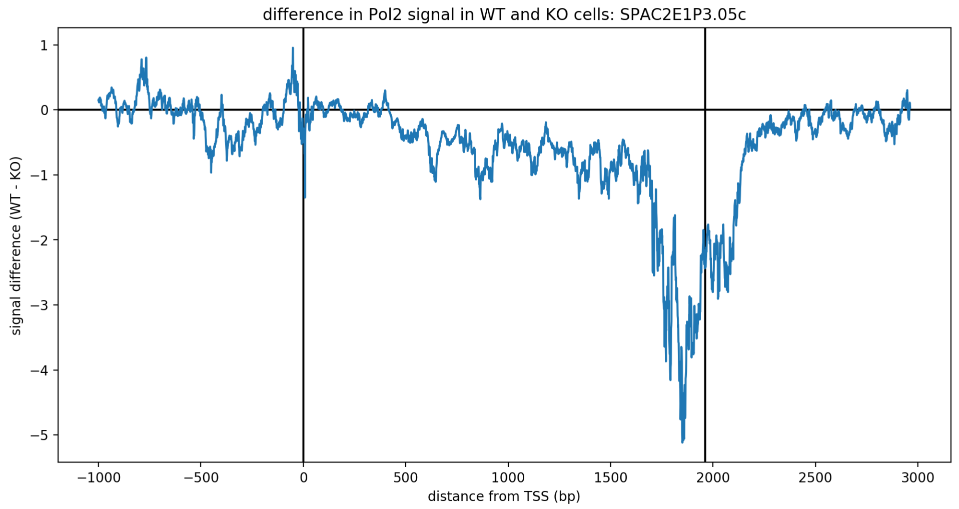
difference in Pol2 signal in WT and KO cells: SPAC2E1P3.05c
signal difference (WT - KO)
distance from TSS (bp)

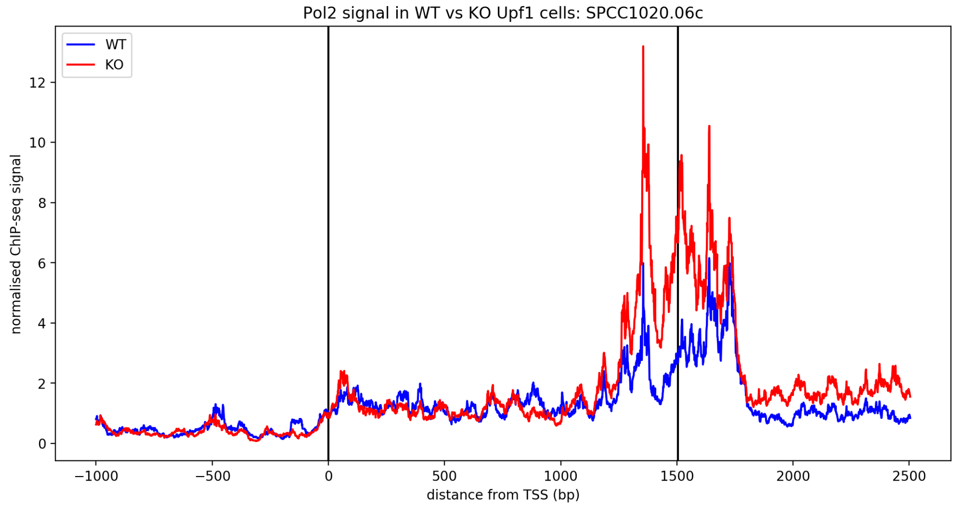
Pol2 signal in WT vs KO Upf1 cells: SPCC1020.06c
WT
KO
normalised ChIP-seq signal
distance from TSS (bp)

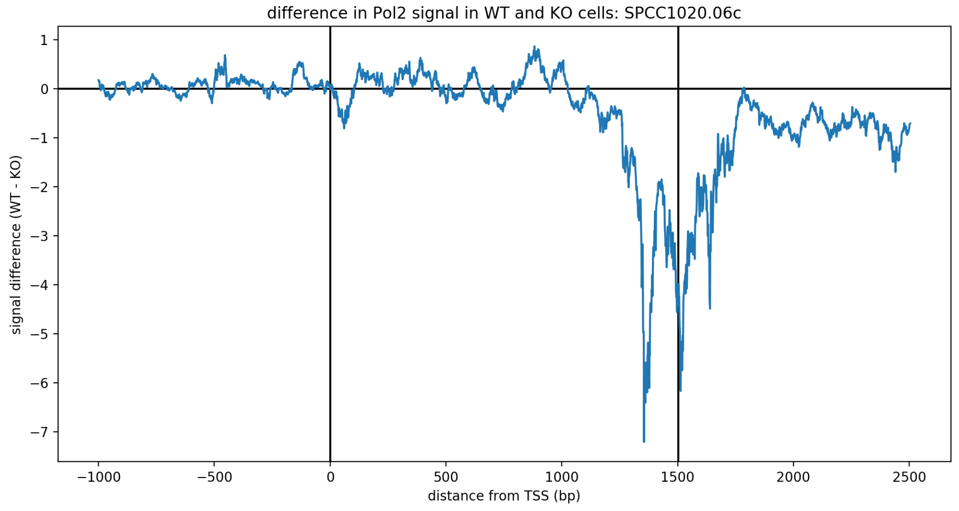
difference in Pol2 signal in WT and KO cells: SPCC1020.06c
signal difference (WT - KO)
distance from TSS (bp)

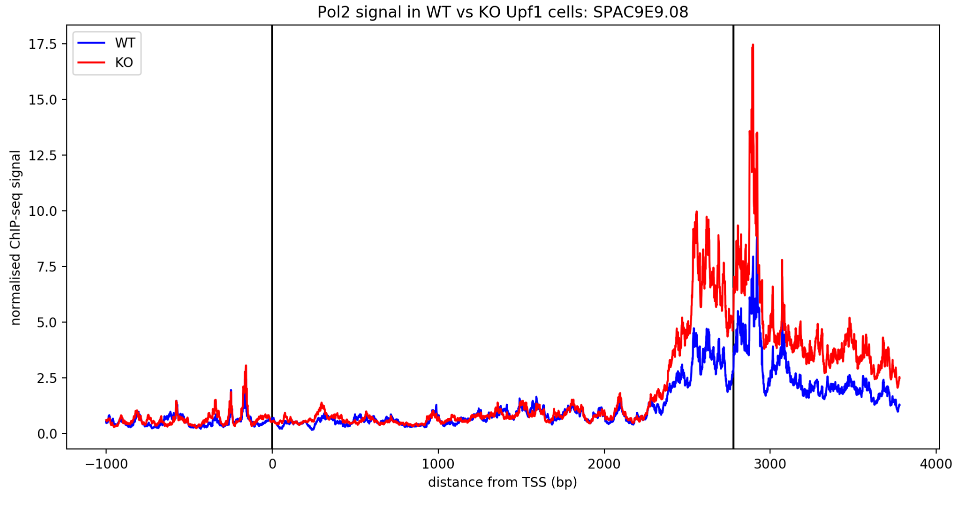
Pol2 signal in WT vs KO Upf1 cells: SPAC9E9.08
WT
KO
normalised ChIP-seq signal
distance from TSS (bp)

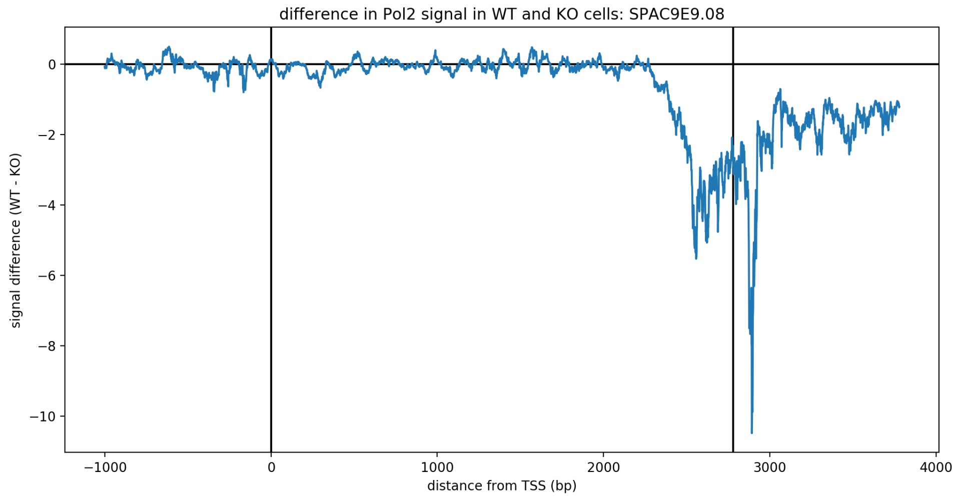
difference in Pol2 signal in WT and KO cells: SPAC9E9.08
signal difference (WT - KO)
0
−2
−4
−6
−8
−10
−1000
0
1000
2000
3000
4000
distance from TSS (bp)

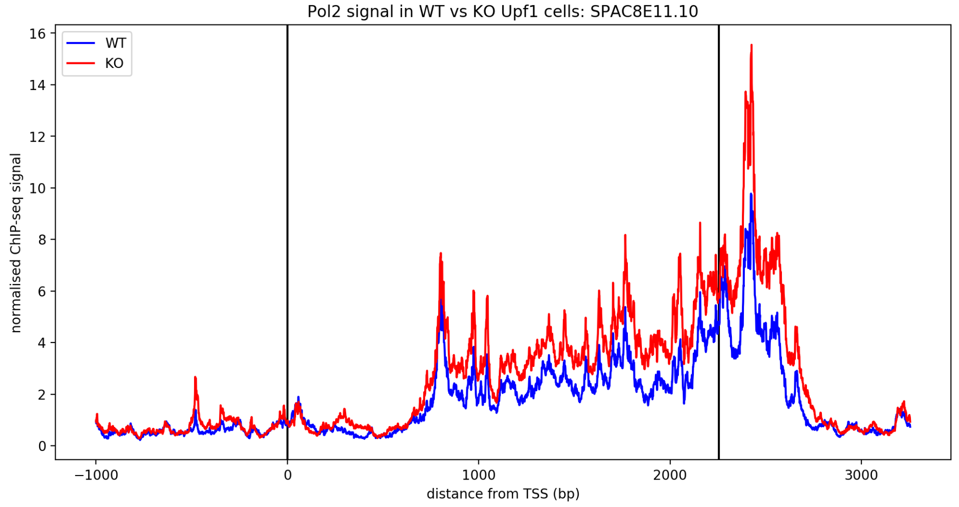
Pol2 signal in WT vs KO Upf1 cells: SPAC8E11.10
WT
KO
normalised ChIP-seq signal
16
14
12
10
8
6
4
2
0
−1000
0
1000
2000
3000
distance from TSS (bp)

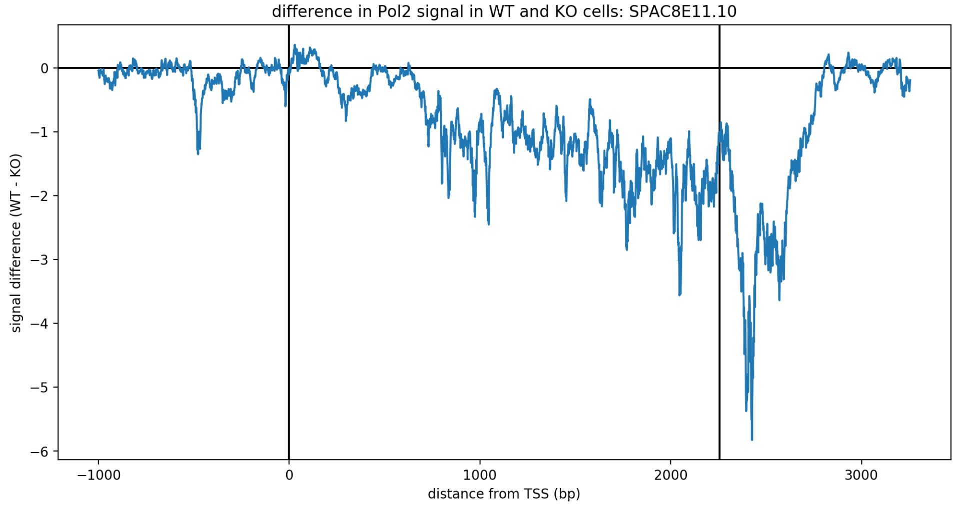
difference in Pol2 signal in WT and KO cells: SPAC8E11.10
signal difference (WT - KO)
0
−1
−2
−3
−4
−5
−6
−1000
0
1000
2000
3000
distance from TSS (bp)

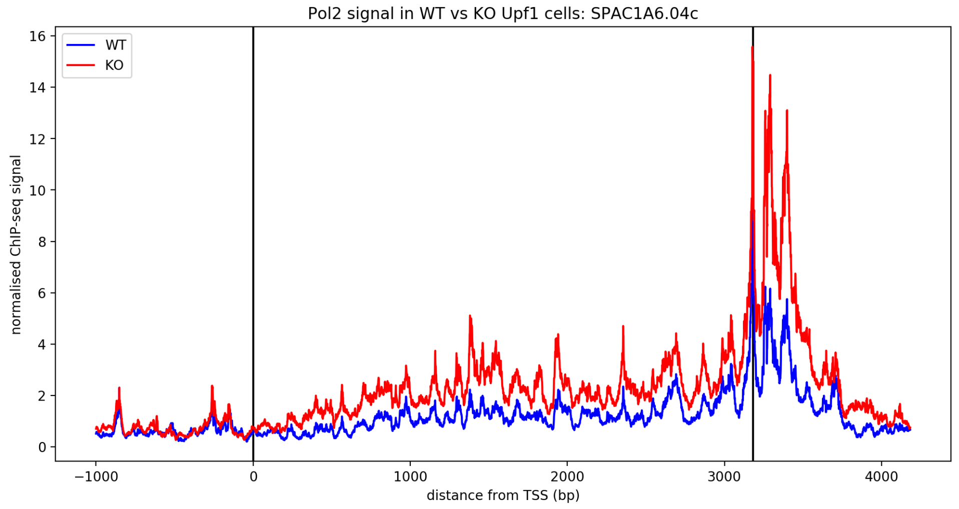
Pol2 signal in WT vs KO Upf1 cells: SPAC1A6.04c
WT
KO
normalised ChIP-seq signal
16
14
12
10
8
6
4
2
0
−1000
0
1000
2000
3000
4000
distance from TSS (bp)

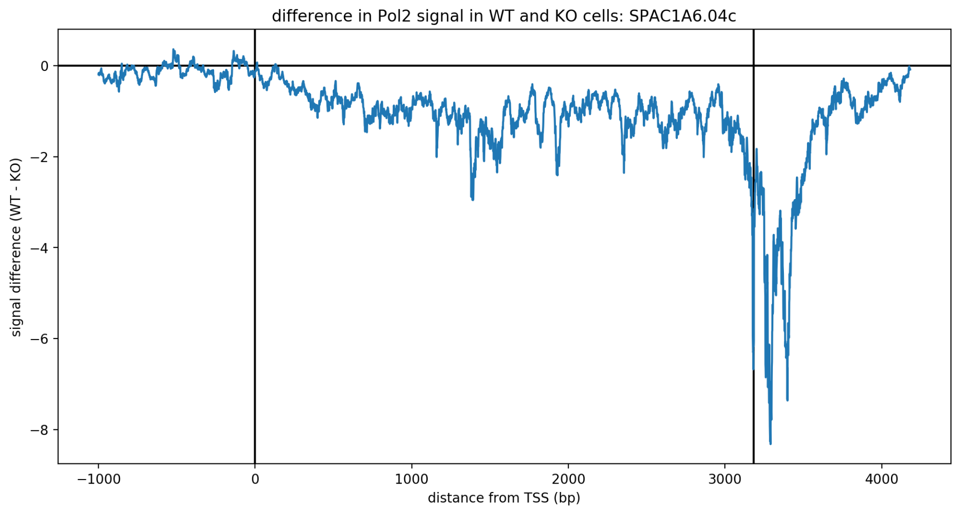
difference in Pol2 signal in WT and KO cells: SPAC1A6.04c
signal difference (WT - KO)
0
−2
−4
−6
−8
−1000
0
1000
2000
3000
4000
distance from TSS (bp)

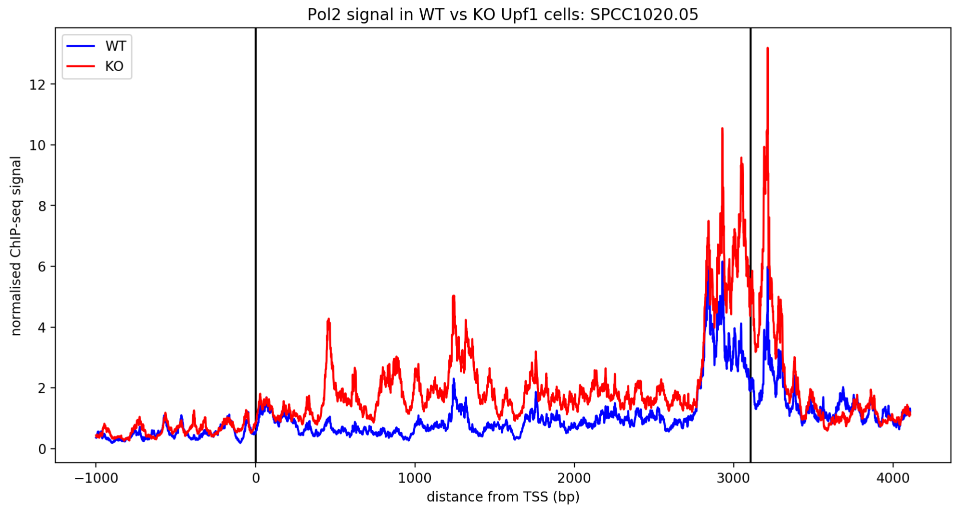
Pol2 signal in WT vs KO Upf1 cells: SPCC1020.05
WT
KO
normalised ChIP-seq signal
12
10
8
6
4
2
0
−1000
0
1000
2000
3000
4000
distance from TSS (bp)

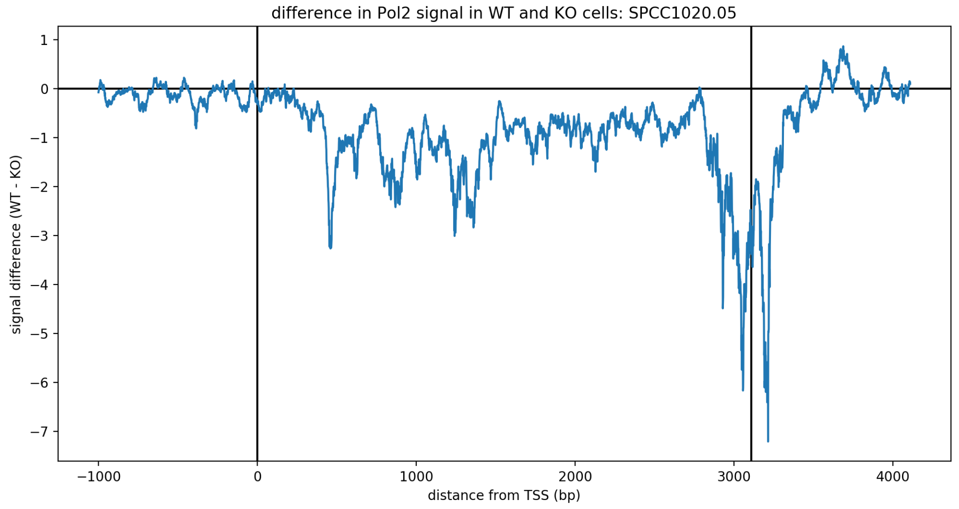
difference in Pol2 signal in WT and KO cells: SPCC1020.05
signal difference (WT - KO)
1
0
−1
−2
−3
−4
−5
−6
−7
−1000
0
1000
2000
3000
4000
distance from TSS (bp)

Pol2 signal in WT vs KO Upf1 cells: SPBC19C2.07

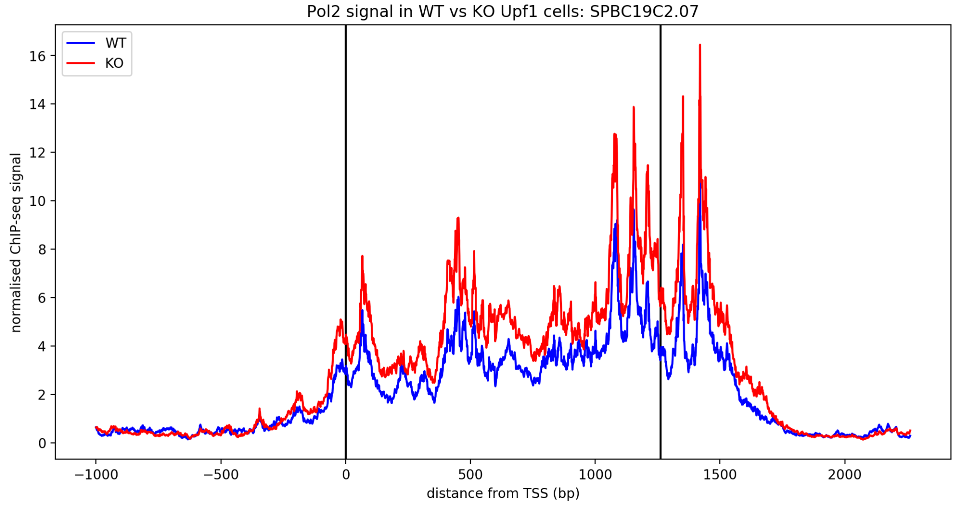


difference in Pol2 signal in WT and KO cells: SPBC19C2.07

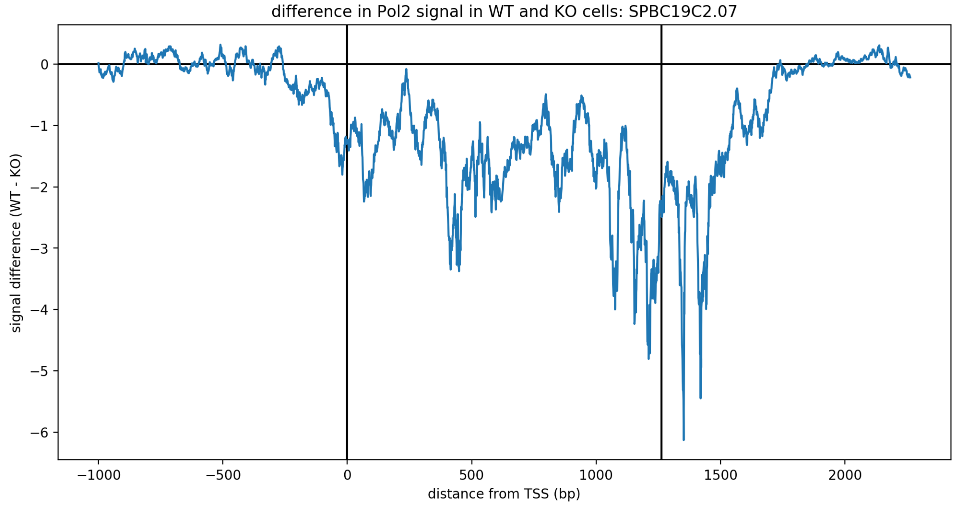


Pol2 signal in WT vs KO Upf1 cells: SPBC887.15c

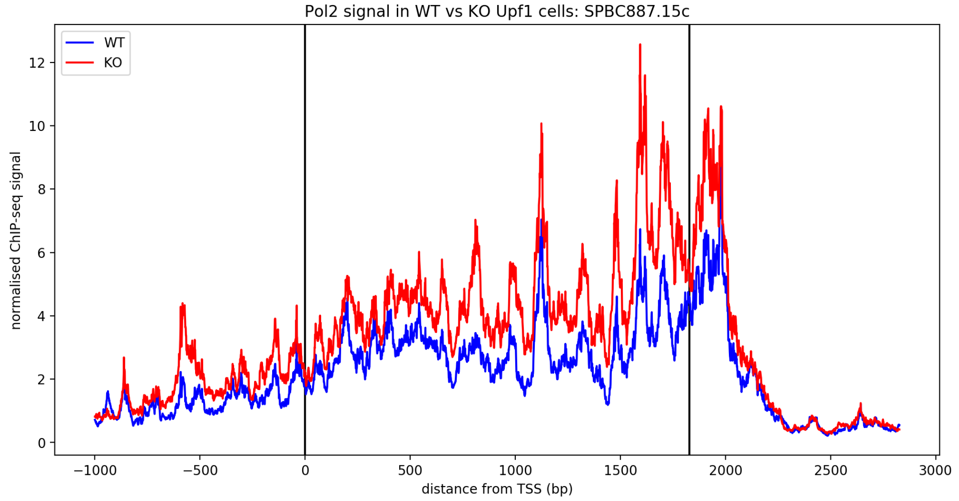


difference in Pol2 signal in WT and KO cells: SPBC887.15c

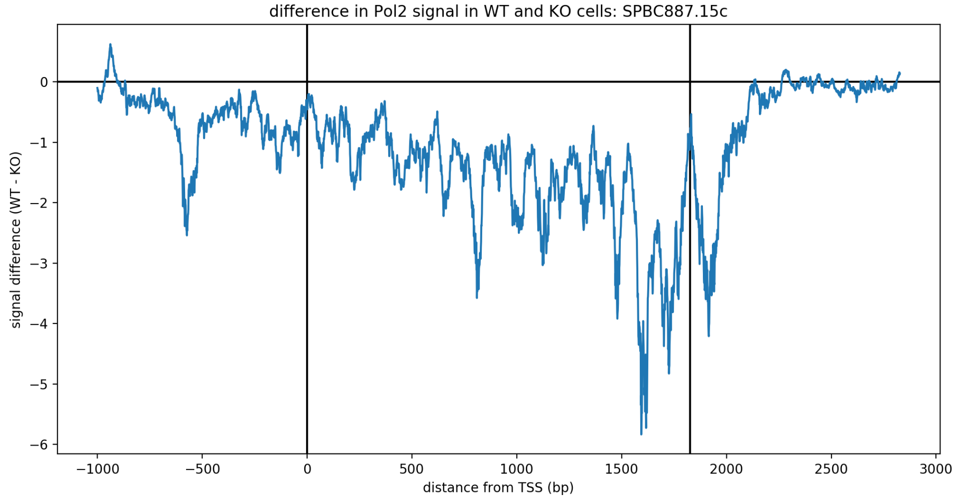

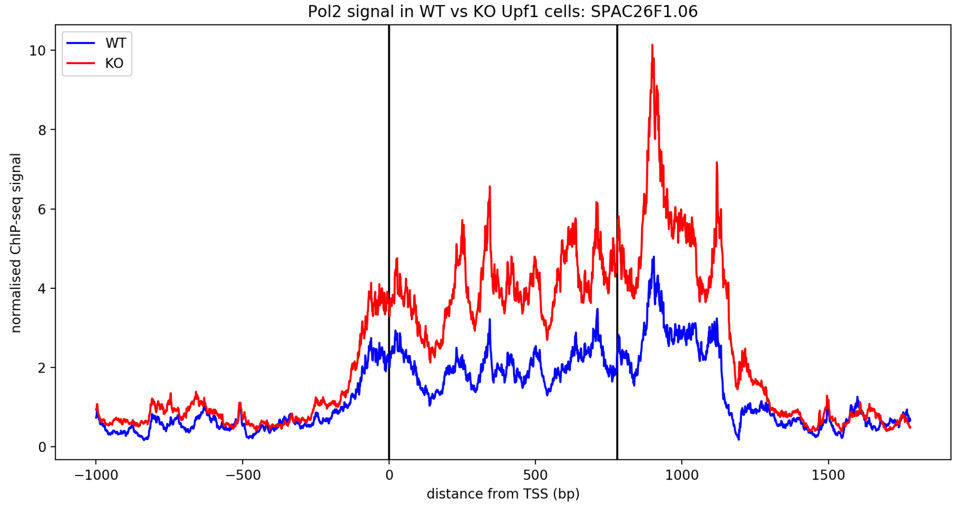
Pol2 signal in WT vs KO Upf1 cells: SPAC26F1.06
WT
KO
normalised ChIP-seq signal
distance from TSS (bp)
0
2
4
6
8
10
−1000
−500
0
500
1000
1500

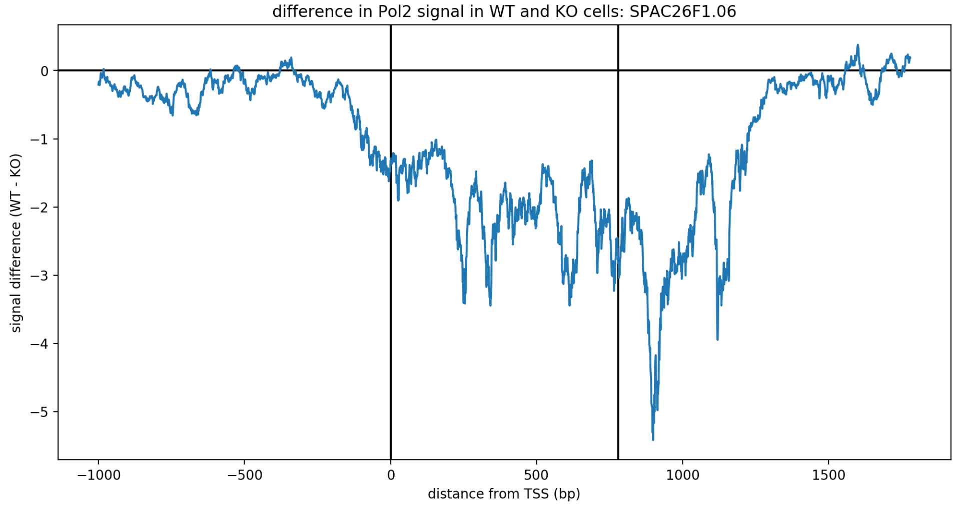
difference in Pol2 signal in WT and KO cells: SPAC26F1.06
signal difference (WT - KO)
distance from TSS (bp)
0
−1
−2
−3
−4
−5
−1000
−500
0
500
1000
1500

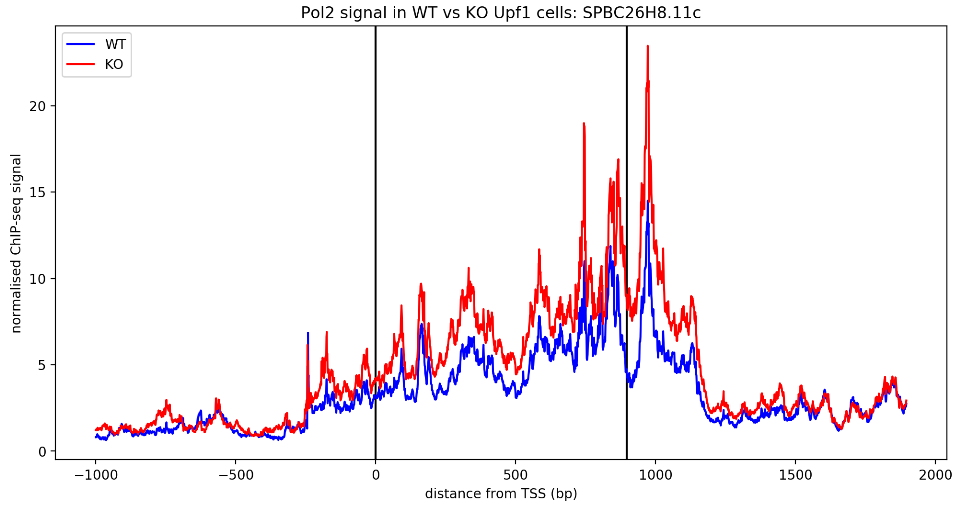
Pol2 signal in WT vs KO Upf1 cells: SPBC26H8.11c
WT
KO
normalised ChIP-seq signal
distance from TSS (bp)
0
5
10
15
20
−1000
−500
0
500
1000
1500
2000

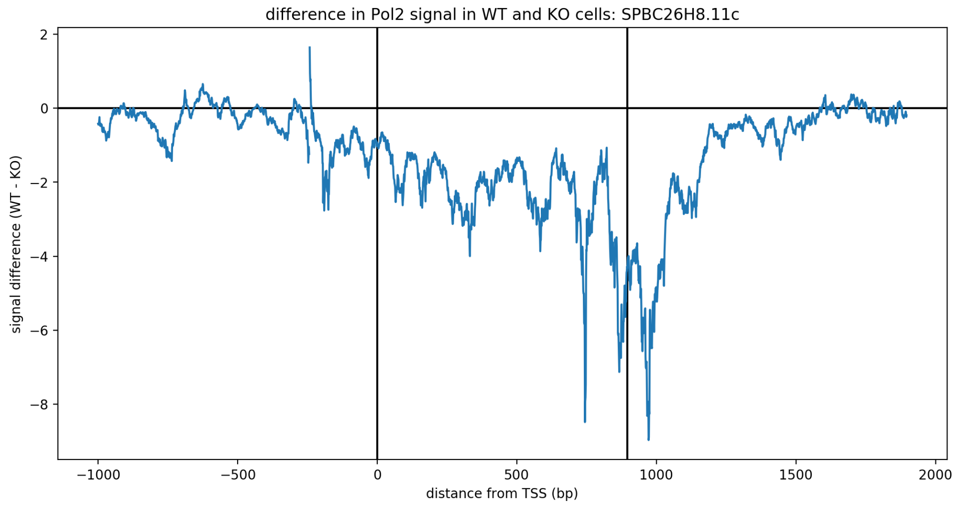
difference in Pol2 signal in WT and KO cells: SPBC26H8.11c
signal difference (WT - KO)
distance from TSS (bp)

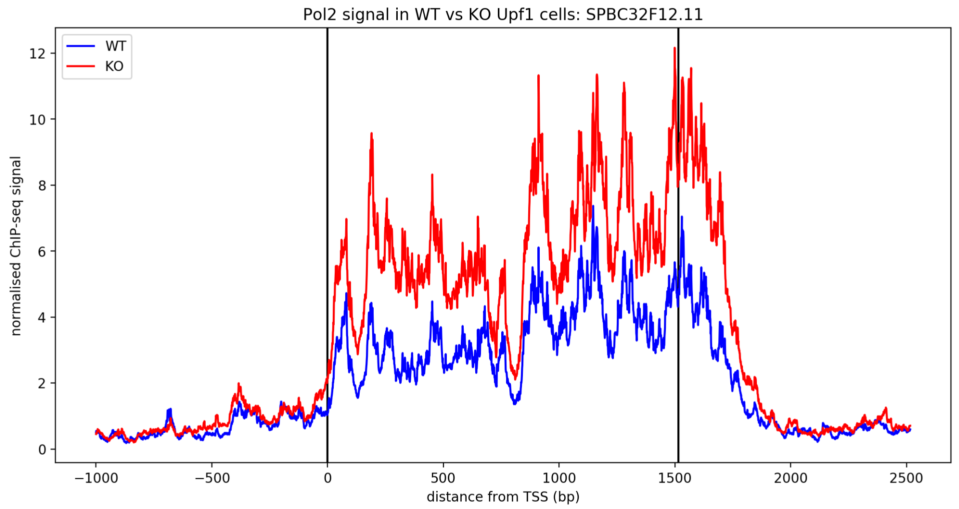
Pol2 signal in WT vs KO Upf1 cells: SPBC32F12.11
WT
KO
normalised ChIP-seq signal
distance from TSS (bp)

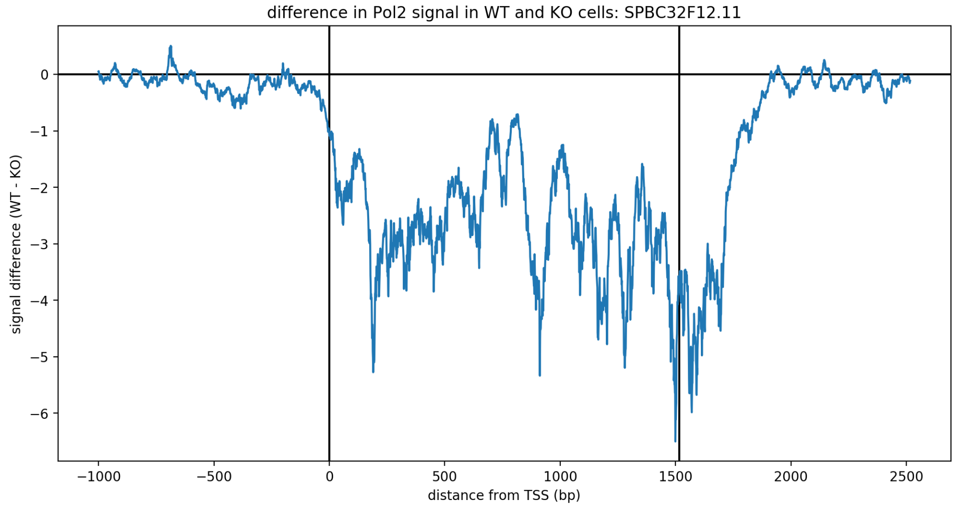
difference in Pol2 signal in WT and KO cells: SPBC32F12.11
signal difference (WT - KO)
distance from TSS (bp)

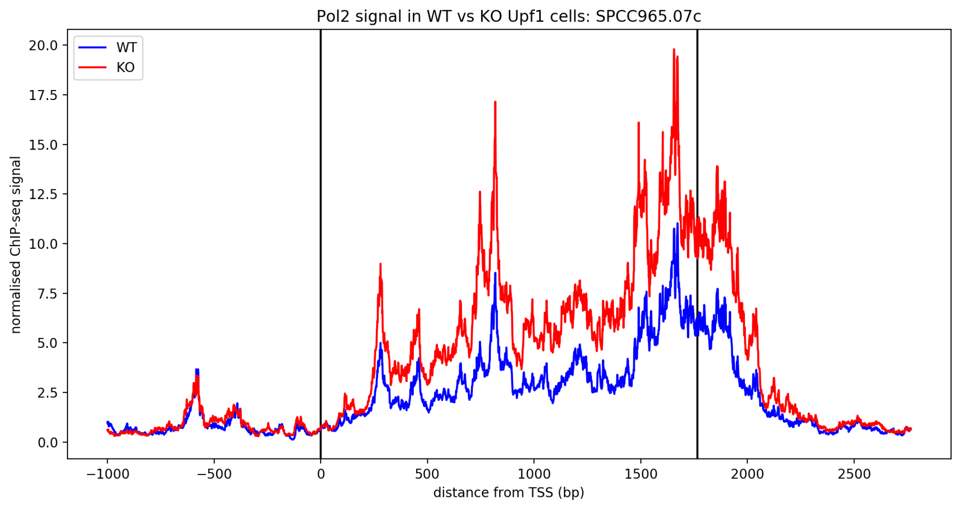
Pol2 signal in WT vs KO Upf1 cells: SPCC965.07c
WT
KO
normalised ChIP-seq signal
distance from TSS (bp)

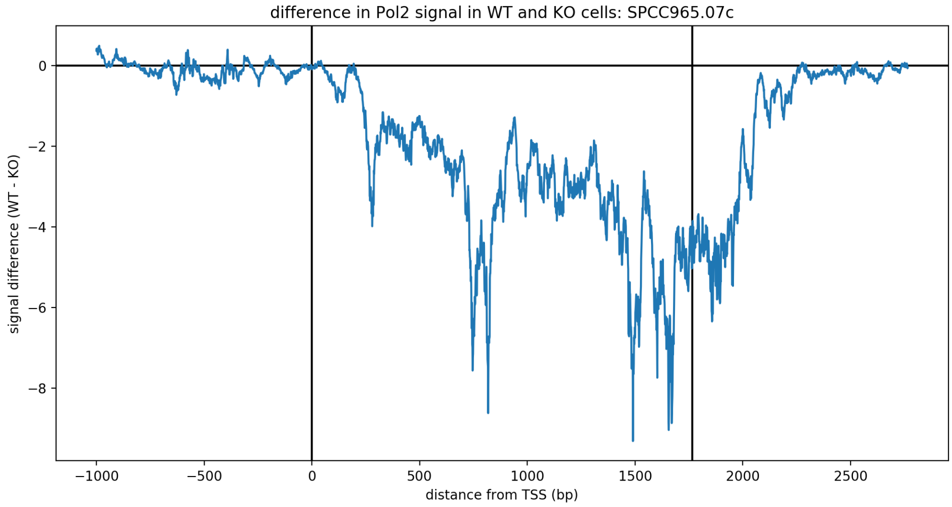
difference in Pol2 signal in WT and KO cells: SPCC965.07c
signal difference (WT - KO)
distance from TSS (bp)
−1000
−500
0
500
1000
1500
2000
2500
0
−2
−4
−6
−8

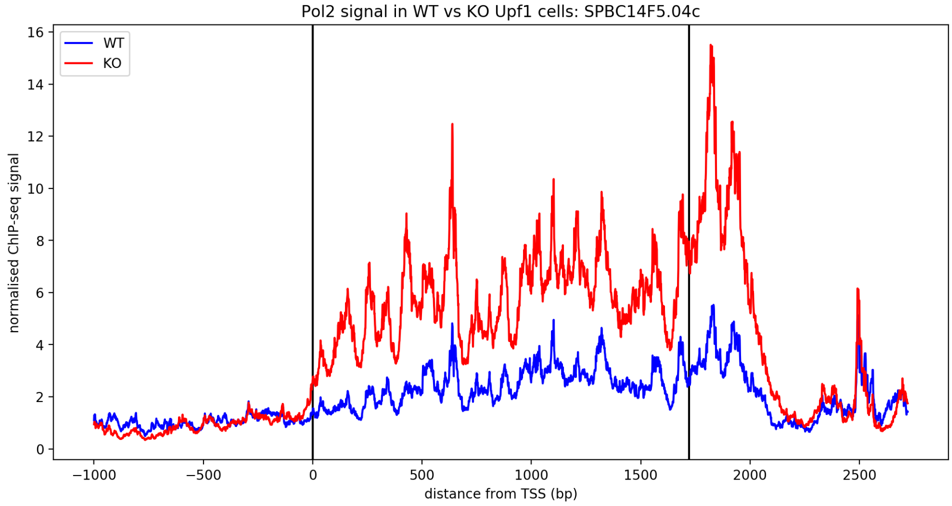
Pol2 signal in WT vs KO Upf1 cells: SPBC14F5.04c
WT
KO
normalised ChIP-seq signal
distance from TSS (bp)
−1000
−500
0
500
1000
1500
2000
2500
16
14
12
10
8
6
4
2
0

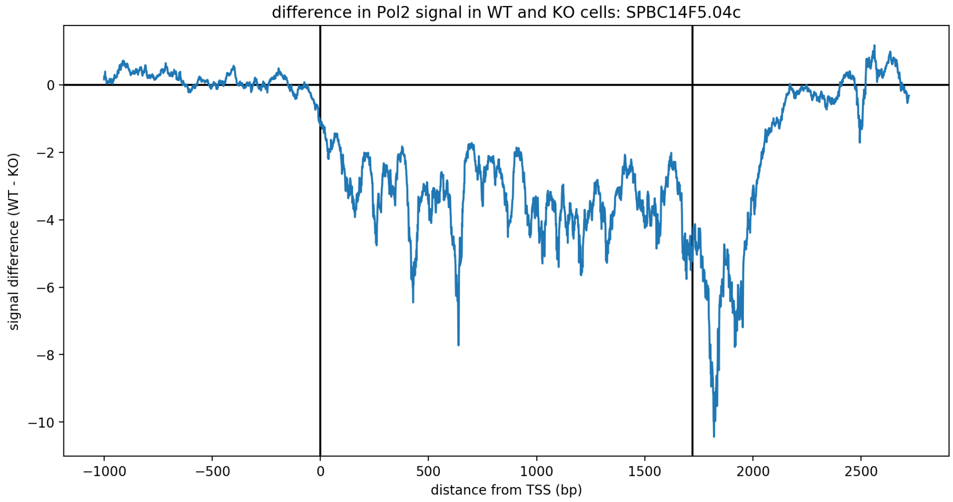
difference in Pol2 signal in WT and KO cells: SPBC14F5.04c
signal difference (WT - KO)
distance from TSS (bp)
−1000
−500
0
500
1000
1500
2000
2500
0
−2
−4
−6
−8
−10

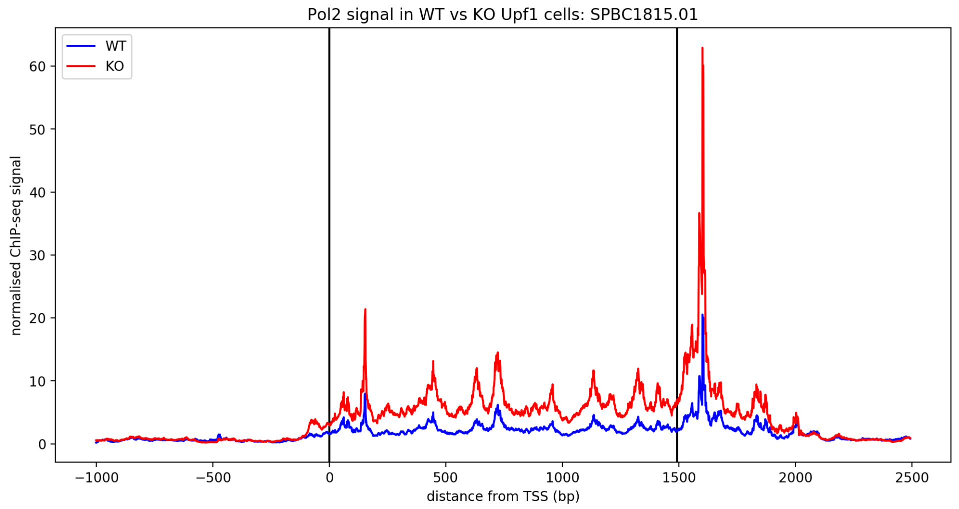

Pol2 signal in WT vs KO Upf1 cells: SPBC1815.01
WT
KO
normalised ChIP-seq signal
distance from TSS (bp)


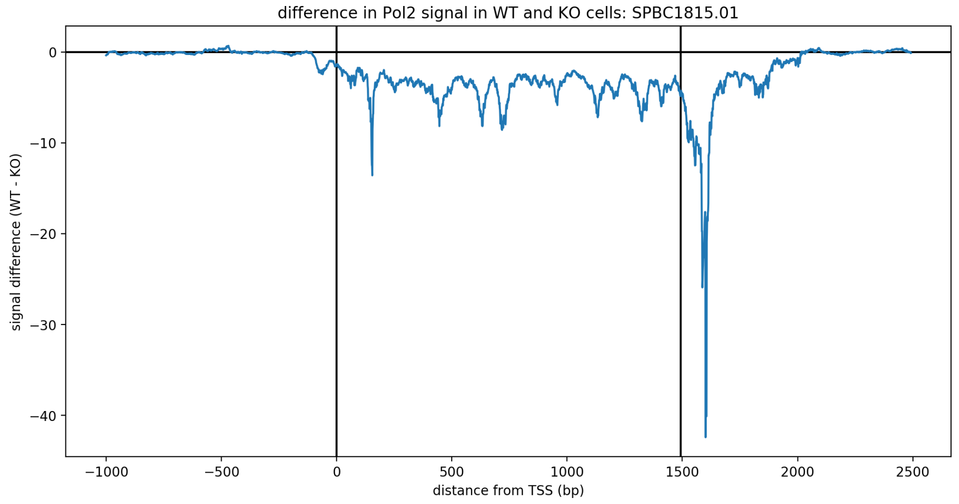

difference in Pol2 signal in WT and KO cells: SPBC1815.01
signal difference (WT - KO)
distance from TSS (bp)


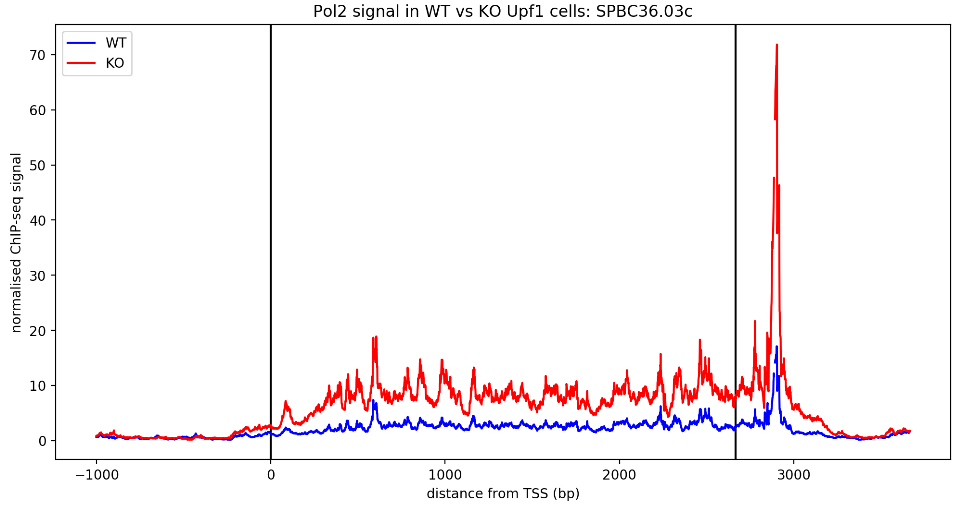

Pol2 signal in WT vs KO Upf1 cells: SPBC36.03c
WT
KO
normalised ChIP-seq signal
distance from TSS (bp)


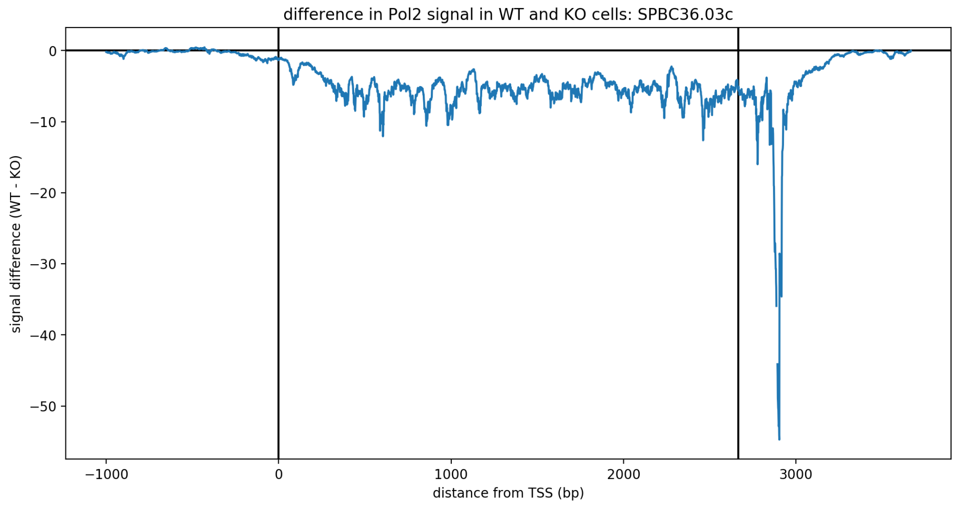

difference in Pol2 signal in WT and KO cells: SPBC36.03c
signal difference (WT - KO)
distance from TSS (bp)

Supplementary File 2

The genes plotted in Supplementary File 2 are a selection of those found to be both strongly associated with Upf1 in WT cells and misregulated in *upf1*Δ (termed KO) cells, which also exhibit increased Pol II occupancy throughout the gene body, at the TES and/or downstream of the TES in *upf1*Δ cells.

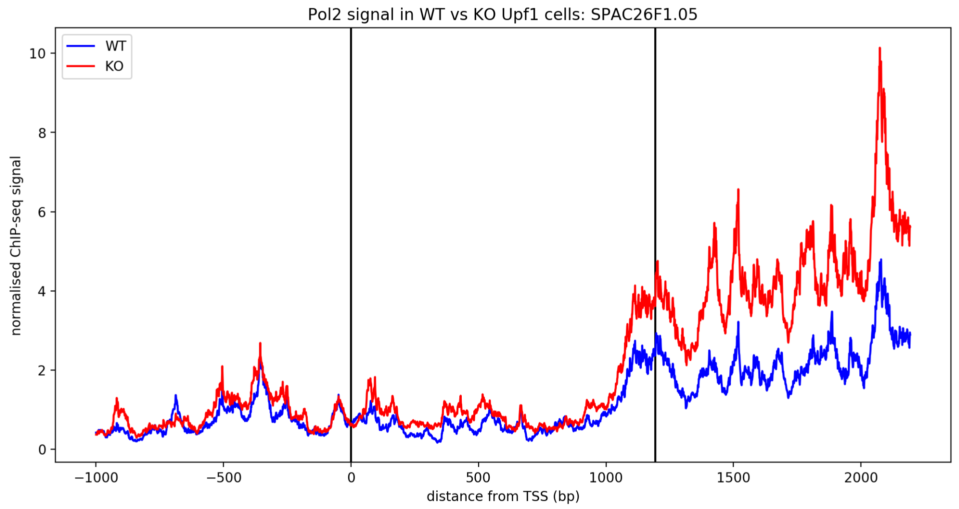
Pol2 signal in WT vs KO Upf1 cells: SPAC26F1.05
WT
KO
normalised ChIP-seq signal
distance from TSS (bp)
0
2
4
6
8
10
−1000
−500
0
500
1000
1500
2000

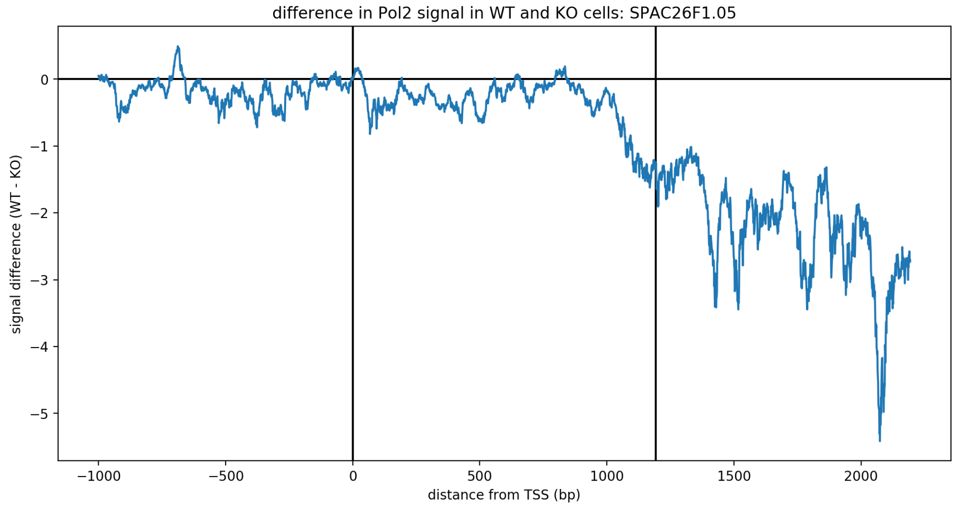
difference in Pol2 signal in WT and KO cells: SPAC26F1.05
signal difference (WT - KO)
distance from TSS (bp)
0
−1
−2
−3
−4
−5
−1000
−500
0
500
1000
1500
2000

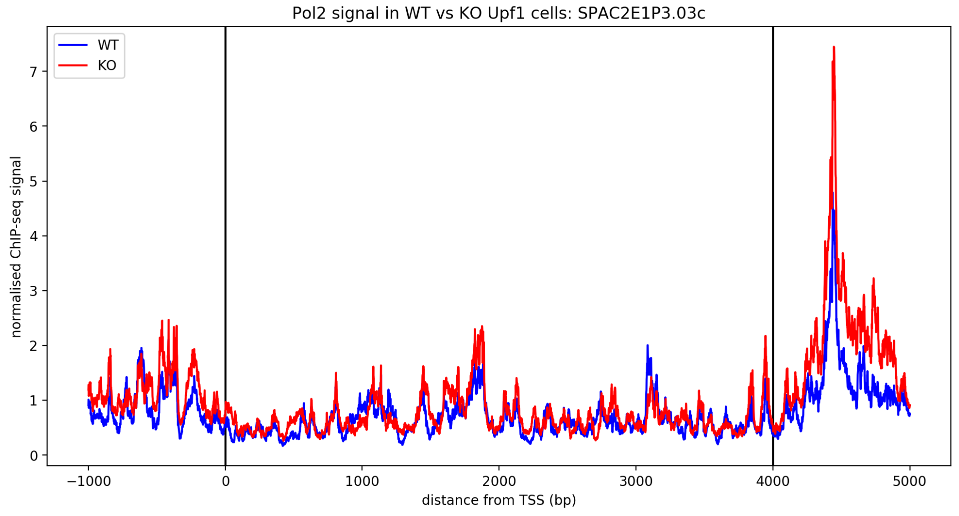
Pol2 signal in WT vs KO Upf1 cells: SPAC2E1P3.03c
WT
KO
normalised ChIP-seq signal
distance from TSS (bp)
0
1
2
3
4
5
6
7
−1000
0
1000
2000
3000
4000
5000

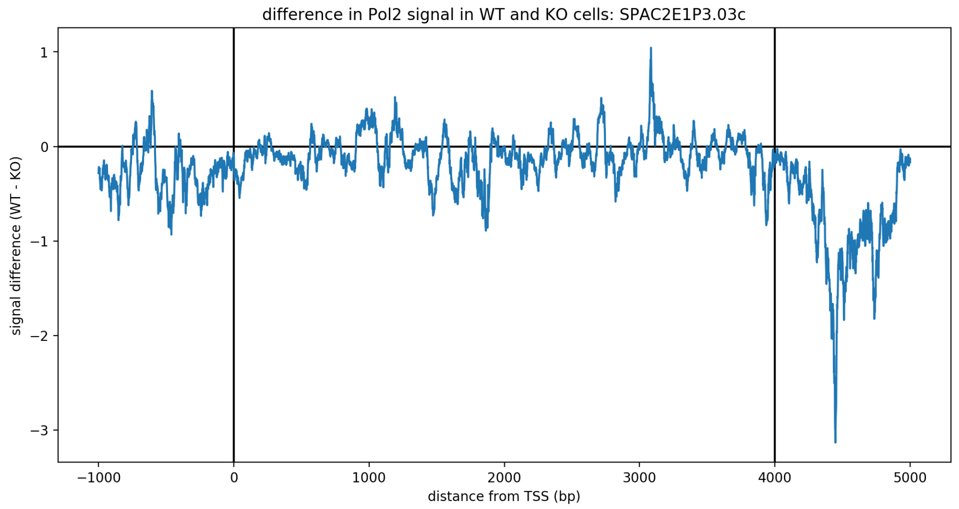

difference in Pol2 signal in WT and KO cells: SPAC2E1P3.03c
signal difference (WT - KO)
distance from TSS (bp)


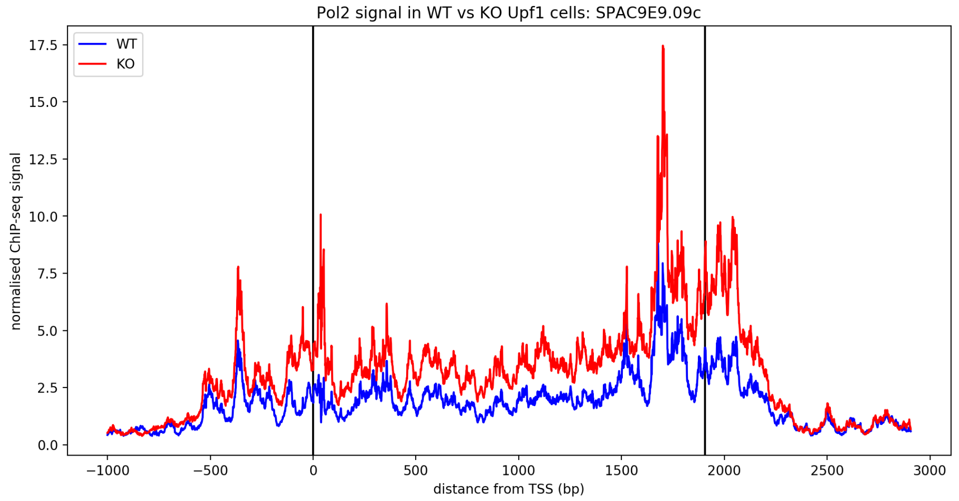

Pol2 signal in WT vs KO Upf1 cells: SPAC9E9.09c
WT
KO
normalised ChIP-seq signal
distance from TSS (bp)


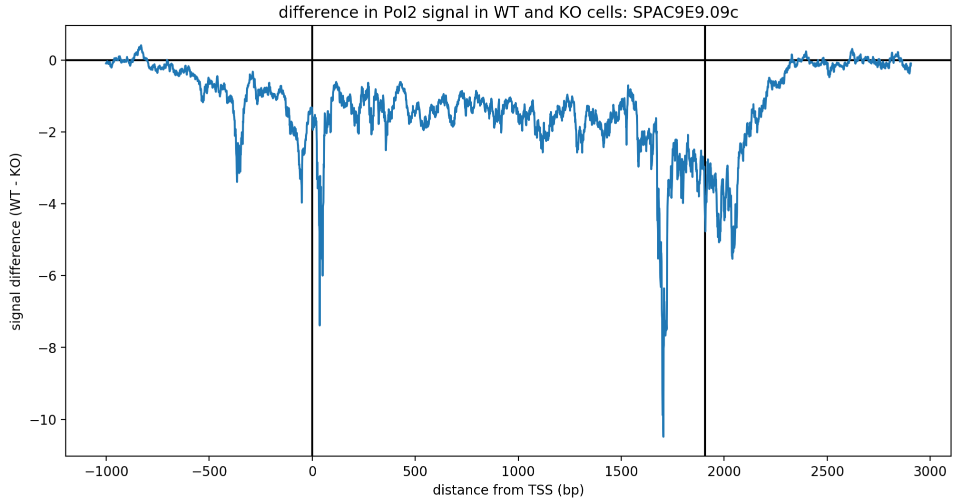

difference in Pol2 signal in WT and KO cells: SPAC9E9.09c
signal difference (WT - KO)
distance from TSS (bp)


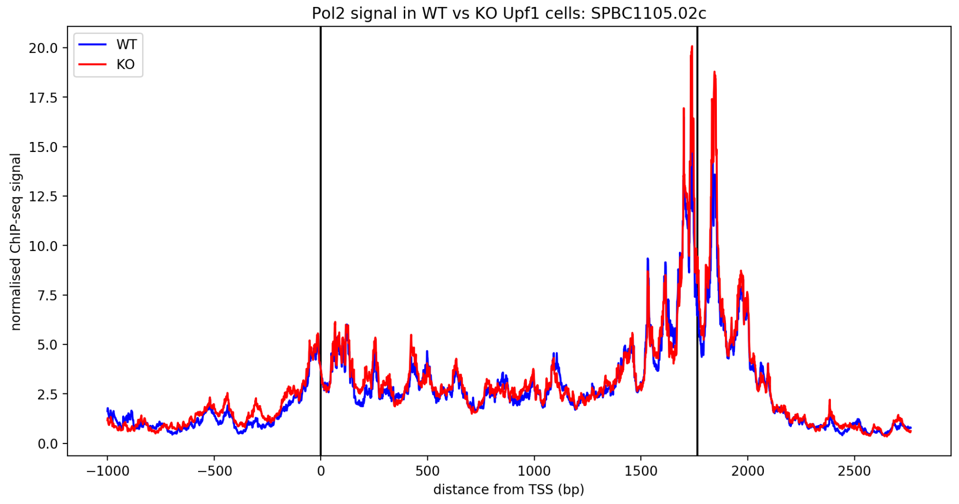

Pol2 signal in WT vs KO Upf1 cells: SPBC1105.02c
WT
KO
normalised ChIP-seq signal
distance from TSS (bp)

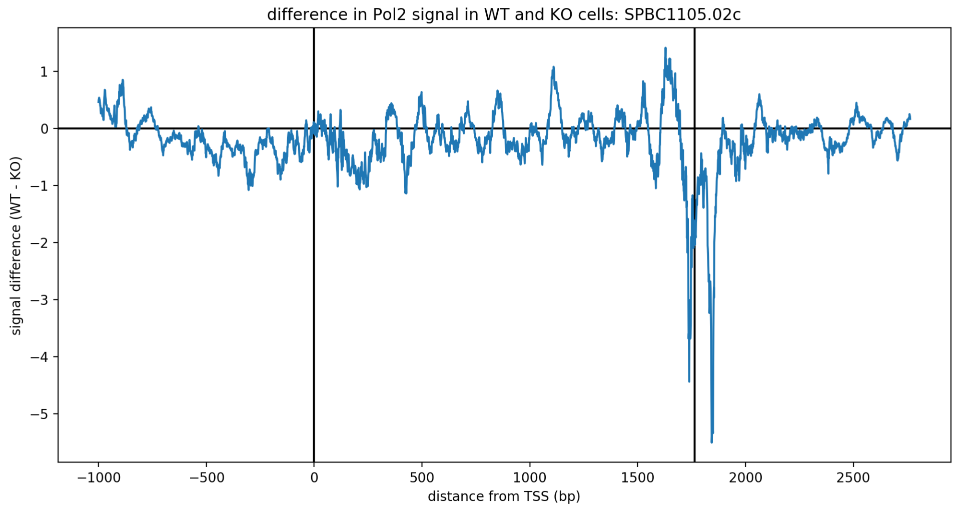
difference in Pol2 signal in WT and KO cells: SPBC1105.02c
signal difference (WT - KO)
distance from TSS (bp)
1
0
−1
−2
−3
−4
−5
−1000
−500
0
500
1000
1500
2000
2500

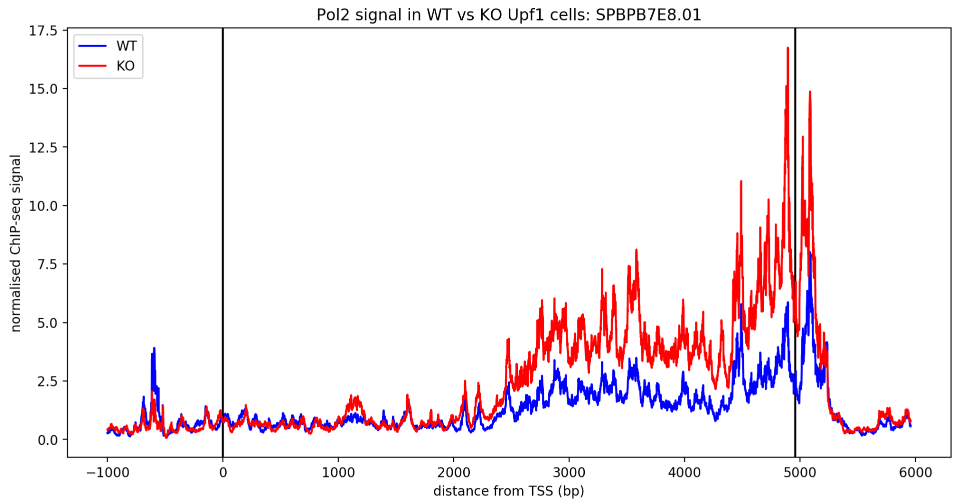
Pol2 signal in WT vs KO Upf1 cells: SPBPB7E8.01
WT
KO
normalised ChIP-seq signal
distance from TSS (bp)
17.5
15.0
12.5
10.0
7.5
5.0
2.5
0.0
−1000
0
1000
2000
3000
4000
5000
6000

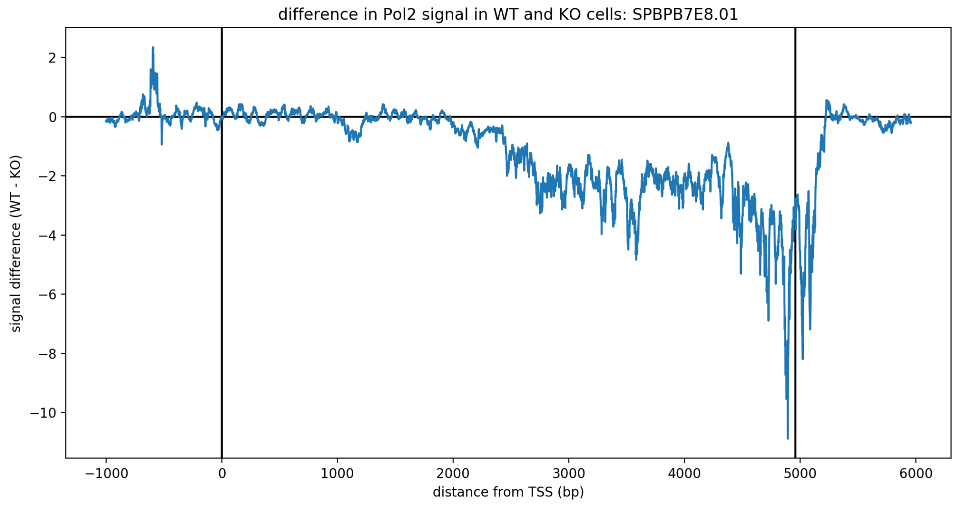
difference in Pol2 signal in WT and KO cells: SPBPB7E8.01
signal difference (WT - KO)
distance from TSS (bp)
2
0
−2
−4
−6
−8
−10
−1000
0
1000
2000
3000
4000
5000
6000

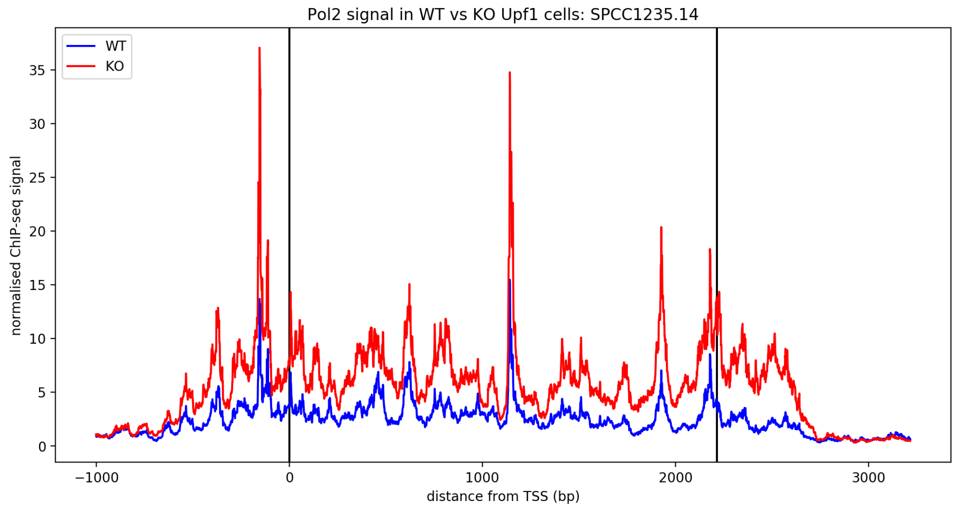
Pol2 signal in WT vs KO Upf1 cells: SPCC1235.14
WT
KO
normalised ChIP-seq signal
distance from TSS (bp)
35
30
25
20
15
10
5
0
−1000
0
1000
2000
3000

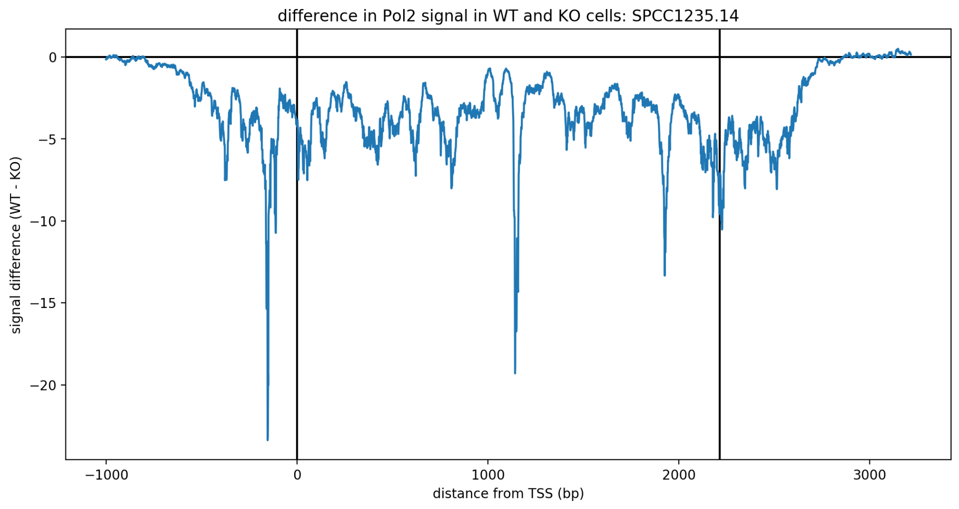


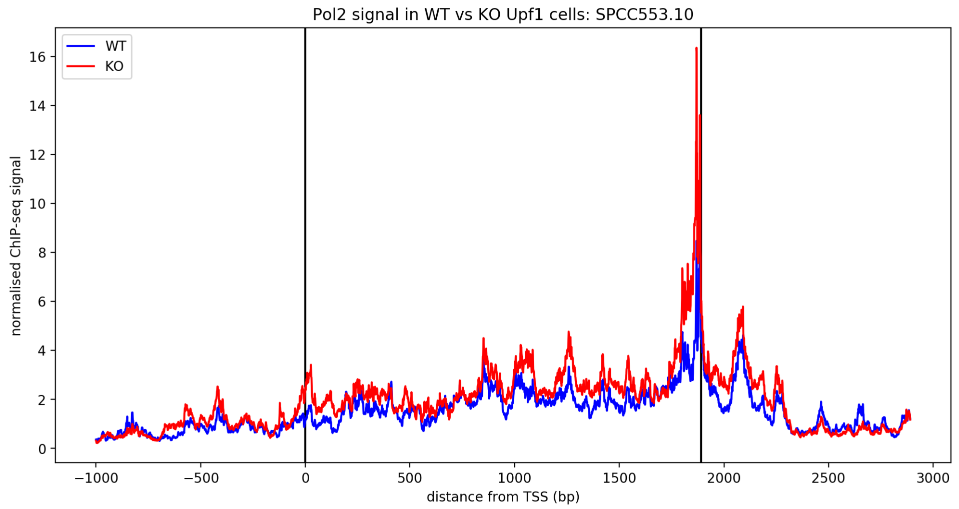


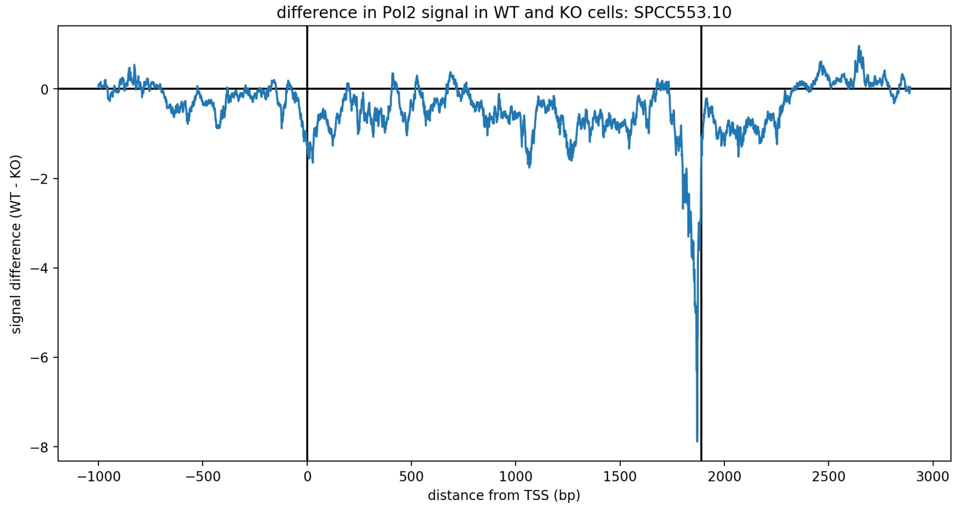


Published from selectedMisregGenesPlots.pmd using Pweave 0.30.3 on 24-06-2020.

File failed to load: https://cdnjs.cloudflare.com/ajax/libs/mathjax/2.7.1/jax/input/LaTeX/config.js
