## Supplementat Table 1 for "Genome-wide chromosomal association of Upf1 is linked to Pol II transcription in *Schizosaccharomyces pombe*"

Table S1 - Upf1 associated genes

| upf1_asyn-pe03systematic_ids_50 | upf1_Spha-pe03systematic_ids_50 |
|---|---|
| SPAC212.11 | SPAC212.11 |
| SPAC212.09c | SPAC212.10 |
| SPAC212.12 | SPAC212.09c |
| SPAC212.06c | SPAC212.08c |
| SPAC212.01c | SPAC212.12 |
| SPAC977.18 | SPAC212.06c |
| SPAC750 | SPAC212.04c |
| SPAC977.14c | SPAC212.01c |
| SPAC1F8.07c | SPAC977.01 |
| SPAC1F8.08 | SPAC977.18 |
| SPNCRNA.624 | SPAC977.02 |
| SPAC24B11.14 | SPAC977.03 |
| SPAC630.08c | SPAC977.04 |
| SPNCRNA.642 | SPAC1F8.07c |
| SPAC630.15 | SPAC11D3.05 |
| SPAC2F7.04 | SPAC5H10.02c |
| SPAC2F7.05c | SPRRNA.10 |
| SPAC13A11.02c | SPAC5H10.06c |
| SPAC1F3.03 | SPAC13G6.09 |
| SPAC4G8.13c | SPAC13G6.10c |
| SPNCRNA.684 | SPAC13G6.12c |
| SPAC16C9.01c | SPAC13G6.13 |
| SPAC16C9.06c | SPAC24B11.09 |
| SPAC22H12.04c | SPAC24B11.13 |
| SPAC222.08c | SPAC806.02c |
| SPSNRNA.02 | SPAC12G12.05c |
| SPAC222.09 | SPSNORNA.29 |
| SPAC222.11 | SPAC12G12.04 |
| SPAC222.17 | SPNCRNA.639 |
| SPAC821.10c | SPAC630.08c |
| SPAC1A6.04c | SPATRNAPRO.01 |
| SPAC30D11.13 | SPAC630.10 |
| SPATRNASER.01 | SPSNORNA.01 |
| SPATRNAMET.01 | SPAC13C5.05c |
| SPAC30D11.12 | SPAC23E2.01 |
| SPAC56F8.14c | SPATRNAVAL.01 |
| SPAC56F8.13 | SPRRNA.11 |
| SPAC10F6.06 | SPAC24H6.13 |
| SPAC1565.01 | SPAC227.12 |
| SPAC19E9.03 | SPAC227.13c |
| SPAC9.04 | SPAC227.18 |
| SPAC9.08c | SPAC13A11.02c |
| SPAC9.09 | SPAC22G7.05 |
| SPAC57A7.05 | SPAC22G7.06c |

SPAC57A7.04c
SPRRNA.53
SPAC1705.02
SPAC1705.03c
SPNCRNA.80
SPAC23H4.06
SPAC23H4.04
SPATRNATHR.01
SPAC343.21
SPAC343.12
SPAC343.20
SPAC664.04c
SPAC664.05
SPATRNALYS.02
SPAC1002.13c
SPAC1002.20
SPAC1002.19
SPNCRNA.770
SPAC140.02
SPNCRNA.778
SPNCRNA.781
SPNCRNA.782
SPAC631.02
SPAC23C11.06c
SPNCRNA.798
SPAC13F5.03c
SPAC18G6.09c
SPAC22H10.06c
SPAC22H10.13
SPAC6B12.07c
SPNCRNA.828
SPAC32A11.02c
SPAC19A8.05c
SPAC19A8.04
SPAC4A8.11c
SPATRNAILE.02
SPAC4F8.08
SPNCRNA.200
SPAC4F8.07c
SPAC644.06c
SPAC6F6.07c
SPAC6F6.08c
SPAC1805.10
SPAC1805.11c
SPAC3F10.17
SPAC3F10.18c
SPRRNA.12
SPATRNAALA.01
SPAC1687.17c
SPAC222.08c
SPSNRNA.02
SPAC222.09
SPSNRNA.03
SPAC821.08c
SPAC821.09
SPAC821.10c
SPAC1A6.04c
SPAC30D11.13
SPATRNASER.01
SPATRNAMET.01
SPAC30D11.12
SPAC56F8.16
SPNCRNA.82
SPNCRNA.727
SPSNORNA.10
SPAC10F6.15
SPAC56E4.03
SPAC56E4.04c
SPAC1565.08
SPAC6F12.04
SPSNORNA.53
SPAC6F12.05c
SPAC6F12.10c
SPAC19E9.03
SPAC9.04
SPAC9.07c
SPAC9.08c
SPAC9.09
SPAC5D6.02c
SPAC5D6.01
SPAC57A7.04c
SPAC167.08
SPAC1705.02
SPAC1705.03c
SPAC23H4.07c
SPAC23H4.06
SPATRNATHR.01
SPAC343.21
SPAC343.12
SPAC343.20
SPAC343.17c
SPAC343.16

SPAC513.01c
SPAC2E1P3.01
SPAC2E1P3.02c
SPNCRNA.862
SPAPB2C8.01
SPNCRNA.865
SPAPB24D3.06c
SPAPB24D3.07c
SPNCRNA.210
SPNCRNA.866
SPNCRNA.31
SPNCRNA.873
SPNCRNA.213
SPAC3G9.11c
SPATRNATHR.04
SPAC3G9.03
SPAC6G10.11c
SPAC26A3.01
SPAC8E11.02c
SPAC3H5.08c
SPAC3A11.10c
SPAC3A11.09
SPNCRNA.913
SPAC17A2.09c
SPNCRNA.917
SPAC17A2.11
SPNCRNA.925
SPAC8C9.04
SPAC8C9.14
SPAC15A10.09c
SPNCRNA.84
SPNCRNA.230
SPNCRNA.231
SPNCRNA.232
SPATRNAALA.04
SPATRNAGLU.03
SPATRNAILE.03
SPNCRNA.233
SPATRNAILE.04
SPNCRNA.333
SPNCRNA.396
SPATRNAGLU.04
SPATRNAALA.05
SPDHI
SPNCRNA.234
SPNCRNA.95
SPAC664.04c
SPAC664.05
SPAC664.06
SPSNORNA.31
SPAC664.07c
SPAC664.11
SPAC1002.13c
SPAC140.01
SPNCRNA.770
SPAC140.02
SPAP14E8.02
SPAP14E8.03
SPAC3H1.07
SPAC3H1.08c
SPAC631.02
SPAC18G6.09c
SPAC13D6.02c
SPNCRNA.812
SPAC6C3.02c
SPAC6C3.04
SPNCRNA.814
SPSNORNA.50
SPAC6B12.15
SPSNORNA.11
SPNCRNA.826
SPAC23H3.09c
SPAC4A8.04
SPSNRNA.06
SPAC4A8.09c
SPAC644.06c
SPAPB2B4.01c
SPSNORNA.54
SPAC6F6.07c
SPAC3F10.18c
SPAC8F11.10c
SPAC513.01c
SPAC2E1P3.02c
SPAC2E1P3.03c
SPAC2E1P3.04
SPAC2E1P3.05c
SPNCRNA.862
SPNCRNA.863
SPNCRNA.865
SPAPB24D3.07c
SPAPB1A11.01
SPRRNA.13

SPAC4H3.10c
SPAC1071.10c
SPNCRNA.92
SPNCRNA.944
SPNCRNA.93
SPAC2F3.09
SPAC11G7.01
SPAPB18E9.04c
SPAPB15E9.01c
SPAPB15E9.02c
SPAPB15E9.06
SPAPB15E9.03c
SPAC27E2.03c
SPNCRNA.53
SPAC27E2.11c
SPAC27E2.13
SPAC27E2.04c
SPAC19G12.08
SPATRNALEU.04
SPAC19G12.09
SPAC19G12.10c
SPAC19G12.13c
SPAC19G12.12
SPAC19G12.14
SPAC23A1.09
SPNCRNA.957
SPAC23A1.10
SPAC26H5.08c
SPAC26H5.09c
SPAC26H5.10c
SPAC25B8.12c
SPNCRNA.968
SPAC1F7.13c
SPATRNASER.03
SPATRNAMET.03
SPAC2C4.04c
SPAC29E6.07
SPAC29E6.08
SPNCRNA.986
SPAC16.02c
SPAC16.05c
SPNCRNA.99
SPAC9E9.02
SPAC9E9.01
SPAC9E9.03
SPAC17C9.16c
SPNCRNA.873
SPAC31G5.11
SPATRNASER.02
SPAC24C9.06c
SPAC24C9.12c
SPAC589.09
SPAC589.10c
SPAC3G9.10c
SPATRNATHR.04
SPAPB1E7.03
SPAPB1E7.04c
SPAPB1E7.07
SPAPB1E7.08c
SPAC26A3.13c
SPAC8E11.10
SPSNORNA.20
SPNCRNA.906
SPAC959.05c
SPAC3H5.10
SPAC3H5.04
SPAC3H5.05c
SPAC328.09
SPAC328.10c
SPAC17A2.08c
SPAC17A2.09c
SPRRNA.16
SPAC8C9.08
SPATRNACYS.03
SPNCRNA.524
SPATRNAARG.02
SPAP7G5.03
SPAP7G5.06
SPNCRNA.84
SPNCRNA.230
SPNCRNA.231
SPNCRNA.232
SPATRNAALA.04
SPATRNAGLU.03
SPATRNAILE.03
SPNCRNA.233
SPATRNAILE.04
SPNCRNA.333
SPNCRNA.396
SPATRNAGLU.04
SPATRNAALA.05
SPDHI

SPAC17C9.13c
SPAC17C9.12
SPAC17C9.03
SPNCRNA.995
SPNCRNA.996
SPATRNAA
SPAC27D7.12c
SPAC637.03
SPAC1093.01
SPAC1834.08
SPATRNATHR.05
SPNCRNA.252
SPAC1834.11c
SPAPYUG7.03c
SPAPYUG7.04c
SPAC1782.07
SPATRNAALA.06
SPAC22F8.07c
SPAC4F10.13c
SPNCRNA.1030
SPAC4F10.14c
SPAC4F10.15c
SPNCRNA.1031
SPAC19B12.02c
SPAC29A4.17c
SPAC29A4.16
SPAC29A4.11
SPAC29A4.22
SPAC29A4.12c
SPNCRNA.1054
SPAC26F1.06
SPAC26F1.05
SPAP8A3.04c
SPNCRNA.1074
SPAP8A3.13c
SPAC29B12.04
SPNCRNA.1082
SPAC29B12.07
SPAC29B12.08
SPNCRNA.1084
SPAC1039.05c
SPNCRNA.274
SPAC1039.06
SPAC922.04
SPNCRNA.282
SPAC869.02c
SPNCRNA.234
SPNCRNA.95
SPAC1556.02c
SPAC1F12.02c
SPAC4H3.10c
SPAC1071.09c
SPAC1071.10c
SPNCRNA.92
SPRRNA.18
SPAC926.05c
SPAC926.09c
SPAC2F3.09
SPAC2F3.18c
SPAPB18E9.02c
SPAPB18E9.05c
SPAPB15E9.01c
SPAPB15E9.02c
SPAPB15E9.03c
SPAC27E2.03c
SPNCRNA.53
SPAC27E2.11c
SPAC27E2.13
SPAC27E2.08
SPAC19G12.06c
SPATRNALEU.04
SPAC19G12.10c
SPAC19G12.15c
SPAC19G12.16c
SPAC23A1.09
SPNCRNA.957
SPAC23A1.10
SPAC26H5.06
SPAC26H5.07c
SPAC26H5.10c
SPAC23D3.12
SPAC29E6.08
SPAC16.04
SPAC16.05c
SPNCRNA.99
SPAC9E9.03
SPAC9E9.08
SPAC9E9.09c
SPATRNAA
SPAC27D7.12c
SPAC1834.08
SPATRNATHR.05

SPAC186.04c
SPNCRNA.61
SPAC186.08c
SPAC750.01
SPAC750.05c
SPAC750.06c
SPAC750.07c
SPAC750.08c
SPBPB21E7.04c
SPNCRNA.1305
SPBPB21E7.08
SPNCRNA.1323
SPBC1198.07c
SPNCRNA.1324
SPBC1198.14c
SPBC660.06
SPBC31E1.04
SPBTRNAGLN.03
SPBC800.04c
SPBTRNAGLY.04
SPBTRNAALA.07
SPBC1773.03c
SPBC1773.04
SPBTRNAGLY.05
SPBTRNAARG.04
SPBTRNAGLN.04
SPBC1271.03c
SPBC428.02c
SPBTRNAHIS.01
SPRRNA.28
SPBC1685.09
SPBC1685.10
SPNCRNA.1366
SPBC1685.12c
SPNCRNA.1367
SPBC1685.13
SPBC649.04
SPBC649.05
SPBC354.11c
SPNCRNA.313
SPBC354.12
SPBC354.13
SPNCRNA.1375
SPBC839.15c
SPNCRNA.1376
SPBC839.16
SPAC19B12.01
SPAC4F10.22
SPAC19B12.02c
SPAPB8E5.03
SPAPB8E5.05
SPAPB8E5.06c
SPAC1B3.05
SPNCRNA.1038
SPSNORNA.42
SPAC1B3.15c
SPAC1B3.16c
SPAC1006.07
SPAC26F1.14c
SPSNORNA.16
SPAC26F1.06
SPAC26F1.05
SPAC13D1.01c
SPAC19D5.09c
SPAC11E3.12
SPAC11E3.13c
SPAC11E3.15
SPAP8A3.02c
SPAP8A3.06
SPAP8A3.07c
SPAC4D7.08c
SPSNORNA.17
SPSNORNA.19
SPAC3G6.05
SPNCRNA.265
SPAC29B12.04
SPAC29B12.08
SPAC29B12.11c
SPATRNASER.04
SPAC922.04
SPAC869.11
SPAC186.05c
SPAC186.06
SPNCRNA.310
SPAC750.04c
SPAC1952
SPAC750.05c
SPAC750.06c
SPAC750.07c
SPAC750.08c
SPBC1348.01
SPBC1348.02

SPBC947.15c
SPNCRNA.1378
SPBC947.14c
SPBC947.04
SPBPJ4664.02
SPNCRNA.1382
SPBC119.02
SPBC119.03
SPNCRNA.324
SPBC119.05c
SPBC577.14c
SPNCRNA.1389
SPNCRNA.1390
SPBC36.03c
SPNCRNA.1391
SPBC36.13
SPBC713.11c
SPBC713.12
SPBC646.17c
SPBTRNAHIS.02
SPBP35G2.16c
SPBP35G2.02
SPBC337.07c
SPBC337.08c
SPBC1709.05
SPBTRNAARG.05
SPBC651.03c
SPBC31A8.02
SPBC3D6.02
SPBC30B4.09
SPNCRNA.570
SPBC3D6.16
SPSNORNA.21
SPBC30B4.04c
SPBC8D2.03c
SPBC8D2.04
SPBC8D2.18c
SPBP22H7.08
SPBC32H8.15
SPBC32H8.12c
SPBC11B10.07c
SPNCRNA.24
SPBC11B10.08
SPBC83.13
SPBC29B5.03c
SPBC28F2.03
SPBC1348.03
SPBC1348.05
SPBPB10D8.04c
SPBPB10D8.05c
SPBPB10D8.06c
SPBPB10D8.07c
SPBC1683.01
SPBC1683.10c
SPNCRNA.100
SPBC1198.06c
SPBTRNAGLN.02
SPBTRNAGLN.03
SPBC800.04c
SPBTRNAGLY.04
SPBTRNAALA.07
SPBC1773.03c
SPBTRNAGLY.05
SPBC582.03
SPBTRNAHIS.01
SPRRNA.28
SPBC428.11
SPSNRNA.04
SPBC1685.10
SPBC1685.11
SPRRNA.29
SPNCRNA.134
SPBC354.12
SPBTRNALYS.06
SPNCRNA.317
SPNCRNA.1375
SPBC839.15c
SPNCRNA.1376
SPBC947.04
SPBC947.03c
SPBPJ4664.02
SPBC119.02
SPBC119.10
SPBC119.11c
SPBC530.02
SPBC530.05
SPBC530.06c
SPBC530.10c
SPBC36.03c
SPBC646.17c
SPBTRNAHIS.02
SPBP35G2.16c

SPBC28F2.12
SPBTRNAASN.01
SPBTRNAMET.05
SPBTRNATYR.02
SPBTRNALEU.06
SPBTRNAGLY.07
SPBTRNALYS.07
SPBTRNAILE.05
SPBTRNAALA.08
SPBTRNAVAL.05
SPBTRNAGLU.06
SPBTRNAA
SPNCRNA.359
SPNCRNA.361
SPNCRNA.362
SPBTRNAILE.06
SPBTRNAALA.09
SPBTRNAVAL.06
SPNCRNA.367
SPNCRNA.368
SPNCRNA.369
SPNCRNA.370
SPBTRNAVAL.07
SPBTRNAALA.10
SPBTRNAILE.07
SPNCRNA.371
SPNCRNA.372
SPNCRNA.373
SPBTRNATYR.03
SPBTRNALEU.07
SPBTRNAGLY.08
SPBTRNALYS.08
SPBTRNAILE.08
SPBTRNAALA.11
SPBTRNAVAL.08
SPBTRNAGLU.07
SPBTRNAARG.07
SPNCRNA.374
SPBC21B10.15
SPBC21B10.03c
SPBC19C2.07
SPBC2F12.05c
SPBC2F12.04
SPBC1D7.04
SPBC1D7.03
SPBC3H7.13
SPBC337.02c
SPBC337.08c
SPBC1709.05
SPBC409.08
SPBC409.09c
SPBC3D6.02
SPSNORNA.21
SPBC30B4.04c
SPBC8D2.03c
SPBC8D2.04
SPBC8D2.05c
SPBC8D2.17
SPBC8D2.18c
SPBC32H8.12c
SPBTRNATHR.06
SPRRNA.33
SPBC83.18c
SPNCRNA.1444
SPBC27.06c
SPBC27.08c
SPBC28F2.12
SPBTRNAASN.01
SPBTRNAMET.05
SPBTRNATYR.02
SPBTRNALEU.06
SPBTRNAGLY.07
SPBTRNALYS.07
SPBTRNAILE.05
SPBTRNAALA.08
SPBTRNAVAL.05
SPBTRNAGLU.06
SPBTRNAA
SPNCRNA.359
SPNCRNA.360
SPNCRNA.361
SPNCRNA.362
SPNCRNA.365
SPBTRNAILE.06
SPBTRNAALA.09
SPBTRNAVAL.06
SPNCRNA.367
SPNCRNA.368
SPNCRNA.369
SPNCRNA.370
SPBTRNAVAL.07
SPBTRNAALA.10

| | |
|---|---|
| SPBTRNAPRO.05 | SPBTRNAILE.07 |
| SPBC16E9.16c | SPNCRNA.371 |
| SPBC1E8.05 | SPNCRNA.372 |
| SPBC1A4.01 | SPNCRNA.373 |
| SPBP23A10.11c | SPBTRNATYR.03 |
| SPNCRNA.385 | SPBTRNALEU.07 |
| SPBC29A3.04 | SPBTRNAGLY.08 |
| SPBC18E5.04 | SPBTRNALYS.08 |
| SPBC18E5.05c | SPBTRNAILE.08 |
| SPBC1815.01 | SPBTRNAALA.11 |
| SPBTRNAPRO.06 | SPBTRNAVAL.08 |
| SPNCRNA.1502 | SPBTRNAGLU.07 |
| SPBC24C6.04 | SPBTRNAARG.07 |
| SPBC19G7.06 | SPNCRNA.374 |
| SPBC19G7.07c | SPBC21B10.04c |
| SPBC1921.05 | SPBC21B10.15 |
| SPNCRNA.1511 | SPBC19C2.07 |
| SPBC1921.07c | SPBC2F12.14c |
| SPBC21D10.12 | SPSNORNA.22 |
| SPBC17F3.01c | SPNCRNA.1459 |
| SPBC557.02c | SPBC1D7.03 |
| SPNCRNA.1527 | SPBC9B6.02c |
| SPBC29A10.08 | SPBC1E8.04 |
| SPBC29A10.09c | SPBC1E8.05 |
| SPBC3E7.16c | SPBP23A10.10 |
| SPBC4F6.04 | SPBP23A10.11c |
| SPBC685.07c | SPNCRNA.1482 |
| SPBC32F12.01c | SPBTRNAASN.02 |
| SPNCRNA.1550 | SPRRNA.39 |
| SPBC32F12.11 | SPBTRNAGLY.09 |
| SPBC19C7.03 | SPBC1711.13 |
| SPBC19C7.04c | SPBC1711.15c |
| SPNCRNA.1554 | SPBC1711.14 |
| SPNCRNA.26 | SPBC17G9.10 |
| SPNCRNA.1561 | SPBC17G9.11c |
| SPNCRNA.411 | SPBC1815.01 |
| SPNCRNA.1562 | SPBTRNAPRO.06 |
| SPSNRNA.01 | SPBC24C6.04 |
| SPBC15D4.05 | SPBC24C6.05 |
| SPBC13E7.08c | SPBC19G7.16 |
| SPBC13E7.09 | SPNCRNA.1508 |
| SPSNRNA.05 | SPBC1921.04c |
| SPBC19F8.08 | SPSNORNA.34 |
| SPBC25H2.07 | SPBC1921.05 |
| SPBC25H2.06c | SPNCRNA.1511 |
| SPBC3B8.09 | SPBC12C2.11 |

SPBC3B8.07c
SPBC3B8.06
SPNCRNA.1596
SPSNORNA.33
SPBC3B8.02
SPNCRNA.1607
SPBC1105.14
SPBC1105.13c
SPBC887.15c
SPNCRNA.1616
SPBC317.01
SPBP8B7.15c
SPBP8B7.16c
SPBTRNALYS.09
SPBTRNATYR.04
SPBC23E6.08
SPRRNA.36
SPBC26H8.06
SPSNRNA.07
SPBC26H8.10
SPBC26H8.11c
SPNCRNA.111
SPBC32C12.02
SPBC215.05
SPBC215.09c
SPBC215.10
SPBC56F2.12
SPBC56F2.02
SPBC14F5.04c
SPBC14F5.05c
SPBC16G5.14c
SPBC1652.01
SPBC8E4.02c
SPBP4G3.02
SPBP4G3.03
SPNCRNA.1695
SPBPB2B2.18
SPBCPT2R1.06c
SPBCPT2R1.08c
SPCC757.05c
SPNCRNA.452
SPCC757.07c
SPCC757.15
SPCC757.09c
SPCC757.11c
SPCTRNAHIS.03
SPBC365.16
SPBC2G5.05
SPBC2G5.06c
SPBC3E7.15c
SPBC3E7.16c
SPBC4F6.04
SPBC32F12.11
SPNCRNA.1551
SPBC19C7.03
SPBC19C7.04c
SPBP4H10.11c
SPBC15D4.04
SPSNRNA.01
SPBC15D4.05
SPSNRNA.05
SPBC17D1.01
SPBTRNAPRO.08
SPBC11C11.09c
SPBC13A2.03
SPBC13A2.04c
SPBPB7E8.01
SPBC1105.01
SPBC1105.02c
SPBC1105.03c
SPBC1105.05
SPBC1105.10
SPBC1105.11c
SPBC887.15c
SPBC16D10.06
SPBC317.01
SPBP8B7.32
SPBP8B7.05c
SPBP8B7.04
SPBP8B7.15c
SPBP8B7.16c
SPBTRNALYS.09
SPBTRNATYR.04
SPNCRNA.445
SPRRNA.36
SPBC26H8.06
SPSNRNA.07
SPBC26H8.10
SPBC26H8.11c
SPBC215.05
SPBC215.08c
SPNCRNA.1659

SPCC757.12
SPCC613.04c
SPCC613.05c
SPCC613.06
SPNCRNA.1109
SPCC330.04c
SPCTRNAGLY.10
SPCC330.05c
SPCC1235.01
SPCC1235.17
SPCC1235.14
SPCC548.05c
SPNCRNA.460
SPCC548.06c
SPCC794.02
SPCC794.01c
SPCC794.08
SPCC794.09c
SPCC553.11c
SPCC736.15
SPNCRNA.1133
SPCC594.01
SPCC594.02c
SPNCRNA.1134
SPCC594.03
SPCC962.05
SPCC962.06c
SPCC1672.01
SPCC1672.02c
SPCC1672.14
SPCC1183.08c
SPCC31H12.04c
SPNCRNA.472
SPNCRNA.1153
SPCC18B5.02c
SPNCRNA.1156
SPCC1020.05
SPCC1020.14
SPCC1393.08
SPCC63.13
SPCC63.14
SPCC24B10.21
SPCPB16A4.03c
SPCPB16A4.06c
SPCC1742.01
SPCC1795.11
SPBC1347.13c
SPBC56F2.12
SPNCRNA.1666
SPBC56F2.11
SPBTRNAGLU.08
SPBC14F5.04c
SPRRNA.37
SPBC14F5.05c
SPBTRNAASN.04
SPRRNA.38
SPBC16G5.14c
SPNCRNA.497
SPBC1652.01
SPBC1289.03c
SPBC1289.17
SPBC1289.15
SPBC8E4.02c
SPBC8E4.01c
SPBP4G3.02
SPBPB2B2.18
SPBPB2B2.19c
SPBCPT2R1.01c
SPBCPT2R1.04c
SPBCPT2R1.06c
SPBCPT2R1.07c
SPBCPT2R1.10
SPBCPT2R1.08c
SPRRNA.50
SPRRNA.51
SPNCRNA.1094
SPNCRNA.1095
SPCP20C8.01c
SPNCRNA.449
SPCP20C8.02c
SPCP20C8.03
SPCC1884.01
SPNCRNA.1096
SPNCRNA.1097
SPCC1884.02
SPCC757.02c
SPNCRNA.1099
SPCC757.03c
SPCC757.04
SPCC757.05c
SPCC757.07c
SPCC757.08

SPCC1259.01c
SPCTRNAALA.12
SPCTRNAVAL.12
SPNCRNA.480
SPCTRNASER.09
SPCTRNAARG.10
SPCTRNAA
SPCTRNAARG.11
SPNCRNA.1172
SPNCRNA.481
SPCTRNALEU.11
SPCC4B3
SPCTRNALYS.10
SPNCRNA.482
SPCTRNAARG.12
SPCTRNAVAL.09
SPCTRNATHR.08
SPNCRNA.483
SPCTRNALEU.12
SPCTRNAGLU.10
SPCTRNALEU.13
SPNCRNA.484
SPCTRNATHR.09
SPCTRNAVAL.10
SPCTRNAARG.13
SPCTRNALYS.11
SPCC550.10
SPCTRNAVAL.11
SPCC550.11
SPCC1322.04
SPNCRNA.1186
SPCC1322.10
SPCC338.12
SPCTRNATHR.10
SPCC1281.06c
SPCC622.08c
SPCC622.09
SPCC622.10c
SPCC622.12c
SPCC1753.02c
SPCC13B11.01
SPCC13B11.02c
SPCC663.03
SPCC663.04
SPCC417.05c
SPCC417.06c
SPCC757.09c
SPCC757.10
SPCC757.11c
SPCTRNAHIS.03
SPCC757.12
SPCC757.13
SPNCRNA.1102
SPCC613.04c
SPCC613.05c
SPNCRNA.1103
SPCC613.06
SPCC330.02
SPCC330.03c
SPCC330.21
SPCC330.06c
SPCC330.09
SPCC330.10
SPCC330.12c
SPCC320.08
SPCC1235.01
SPCC1235.02
SPCC1235.03
SPCC1235.14
SPCC548.03c
SPCC548.05c
SPNCRNA.460
SPCC548.06c
SPCC794.08
SPCC794.09c
SPCC794.12c
SPCC553.11c
SPCC553.10
SPCC736.15
SPCC594.02c
SPCC306.11
SPCC4G3.19
SPCC4G3.02
SPCC364.07
SPCTRNASER.07
SPCC364.04c
SPCC364.03
SPCP31B10.06
SPCP31B10.07
SPCP31B10.08c
SPSNORNA.32
SPCC1672.10

SPCC417.08
SPCTRNAASN.06
SPRRNA.06
SPCC1223.01
SPNCRNA.10
SPCC1223.02
SPNCRNA.1239
SPCC297.03
SPNCRNA.1240
SPCC737.04
SPNCRNA.1241
SPNCRNA.1242
SPCC74.03c
SPCC4F11.03c
SPCC4F11.04c
SPCC1906.01
SPCC1906.02c
SPCC1739.08c
SPCC1739.10
SPCC1739.13
SPCPB1C11.02
SPNCRNA.1258
SPCC576.03c
SPCC576.08c
SPCC576.11
SPNCRNA.1263
SPCC576.17c
SPCC126.12
SPCC830.07c
SPCC965.07c
SPNCRNA.550
SPCC70.03c
SPCC70.10
SPCP1E11.04c
SPCP1E11.05c
SPCP1E11.06
SPCP1E11.07c
SPCP1E11.08
SPCP1E11.09c
SPCC569.08c
SPCC569.09
SPNCRNA.1296
SPNCRNA.1297
SPCC569.05c
SPCC1020.06c
SPCC1020.05
SPCC1020.14
SPCC1393.08
SPRRNA.25
SPCC1393.10
SPRRNA.55
SPCC24B10.14c
SPCTRNAARG.09
SPRRNA.26
SPNCRNA.475
SPNCRNA.526
SPCC24B10.21
SPCC24B10.22
SPCTRNAASN.05
SPCPB16A4.03c
SPNCRNA.477
SPCPB16A4.04c
SPCC1742.01
SPCC1795.12c
SPCC1795.11
SPCC1795.05c
SPNCRNA.128
SPCC1795.04c
SPCC1259.08
SPCC1259.09c
SPCTRNAALA.12
SPCTRNAVAL.12
SPNCRNA.480
SPCTRNASER.09
SPCTRNAARG.10
SPCTRNAA
SPCTRNAARG.11
SPNCRNA.1172
SPNCRNA.481
SPCC4B3
SPCTRNALYS.10
SPNCRNA.482
SPCTRNAARG.12
SPCTRNAVAL.09
SPCTRNATHR.08
SPNCRNA.483
SPCTRNALEU.12
SPCTRNAGLU.10
SPCTRNALEU.13
SPNCRNA.484

SPCTRNATHR.09
SPCTRNAVAL.10
SPCTRNAARG.13
SPNCRNA.485
SPCTRNALYS.11
SPCTRNAPHE.05
SPCC4B3.18
SPCC4B3.03c
SPNCRNA.1186
SPCC1322.10
SPCC1281.06c
SPNCRNA.1194
SPCC622.08c
SPCC622.09
SPCC622.10c
SPCC622.12c
SPCC11E10.01
SPCC162.04c
SPCC13B11.01
SPCC13B11.02c
SPCC417.08
SPCTRNASER.11
SPCTRNAMET.07
SPCTRNAASN.06
SPRRNA.06
SPCC417.15
SPNCRNA.32
SPCC417.16
SPCC285.07c
SPCC1223.01
SPNCRNA.10
SPCC1223.02
SPNCRNA.1236
SPCC1223.10c
SPCC1223.13
SPCC74.06
SPCC18.01c
SPCC1906.01
SPCC1739.01
SPCTRNAGLY.12
SPCPB1C11.01
SPCTRNASER.13
SPCC576.03c
SPCC576.08c
SPCC576.09
SPNCRNA.119

SPCC576.10c
SPCC576.11
SPCC576.12c
SPCC1620.14c
SPCC1919.15
SPNCRNA.1276
SPCC1840.01c
SPCC1840.02c
SPCC1840.03
SPCC965.04c
SPNCRNA.1280
SPCC965.07c
SPRRNA.07
SPNCRNA.121
SPCC1494.11c
SPCC70.03c
SPNCRNA.1290
SPCC70.04c
SPCC70.12c
SPCC70.05c
SPCC70.08c
SPCC70.09c
SPCC70.10
SPCC1827.01c
SPCC1827.02c
SPCC1827.04
SPCC1827.05c
SPCC1827.06c
SPNCRNA.1294
SPCC1827.07c
SPCP1E11.03
SPCP1E11.04c
SPCP1E11.05c
SPCP1E11.06
SPCP1E11.08
SPCP1E11.09c
SPCP1E11.10
SPCP1E11.11
SPCC569.08c
SPCC569.09
SPCC569.07
SPNCRNA.1295
SPNCRNA.29
SPCC569.06
SPNCRNA.1296
SPNCRNA.1297

SPCC569.05c
SPCC569.04
SPCC569.03
SPCC569.02c
SPNCRNA.1298
SPCC569.01c
SPRRNA.52
SPNCRNA.1299
