## Supplementat Table 2 for "Genome-wide chromosomal association of Upf1 is linked to Pol II transcription in *Schizosaccharomyces pombe*"

Table S2 up_and_downregulated_genes_upf1_rodriguez_dset

| UP REGULATED GENES IN UPF1 KO | DOWN REGULATED GENES IN UPF1 KO |
|---|---|
| SPBC1773.13 | SPBC26H8.11c |
| SPNCRNA.131 | SPAC16A10.05c;dad1 |
| SPAC1039.02 | SPAPJ760.02c;app1 |
| SPBC1773.03c | SPAC10F6.06;vip1 |
| SPBC21C3.16c;spt4 | SPAC26F1.04c;mrf1;etr1 |
| SPBP8B7.12c;sma3;fta3 | SPBP8B7.26 |
| SPBC660.15 | SPBC428.10 |
| SPBC16G5.03 | SPBC27B12.02;SPBC30B4.09;SPBC30B4.10 |
| SPCC550.08 | SPBP8B7.17c |
| SPBC1773.14;arg7 | SPAC24C9.10c;mrp4 |
| SPBP7E8.01;SPBPB7E8.01 | SPBC365.02c;cox10 |
| SPBP23A10.03c | SPNCRNA.65 |
| SPBC1778.05c | SPAC869.09 |
| SPAC22F3.05c;alp41 | SPAC227.15 |
| SPAC27D7.11c | SPBC119.02;ubc4 |
| SPBPB2B2.06c | SPCPB16A4.06c |
| SPBC18E5.09c | SPBC1105.11c;clo5;hht3;H3.3;h3.3 |
| SPAC688.06c;slx4 | SPCC1322.10 |
| SPCC965.05c;thp1;tng | SPAC18B11.07c;sng1;rhp6;ubc2 |
| SPAC56F8.14c | SPBC32F12.06;pch1 |
| SPCC622.17;apn1 | SPAC24B11.13;SPAC806.01;hem3 |
| SPAC139.04c;fap2 | SPBC14C8.03 |
| SPBC1347.07;rex2 | SPCC5E4.05c |
| SPAPB24D3.03 | SPBC2F12.05c |
| SPAC57A10.03;cyp2;cyp1 | SPBC16G5.16 |
| SPCC132.05c;SPCC338.19 | SPAC24B11.05 |
| SPAC20G4.05c | SPBC947.10 |
| SPAC11D3.18c | SPBC11C11.06c |
| SPAC4F10.12;sma1;fta1 | SPAC26F1.07 |
| SPCC1494.01;SPCC965.15 | NA |
| SPAC11G7.05c | SPAC1B3.03c;cyp5;wis2 |
| SPCC132.04c | SPNCRNA.92 |
| SPBC24C6.11;cwf14 | SPAC26H5.05 |
| SPCC794.05c | SPAC13G7.10 |
| NA | SPCC1919.04 |
| SPCC970.02 | SPCC1322.05c |
| SPAC23D3.12 | SPBC215.13 |
| SPCC1450.01c;SPCC191.12c | SPBC30D10.14 |
| SPAC2E1P3.03c;Tf2-3 | SPBC21C3.19 |
| SPAC27E2.08;Tf2-6 | SPNCRNA.41 |
| SPBC25B2.04c | SPAC821.10c;sod1 |
| SPBC1289.02c;uap2 | SPAC2C4.17c |
| SPAC5H10.05c | SPAC32A11.03c |
| SPAC13D1.01c;Tf2-7 | SPAC9G1.11c;spn4 |
| SPBC8E4.02c | SPCC553.04;cyp9 |
| SPAC25H1.04 | SPAC1639.01c;SPAC806.09c |
| SPBC146.10;mug57 | SPAC1565.01 |

| | |
|---|---|
| SPAC922.07c | SPNCRNA.58 |
| SPAC11H11.05c;sma6;fta6 | SPBC20F10.04c;rad62;nse4 |
| SPAC19D5.09c;SPAC13D1.02c;Tf2-8 | SPAC6F12.03c;fsv1 |
| SPBC18E5.14c | SPBC215.11c |
| SPAC6F6.04c | SPAC1B9.03;SPCC188.02;par1 |
| SPAC922.03 | SPCC757.07c;cta1;ctt1 |
| SPCC18B5.02c | SPAC3C7.12;noc1;tip1 |
| SPAC167.08;SPAC1705.01c;Tf2-2 | SPAC637.13c |
| SPCC663.07c | SPBC887.04c;lub1 |
| SPNCRNA.132 | SPBC36.04;cys1a;cys11 |
| SPCC1884.02;SPCC757.01;nic1 | SPBC16C6.02c;vps13B;vps13b;vps1302 |
| SPAC9.04;Tf2-1 | SPAC21B10.08c;SPBC21B10.08c |
| SPCC1281.07c | SPAC1565.06c;spg1;sid3;pld4 |
| SPCC285.05 | SPBC21C3.20c;git1 |
| SPCC576.17c | SPBP8B7.21;ubp3;ubp29 |
| SPAC1039.04 | SPAC25B8.04c |
| SPCC965.14c | SPBC21.06c;its10;pld1;cdc7 |
| SPCC1235.01;SPCC320.02c | SPBC83.17 |
| SPBC21C3.05;sap62 | SPBC24E9.03c;SPBC839.03c |
| SPBC1683.12 | SPBC1604.03c |
| SPAC1039.03 | SPBC725.02;mpr1;spy1;sph1 |
| SPAC22F8.11;plc1 | SPAC8C9.05 |
| SPAC11D3.01c | SPCC1259.08 |
| SPAC3F10.09;his6 | SPAC23G3.07c |
| SPBC1E8.04;SPBC1E8.04c;Tf2-10-pseudo | SPAC1952.04c |
| SPCC1840.10;lsm8 | SPBC3B8.07c;dsd1 |
| SPBC9B6.02c;SPBC9B6.02;Tf2-9 | SPCC1827.03c |
| SPBC1773.15 | SPBC3E7.10 |
| SPAC3H5.04;aar2 | SPAC5H10.13c;gmh2 |
| SPCPB1C11.03 | SPAC821.03c |
| SPBC1773.17c;SPBP26C9.01c | SPBC1711.15c |
| SPAC8A4.02c;SPCC188.09c | SPBC32F12.09;rum1 |
| SPCC970.12;mis18 | SPBC25H2.07;tif11 |
| SPCC663.12;cid12 | SPAC2H10.01 |
| SPBC409.12c | SPAC222.09;seb1 |
| SPBC17D1.01;SPBC17D11.09 | SPAC8E11.02c;rad24 |
| SPAC25H1.03;mug66 | SPAC23H3.09c;gly1 |
| SPAC323.05c | SPBC83.13 |
| SPBC26H8.01;nmt2;thi2 | SPAC13G7.11 |
| SPBC8D2.07c;pi057;sfc9 | SPAC2F3.11 |
| SPBC1861.05 | SPAC328.03;tps1;ggs1 |
| SPCC965.10 | SPAC16E8.11c;tfb1 |
| SPCC162.02c | SPCC622.08c;hta1 |
| SPBC18E5.08 | SPAC1786.02 |
| SPCC1235.14;ght5 | SPAPB8E5.02c;rpn502;rpn5-b;rpn5 |
| SPAC1039.07c | SPBP35G2.02 |
| SPBC3B8.08 | SPNCRNA.48 |

| | |
|---|---|
| SPBC16E9.12c;pab2 | SPAC2F3.18c;SPAC2F3.17c |
| SPAC1B3.02c | SPBC14F5.13c |
| SPBC16E9.07 | SPAC1348.07;SPBC1348.07 |
| SPBC11C11.11c;SPBC3B8.12 | SPAC11E3.09;pyp3 |
| SPAC2F7.05c;tif5;eif5 | SPAC10F6.11c |
| SPAC8F11.10c;SPACUNK4.10c;SPACUNK4.18;pvg1 | SPAC21E11.03c;pcr1;mts2 |
| SPAC1B3.16c;vht1 | SPBC106.11c |
| SPCC777.10c;ubc12 | SPBPJ4664.03;mfm3 |
| SPCC794.03 | SPAPB1A10.07c |
| SPAC20G8.04c | SPAC6B12.07c |
| SPBC887.15c | SPAPB1E7.10;rpc17 |
| SPBC409.05;skp1;sph1;shp1;psh1 | SPAC17A2.10c |
| SPCC4B3.10c;ipk1 | SPAC2E1P5.03 |
| SPBC21C3.09c | SPAC644.06c;clo3;cdr1;nim1 |
| SPBC3E7.08c;rad13 | SPAC4C5.01 |
| SPBC1683.07;mal1 | SPAC9E9.04 |
| SPAC1296.03c;sxa2 | SPBC12D12.08c;SPBC24C6.01c;ubl1;nedd8;ned8 |
| SPAC222.05c;mss1 | SPAC23A1.03;apt1 |
| SPAC144.05 | SPBC32F12.03c;gpx1 |
| SPAC343.08c;mrp17 | SPBC23G7.10c |
| SPAC6G9.04;mug79 | SPCC338.12 |
| SPAP7G5.03 | SPBC36.02c |
| SPAC17A5.04c;mde10 | SPAC25G10.09c;SPAC27F1.01c |
| SPAPB15E9.03c;Tf2-5 | SPAC343.12;rds1 |
| SPBC725.12 | SPAC1687.16c |
| SPCC965.08c;alr1 | SPCC4G3.10c;rhp4b;rhp42 |
| SPBC28F2.06c;mdm12 | SPBC31A8.01c;SPBC651.13c;cwl1;cwl1B51.12c;rtn1 |
| SPCC16C4.11;pef1 | SPNCRNA.123 |
| SPAC1093.04c | SPAC21B10.02;SPBC21B10.02 |
| SPBC1271.07c | SPBC21H7.06c |
| SPBP8B7.05c | SPBC8E4.01c;SPBP4G3.01 |
| SPCC1919.08c;mrpl33 | SPBP8B7.15c |
| SPAC806.05 | SPBPB2B2.07c |
| SPBC1347.08c | SPAC19G12.10c;pcy1;cpy1 |
| SPAC29E6.01;SPAC30.05;pof11 | NA |
| SPCC320.06 | SPCC553.10 |
| SPCC1020.07 | SPBC947.04 |
| SPAC24H6.12c;uba3 | SPBC2F12.15c |
| SPAP27G11.16 | SPAC18B11.09c |
| SPAC17C9.02c;lys7 | SPBC1105.09;ubcX;ubc15 |
| SPAPYUG7.06;hag1 | SPAC3C7.14c;uhp1;p25;obr1 |
| SPBC1198.01 | SPAC1348.01;SPBC1348.01 |
| SPAC4F10.16c | SPAC3A11.11c |
| SPBC409.04c;mis12 | SPAC4F10.19c |
| SPAC607.06c | SPAC3H1.11 |

SPAC13G6.01c;SPAC5H10.14c;rad8B3F5.10;rad8
SPAC3H1.06c
SPBC16E9.18;SPBC1E8.01
SPAC30D11.10;rad22;rad22A
SPAC31G5.08;ups;ups1
SPCC320.14;SPCC330.15c
SPCC1672.06c;asp1
SPBC12C2.11;SPBC21D10.02
SPCC790.03
SPAC1039.06
SPAC2G11.07c;ptc3
SPCC417.09c
SPAC56F8.07
SPBC25D12.05;trm1
SPCC338.14;ado1
SPCC4G3.19;alp16
SPBC1539.09c;trp1
SPAC1556.03;azr1
SPBC19F8.07;mcs6;mcs2;crk1;mop1
SPBC713.07c
SPCC162.03
SPCC736.08
SPCC2H8.02
SPAC186.03
SPCC13B11.04c;SPCC777.01c
SPAC23H4.10c;thi4
SPAC3A12.06c
SPAC5H10.08c;pan6
SPAC2F3.09;hem1
SPCC18.11c;sdc1
SPBC23G7.14
SPCC10D6.04;SPCC737.07c
SPAPB2B4.02;grx4;grx5
SPAC26H5.06;pot1
SPAC359.03c;SPBC359.03c
SPCC1020.03
SPAPB21E7.07;SPBPB21E7.07;aes1
SPBC2D10.13;est1
SPCC550.02c;cwf5;ecm2
SPAC5H10.10
SPBC887.05c;cwf29
SPAC27D7.04;omt2
SPAC1071.04c
SPCC777.03c
SPAC4G8.07c
SPAC20H4.04
SPBP16F5.08c
SPAC222.11;hem13
SPAC26F1.10c;pyp1
SPAC4F10.20;grx1
SPBC1198.02;dea2
SPAC977.14c
SPAC750.01
SPAC25B8.18
SPAC13A11.03;mug32;mcp7
SPAPB24D3.07c
SPAC24B11.06c;spc1;phh1;sty1
SPCP1E11.07c;cwf18
SPAC922.04
SPBC1861.01c;SPBC56F2.13;cnp3
SPBC23G7.12c;let1;rpt6
SPAP8A3.04c;hsp9;scf1
SPBP4G3.02;pho1
SPBC3E7.06c
SPAC212.09c
SPCC794.12c;mae2
SPAC750.08c
SPBC1861.02;abp2
SPAC16C9.06c

SPAC30C2.07
SPAC24C9.07c;pgs2;bgs2;meu21;gls2
SPAC4A8.08c;vas1
SPBC2G2.17c
SPAC1F12.04c
SPAC21E11.04;ppr1
SPAC23C4.03
SPAC17A5.05c
SPAC24C9.15c;mde9;meu28;spn5
SPCC188.10c;SPCC584.07c
SPAC29B12.08
SPAC15F9.01c
SPBC23E6.08;sat1
SPBC3H7.05c
SPAC1039.10;SPAC922.01;hpm1;mmf
2;hpf1
SPAC144.09c;sfc2;TFIIIA
SPAC19D5.07
SPCC10D6.08;SPCC737.03c
SPBC25B2.08
SPCC965.12
SPAC18B11.03c
SPAC21B10.06c;SPBC21B10.06c
SPBC19C2.06c
SPCC63.03
SPBC418.01c;SPBC887.20c;his4
SPCC63.13
SPAC806.06c
SPAC3F10.10c;map3
SPAC20G4.08;SPAC4F10.01
SPAP27G11.12
SPAC11D3.10
SPBC354.03;swd3
SPAC9E9.09c
SPBC16D10.05;mok13
SPAC27D7.09c
SPBC1D7.05;SPBC1D7.05c;SPBC2F12.
01;byr2BF12.02c;byr2;ste8
SPBC2G2.02
SPAP7G5.04c;lys1
SPBC428.17c
SPCC1183.04c
SPAC19E9.03;SPAC57A10.01;pas1
SPAC13G7.12c
SPCC191.11;inv3;inv1
SPBP22H7.02c;pi029
SPAC806.04c
SPAP27G11.04c
SPBC29A3.14c;trt1

SPCC1020.13c;SPCC14G10.05
SPCC126.14;prp18
SPCC965.06
SPAC2E1P3.04
SPAC9.09
SPBC887.06c;grd19;snx3
SPAC3F10.02c;TRK;trk1;TKH;sptrk
SPBC21C3.15c
SPBC19F8.05
SPBC18B5.10c;SPCC18B5.10c
SPCC1235.02;SPCC320.01c;bio2B20.0
2c;bio2
SPAC4F10.08
SPAC22A12.08c
SPAC222.15;SPAC821.01;meu13
SPBC13E7.06
SPAC6F6.16c
SPAPYUG7.04c;rpb9
SPAC823.07
SPAC1782.09c;flp1;clp1
SPAC227.11c
SPAC1002.07c;ats1
SPCC1682.09c
SPAC24H6.06;sld3
SPAC29A4.14c
SPBC776.14;plh1
SPAC17G6.03
SPBC1773.06c
SPBC28F2.05c
SPAC1399.02
SPBC577.15c
SPAC8C9.10c
SPCC16C4.06c
SPAC2E1P3.05c
SPAC343.18
SPCC63.06
SPCC757.09c;rnc1
SPAC13F5.09;SPAC140.02;gar2
SPCC757.05c
SPAC13A11.04c;ubp28;ubp8
SPAC25B8.03
SPAC27F1.09c;sap155;prp10
SPAC25B8.14;mal2
SPAC17A5.06;ercc3sp
SPAC17A2.02c
SPBC21C3.06
SPCC63.04;mok14
SPNCRNA.09

SPBC16D10.04c;dna2
SPAC11D3.02c
SPBC3B8.01c
SPBC16E9.10c
SPAC6G10.02c;tea3
SPAC1093.03
SPAC144.03;min3;ade2;min10
SPBC1A4.09
SPBC530.07c
SPCC594.05c;spp1
SPCC895.09c;ucp12
SPAP27G11.15;slx1
SPCC1906.04;wtf6;wtf20
SPAC13G7.13c;SPAC6C3.01c;msa1
SPBC12D12.01;SPBC16H5.01c;sta1;sa
d1
SPCC4B3.08
SPAC1A6.05c
SPAC25B8.16
SPAC23G3.02c;sib1
SPBC1198.09;ubc16
SPAC9E9.06c;THRC;thrc
SPBC11G11.02c;end3
SPAC4F10.17
SPAC2F7.06c;pol4;Pol-mu
SPAC19A8.10
SPBC1105.02c;lys4
SPCC895.06
SPCC1529.01;SPCC794.14
SPAC10F6.14c
SPCC74.06;phk2;mak3
SPAC637.11;suv3
SPBP8B7.01c
SPAC1B3.10c
SPCC306.03c;cnd2
SPBC3H7.07c
SPAC3A11.08;Cul-4;pcu4;cul4
SPBC16G5.17
SPAC3A12.02
SPBC4B4.04
SPAC4G9.10;arg3
SPCC4B3.11c
SPAC824.06;tim14
SPAC12B10.05
SPAC27D7.08c
SPCC645.12c
SPCC330.08;gmd3;alg11
SPCC830.03

SPBC660.08
SPAC29E6.02;SPAC30.06;prp3
SPBC13A2.03
SPBC1198.14c;SPBC660.04c;fbp1
SPCC1739.01;SPCC1906.05
SPAC1002.15c;pmc5;med6
SPBC17D1.07c
SPAC1782.01;SPAPYUG7.07
SPBC1778.07
SPBC409.10;ade7
SPCC777.09c;arg1
SPAC26F1.05
SPAC4A8.14;prs1
SPAC1527.01;SPAC23D3.15;mok11
SPBC16C6.01c;SPBC543.11c
NA
SPAC144.02
SPBC215.08c;arg4
SPAC25B8.05
SPCC1840.04;mca1
SPAC4G9.20c
SPCC1450.08c;wtf8;wtf16
SPAC22A12.10
SPBC24E9.14c;SPBC839.14c
SPAC8F11.05c
SPBC725.17c;rrn11
SPBC27B12.03c;pi075
SPBC14F5.01;SPBC1861.10
SPAC824.03c
SPBC119.10
SPAC29A4.07;srb6;med22
SPBC1A4.07c
SPAPB21E7.08;SPBPB21E7.08
SPAC4G8.06c;trm12
SPAC4G9.08c;rpc2
SPBC1306.02;SPBC4.08
SPAC6F12.1;SPAC6F12.10c;ade3;min1
1
SPCC417.02;hos3;hsk3;dad5
SPCC16C4.14c;sfc4
SPAC23H3.12c
SPAC6G10.08;idp1
SPCC1682.08c
SPAC25B8.15c
SPCC550.11
SPAC1B2.05;SPAC3F10.01;mcm5;nda4
BC7.13c;nda4
SPAC22H10.04
SPAC144.06;apl5

SPAP27G11.09c
SPAC926.05c
SPBC27B12.12c;pi066
SPAC1399.01c
SPCC1840.03;pse1;sal3
SPBC3H7.06c;pof9

SPCC24E4.01;SPCC569.08c;ade5;ade8
SPAC13G6.06c
SPAC1002.14;itt1
SPCC613.02
SPBC4B4.06;vps25
SPBC16D10.06
