## Supplementat Table 3 for "Genome-wide chromosomal association of Upf1 is linked to Pol II transcription in *Schizosaccharomyces pombe*"

Table S3 strains_list

| | | |
|---|---|---|
| WT | *h- ade6-M216 leu1-32 ura4-D18* | Current Genetics 1996 30(4):284-93, |
| WT | *h+ ade6-210 arg3D his3D leu1-32 ura4DS/E* | |
| upf1-HA-Cdc25 | *h? cdc25-22 ade6-704 leu1-32 ura4-D18 upf1-3HA::kanMX6* | This study |
| upf1:flag | h+ ade6-210 arg3D his3D leu1-32 ura4DS/E upf1:5flag::hphMX6 | This study |
| JY741/FLAG-Rpb3 | h- flag-rpb3 ade6-M216 ura4-D18 leu1 | from Japanese National BioResource Project--Yeast |
| upf1Δ FLAG-Rpb3 | h- flag-rpb3 ade6-M216 ura4-D18 leu1 upf1::hphMX6 | Mol Cell Biol. 2006 26(17): 6347–6356 |
| FLAG-Rpb3-upf1-HA | *flag-rpb3:Upf1-3HA, ade6-M216 ura-D18 leu h-* | This Lab |
| FLAG-Rpb3:upf1Δ | *h-flag-rpb3 ade6-M216 ura4-D18 leu 1 Upf1::hphMX6* | This Lab |
