## Supplementat Table 4 for "Genome-wide chromosomal association of Upf1 is linked to Pol II transcription in *Schizosaccharomyces pombe*"

Table S4 Primers table

| | Primer name | sequence | Comments |
|---|---|---|---|
| *Pma1* | Pma1-p1-F | GTCTTCGTGATTGGGTCGAT | |
| | Pma1-p1-R | GGGGTCACCATAGTGCTTGT | |
| | Pma1-p2-F | ATCCCGTTTCCAAGAAGGTT | |
| | Pma1-p2-R | GAGGATCGGAACAAGGCATA | |
| | Pma1-p3-F | GTCTTTCCACCGTCATTGGT | |
| | Pma1-p3-R | ACGGAGAACGGCAACAATAG | |
| | Pma1-p4-F | GAAACTATAGGTTAATGGAAG | |
| | Pma1-p4-R | GTTTCCTGCCGGCTTGTC | |
| *Act1* | Act1-t1-F | GCTCAATGTTATCCGTTTCCG | |
| | Act1-t1-R | GTAGTTGGTAAACGGTAAGTTATAACAC | |
| | Act1-t2-F | GGAAGAAGAAATCGCAGCGT | |
| | Act1-t2-R | CATATCATCCCAGTTGTTGACAATAC | |
| | Act1-t3-F | GAAATGTGATGTTGATATTCGTAAAG | |
| | Act1-t3-R | GCTCTCATCATACTCTTGCTTGG | |
| | Intergenic-RT-F | AGAGGCACATAGTAGGGGAACT | |
| | Intergenic-RT-R | TCCCATCTCCCACTGTTAATTGA | |
| | tf2 | ACACCAACACAAACCCAAGCGA | from 2035-2056 bp 0f ORF of tf2-1 |
| | tf2 | ACGGCTCCTACAGCGACATCT | from 2165-2145 bp 0f ORF of tf2-1 |
| | tRNAMet#1F | AAAAGAAAACGGTCAGGGAGG | Pebernard, 2008; SPBTRNAMET.05 |
| | tRNAMet#1R | GAGCCTCACCAGGAGCATTATAG | Pebernard, 2008; SPBTRNAMET.05 |
| | tRNAAla#1R | CCTGCAAACGTATGTTACGTAAGG | Pebernard, 2008 |
| | tRNAAla#1F | TCCAATTATTAAGTGAATGCTCTCG | Pebernard, 2008 |
| | tRNAAsnF | GGTCGGGTAGCATAGTTGGTT | SPBTRNAASN.01 |
| | tRNAAsnR | AGAAAACGGTCAGGGAGGGA | SPBTRNAASN.01 |
| | tel | TCA AAG TTG GCG ACG TTG CTG ATG | detect telomeric region; Rozenzhak S, 2010 |
| | tel | AAG CAA TGT GTG GAG CAA CAG TGG | detect telomeric region; Rozenzhak S, 2011 |
| | Upf1-F | TGTTACAATTATTTACACTTTGCAAATTGACGGC | For HA tagging of Upf1 gene |
| | Upf1-R | ATATCAACAAATAAAAGATATGTTGGCATTCGTA | For HA tagging of Upf1 gene |
| *Intergenic* | Intergenic_R | GCGAAACCAGTATGGACGAT | |
| | Inttergenic_F | AACGGGCAAATGTAAAGACG | |
| *SPBC609.01* | SPBC609.01-1F | AAGGGATGCAGACAACTCCA | |
| | SPBC609.01-1R | GGTTTCAAAGGCGTCAGGAA | |
| | SPBC609.01-2F | CACACTGAGCAACTTCTACCG | |
| | SPBC609.01-2R | TCAACAGCCAAAACAAAAGCC | |
| *met26* | met26-1F | CGTGAGTTGAAGAAGGCCAC | |
| | met26-1R | CAACACCTTGGGCCTTTTGA | |
| | met26-2F | AAGTTGCTTCTTCCTCCAGC | |
| | met26-2R | GCCGTCATTAGCAGCCTTAG | |
| | met26-3F | GGATGCTGACGTTGTTTCCA | |
| | met26-3R | AACAGGAGGAACACGAGGAG | |
| *ght5* | ght5-1F | TCATGTTGGTCTTCGTGTCC | |
| | ght5-1R | CGCGCCGAAGAATACGAATA | |
| | ght5-2F | CGCTACTTGGCCCGAAATTT | |
| | ght5-2R | CCCTCAAACACCTCGAAACC | |
| | ght5-3F | ATCTCTGGTGCTAAGCCCTG | |
| | ght5-3R | ACGTTTTCAAGATGAGCGGC | |
| *mug106* | mug106-1F | TCCTTTTCCACCTGCAAACG | |
| | mug106-1R | TTCATGTGCTGGATTGGGTT | |
| | mug106-2F | TCTCCCATTTTGCACGAGGA | |
| | mug106-2R | TCGCCTACTAACACCGGTAC | |
| *tpi1* | tpi1-1F | CGTTGGTGATGTCGAAACTGT | |
| | tpi1-1R | GTAGGCACCGTTCTTCTTGTC | |
| | tpi1-2F | CTGTCTGGGCCATTGGTACT | |
| | tpi1-2R | GTAGATGACACGGAGACCCT | |
| *ada2* | ada2-1F | TGTTCCAAAAGGCAACATGG | |
| | ada2-1R | TCTTACTGCCGAAGCGAATG | |
| | ada2-2F | CACCATCCGTCTCATCCGTA | |
| | ada2-2R | CTGCAATGTCAGCCCAGTTT | |
| | ada2-3F | CAGCGACTCCCACAATGTTT | |
| | ada2-3R | TGGAAGTCAGTGAAGCCGAT | |
